# Integrating nekton communities, salt pool dynamics, and fish foraging patterns to assess salt marsh restoration success in Maritime Canada

**DOI:** 10.1101/2025.10.31.685895

**Authors:** K.C. Endresz, N.R. McLellan, M.A. Barbeau

## Abstract

Salt marsh restoration aims to recover areas and ecological functions that have been impacted by human activities. We evaluated different structural and functional metrics to assess the response of nekton to salt marsh restoration in two distinct case studies in Maritime Canada in 2022–2023. We compared the visiting nekton communities in intertidal creeks and on marsh platforms, the abiotic conditions and small-bodied faunal communities in salt pools, and the foraging patterns of two fishes between reference and restoring marshes. In both case studies, similar nekton species were observed between reference and restoring marshes; however, nekton communities differed by site type (14-52% of the variation). Pools in restoring marshes were providing basic nekton functions but had yet to offer the enhanced habitat support as those in reference marshes. Gut fullness was similar in tomcods (*Microgadus tomcod*) and mummichogs (*Fundulus heteroclitus*) between site types suggesting that restoring marshes were fulfilling foraging function for these species despite site type differences (11-44% of the variation) in prey composition being detected. Although the nekton community structure and functions in restoring marshes did not mirror those in references marshes, our findings highlight the value of restoring salt marshes in terms of nekton habitat. This study also demonstrates the advantages of implementing a multifaceted approach to understand the response of biotic communities to salt marsh restoration.

## 1. INTRODUCTION

Salt marshes are prominent intertidal ecosystems, characterized by dense grassy vegetation and high levels of productivity (Bertness 2007, Mcowen et al. 2017). When tidally inundated, diverse nekton communities take advantage of these valuable and intermittently available habitats (Rozas 1995). Intertidal creeks facilitate access to the resource rich marsh platform, and salt pools offer aquatic refuge in between periods of tidal flooding (Cattrijsse et al. 1994, Rountree & Able 2007, Mace et al. 2019). Together, these subhabitats provide foraging, nursery, overwintering, and spawning grounds to numerous nekton species (Smith & Able 1994, Minello et al. 2003, Able et al. 2012). Despite the recognition of salt marshes’ important ecological role to a diversity of nekton, their loss is considerable worldwide (Gedan et al. 2009, Mcowen et al. 2017). Maritime Canada is one region that has experienced extensive losses of this valuable ecosystem due to human activities, with < 40% of original salt marsh area remaining (Thomas 1983, Hanson & Calkins 1996).

Conversion of salt marsh to terrestrial land in Maritime Canada began in the late 1600s and early 1700s when areas were diked by Acadian settlers and subsequently other colonists (Bleakney 2004, Rudin 2021). In the 1970s–1990s, areas of salt marsh (often in the footprint of former dikeland) were diked and transformed into freshwater wetland impoundments to create habitat for waterfowl species (Loder et al. 2017). The dynamic coastal environment has over time taken a toll on the dikes (e.g., Boone et al. 2017). Maintenance of these dikes is costly and logistically challenging especially in the face of heightening climate change impacts (Sherren et al. 2021). Consequently, more interest and efforts are being directed to restoring salt marshes back to their former natural state. Furthermore, wetlands, including salt marshes, in Maritime Canada are now protected under various federal and provincial legislations that aim to conserve these ecosystems and ensure any further loss or impact is compensated for (Government of Canada 1985, Environment Canada 1991, Government of New Brunswick 2002, Government of Nova Scotia 2011). Unavoidable destruction or degradation of wetland habitats requires offsetting, leading to wetland restoration opportunities including salt marshes, which are often prioritized due to historic loss and the functions they provide. Fish habitat impacts, as determined through legislation (Government of Canada 1985), can also be offset through salt marsh restoration, leading to additional opportunities (Boone et al. 2017, Ducks Unlimited Canada pers. comm.). Given the ecological importance of salt marshes to nekton, and as restoration opportunities arise, it is essential that we understand the degree of habitat support provided by recovering areas.

The overarching goal of ecosystem restoration is to return lost or degraded habitats back to their approximate former natural state resulting in positive ecological outcomes (Simenstad & Cordell 2000). Complexities arise when attempting to determine whether a restoration can be deemed successful (Billah et al. 2022); therefore, monitoring structural and functional responses requires appropriate metrics (Thom et al. 2011; Krueger et al. 2017). For salt marshes and nekton support, comparing nekton communities, abiotic and biotic variables of an important subhabitat, and usage patterns by key species between reference (minimally impacted) and restoring sites enable measurement of recovery trajectories and effectiveness (Weinstein et al. 2019). Individually, each metric gives valuable insights depending on monitoring goals; collectively, they provide a comprehensive understanding of how nekton respond to salt marsh restoration.

Selecting focal fish species for more in-depth study of nekton response to salt marsh restoration provides a direct measure of habitat support. Atlantic tomcods (*Microgadus tomcod,* hereafter tomcod) are a transient species that use salt marshes along the northeastern coast of North America as nursery and feeding grounds (Stewart & Auster 1987). Tomcods hold unique cultural value for Mi’kmaq people as the timing of their anadromous spawning run once provided a critical food source during winter months (Kavanaugh et al. 2004). Mummichogs (*Fundulus heteroclitus*) are a ubiquitous resident fish in salt marshes along the Atlantic coast of North America. They are integral in salt marsh trophic dynamics because they link terrestrial and nearshore aquatic food webs and are a main food source for birds and larger-bodied transient fishes (Kneib 1986; Lesser et al. 2021). Knowledge of how target fish species respond to salt marsh restoration helps develop or modify strategies to improve outcomes.

We evaluated the recovery status, in terms of nekton response and support, by integrating structural and functional metrics in two distinct salt marsh restoration projects (i.e., case studies; Aulac, New Brunswick and Wallace Bay, Nova Scotia) in Maritime Canada. More specifically, our objectives were to compare within each case study (i) the nekton community in intertidal creeks and on marsh platforms, (ii) the environmental conditions and fauna in salt marsh pools, and (iii) the gut fullness and contents of tomcods and mummichogs between reference and restoring marshes. To address these objectives, we conducted extensive sampling over two years (May to July 2022-2023) and retained tomcods and mummichogs for gut content analysis.

Through complementary analytic methods, we examined univariate indicators (e.g., species richness, gut fullness) and assemblage structure (nekton communities, potential prey communities, pool abiotic conditions, pool vegetation, and gut contents) quantifying spatiotemporal scales of variation and dissimilarity, and for assemblage composition, contribution of taxa or response variables to multivariate dissimilarity, between reference and restoring marshes. The findings from our Maritime Canadian case studies provide insights into the role restoring salt marshes play in providing habitat to nekton.

## 2. MATERIALS & METHODS

Sampling occurred during periods of spring tides (higher high tides) in May-July 2022-2023 (Table S1). Differences in the tidal regime, age, and former state of the restorations resulted in each project being treated as a case study to focus comparisons on reference and restoring marshes. Our reference sites were minimally impaired, unrestored natural salt marsh and were representative of the expected ecological conditions of other salt marshes in the region (Virgin et al. 2020, Endresz et al. 2026).

### 2.1. Overview of restoration projects

The salt marsh restoration in Aulac, New Brunswick (45°51.07’N, 64°17.64’W) is in the upper Bay of Fundy and experiences megatides (amplitude > 8 m; Canadian Hydrographic Service 2022) (Fig. 1). The Aulac restoration began in October 2010 when three sections of dike that protected pastureland were breached to revert the areas back to salt marsh (Boone et al. 2017, Virgin et al. 2020). The salt marshes in Aulac were first diked in ∼1750 and thus have not been fully and continually exposed to the natural tidal regime for over 250 years (Maritime Marshland Rehabilitation Administration 1953, Nova Scotia Department of Agriculture and Marketing 1987). We treated the two pastureland parcels as one restoring site for our project due to their relatively small size (together covering ∼20 ha) and proximity to each other. At the time of sampling, the Aulac project was in its twelfth and thirteenth year of restoration. The reference marsh (∼18 ha) is located adjacent to the restoring site and supports a typical salt marsh plant community for the upper Bay of Fundy, with narrow low marsh zones of saltwater cordgrass *Spartina alterniflora* (syn. *Sporobolus alterniflorus*; Peterson et al. 2014, Bortolus et al. 2019) at the seaward edge, and surrounding creeks and pools, and an extensive high marsh zone dominated by salt marsh hay *S. patens* (syn. *Sporobolus pumilus*) (Virgin et al. 2020).

**Fig. 1.**
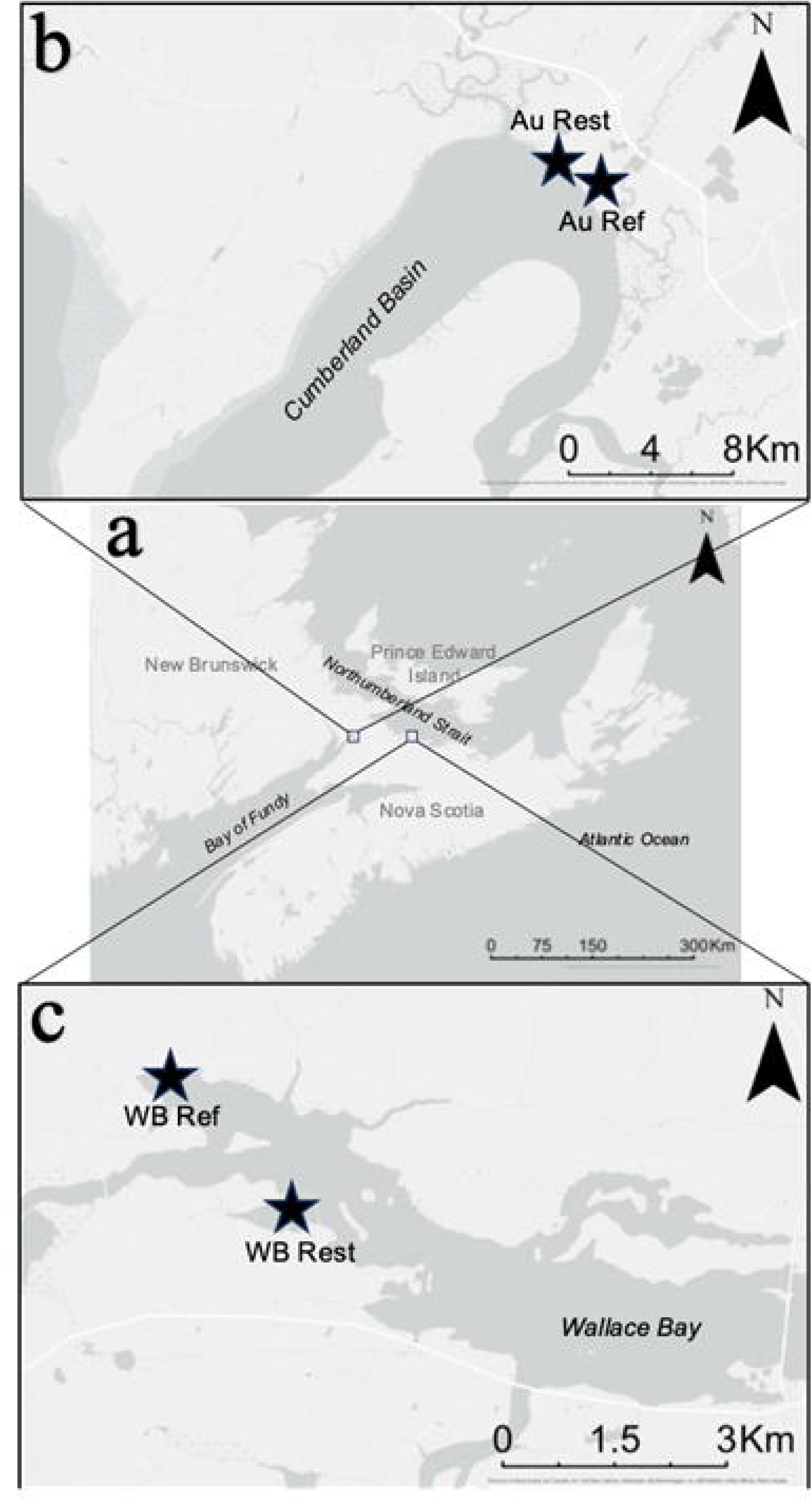
a) Map of Maritime Canada indicating the study sites in Aulac (Au), New Brunswick and Wallace Bay (WB), Nova Scotia (rectangles indicate the study area), and closer view of the reference (Ref) and restoring (Rest) salt marshes sampled in b) Aulac and c) Wallace Bay in 2022–2023. Base maps from ArcGIS Pro, 10 April 2025.

The salt marsh restoration in Wallace Bay, Nova Scotia (45°49.65’N, 63°32.63’W) is on the Northumberland Strait and experiences microtides (amplitude usually < 2 m; Canadian Hydrographic Service 2022) (Fig. 1). The restoring site (∼20 ha) in Wallace Bay was a freshwater wetland impoundment, created by Ducks Unlimited Canada and the Canadian Wildlife Service in the 1970s (Environment and Climate Change Canada 2018). The Wallace Bay restoring site had been diked since the 1700s and used as pastureland and hayland; however, this agricultural land had long been abandoned before it was converted to a freshwater wetland impoundment. Two sections of the dike were breached along with the removal of a water control structure in August 2020 (Lawrence & Van Wychen 2021). At the time of sampling, the Wallace Bay project was in its second and third years of restoration. The reference site (∼30 ha) is connected to an extensive network of salt marshes located ∼1 km away from the restoring site. It has similar areal extents of low marsh zone with *S. alterniflora* and high marsh zone dominated by *S. patens* (Lawrence & Van Wychen 2021; Dickinson 2024).

### 2.2. Sampling intertidal creek and marsh platform nekton communities

To examine the visiting nekton community, we captured nekton in creeks and on the marsh platform in reference and restoring sites using methodology described in Endresz et al. (2026). Briefly, we deployed two fyke nets (68.5 cm × 99 cm frame, 3-mm mesh, two 5-m-long wings set 7.45 m apart at the leading edge) each at a creek mouth for an overnight high tide, as well as conducted three replicate seine net hauls (10 m × 1.2 m, with a 0.6 m × 0.6 m × 1.2 m pocket, 3-mm mesh, 30-m long transects) on the marsh platform during a daytime high tide. We counted and identified captured nekton to species, and measured body size (total length [TL] for fish and shrimps, and carapace width for crabs) of the first 30 haphazardly selected individuals of each species (Table S2.1). Juvenile and adult fish were differentiated based on size classification in the literature (Table S2.2). All nekton were released back into the water after processing except for up to ten tomcods (size range 10□20 cm TL) and six mummichogs (size range 5□11 cm TL). These fish were retained from each site during each round, from fyke nets fishing the ebb tide and from seine hauls at high tide for gut content analysis (see below). For each sampling round, we recorded surface water conditions (pH, salinity, temperature, and dissolved oxygen concentration) of the surrounding bay at high tide by taking measurements at four locations in the main salt marsh channel or in the bay using a YSI Professional Plus meter.

### 2.3. Sampling pool environmental conditions and small-bodied fauna

To examine salt pool quality (including water conditions and potential food sources in terms of invertebrate communities), potential nursery habitat (by sampling very small juvenile fish), and use by nekton, we haphazardly selected four pools based on predefined criteria (surface area 32□120 m^2^ and water depth >20 cm) in each site. In reference sites and the Wallace Bay restoring site, these were salt pools (i.e., depressions on the marsh platform that retain water at low tide). In the Aulac restoring site, true salt pools were not present and so creek pools (i.e., depressions within the intertidal creek network that retain water at low tide) were selected instead (Figs. S3.1 and S3.2). We recognize the inherent difference between these two subhabitats and account for this in our interpretations of the effect of site type on pool abiotic conditions and fauna in Aulac.

In each pool, a minnow trap (dimensions: 42 cm long, 23 cm maximum diameter, 2.2 cm diameter entrance holes, and 0.7 cm x 0.7 cm mesh size) baited with two saltine crackers was deployed at low tide in the evening, just below the surface of the water, to sample nekton over a night high tide. As well, an invertebrate activity trap (2 L plastic pop bottle with the top cut off, mesh (0.3 cm by 0.4 cm) secured over the spout, and the top placed inverted into the bottle and secured with tape; Fig. S3.3) was paired with the minnow trap, to sample aquatic invertebrates and small juvenile fish. Upon trap deployment, we measured water conditions (depth, pH, salinity, temperature, and dissolved oxygen concentration) with a meter stick and a YSI Professional Plus meter. We measured sediment penetrability at the bottom of each pool using a handmade penetrometer (a guiding ABS plastic pipe with a copper rod (852.8 g, 1.39 m long, 16 mm diameter) released 15 cm above the sediment surface). We estimated percent cover of ditch grass (*Ruppia maritima*) and green macroalgae in each pool. Traps were retrieved the ensuing early morning and all fauna captured were identified to the lowest taxonomic level possible (Table 1) and counted before being released.

**Table 1.**
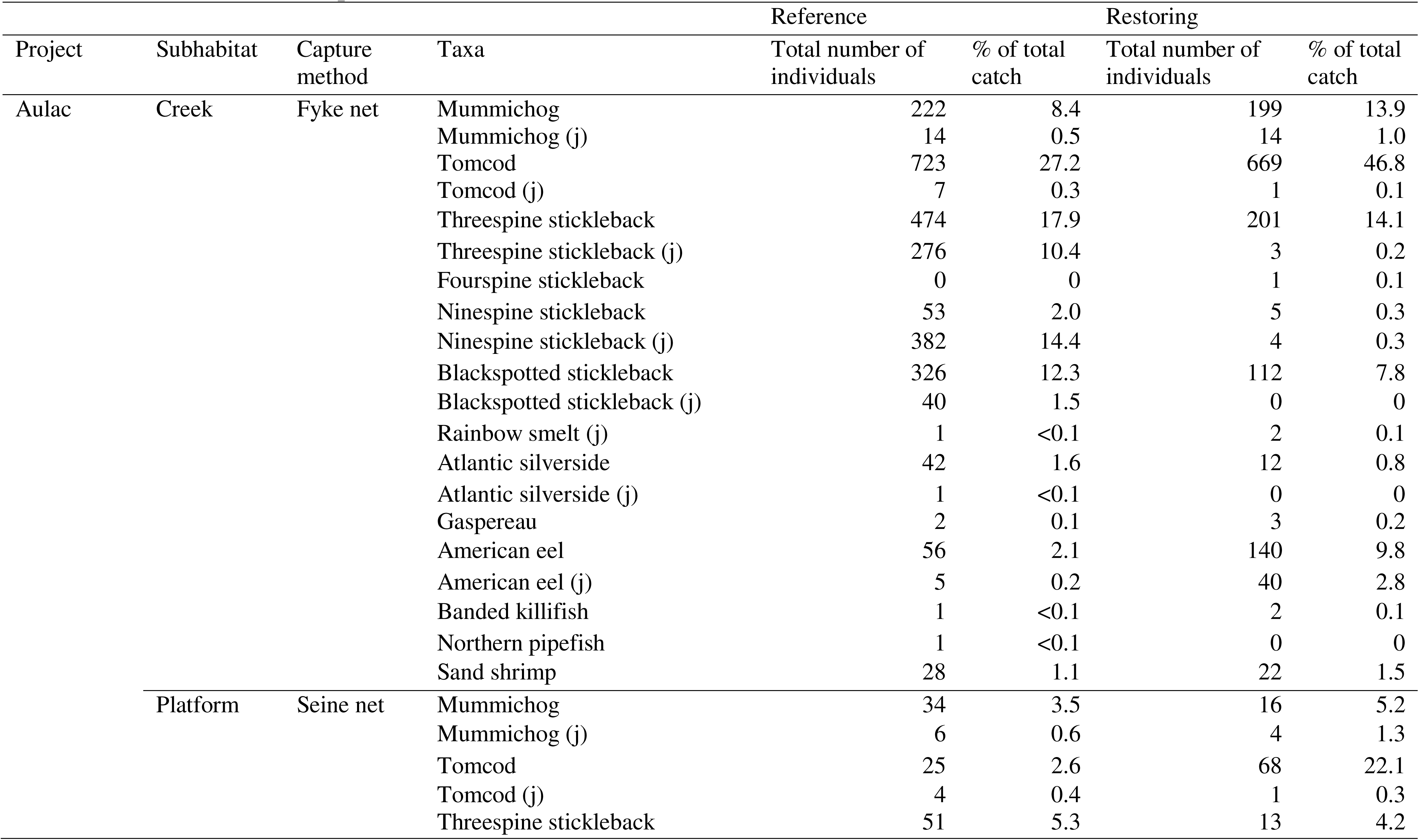
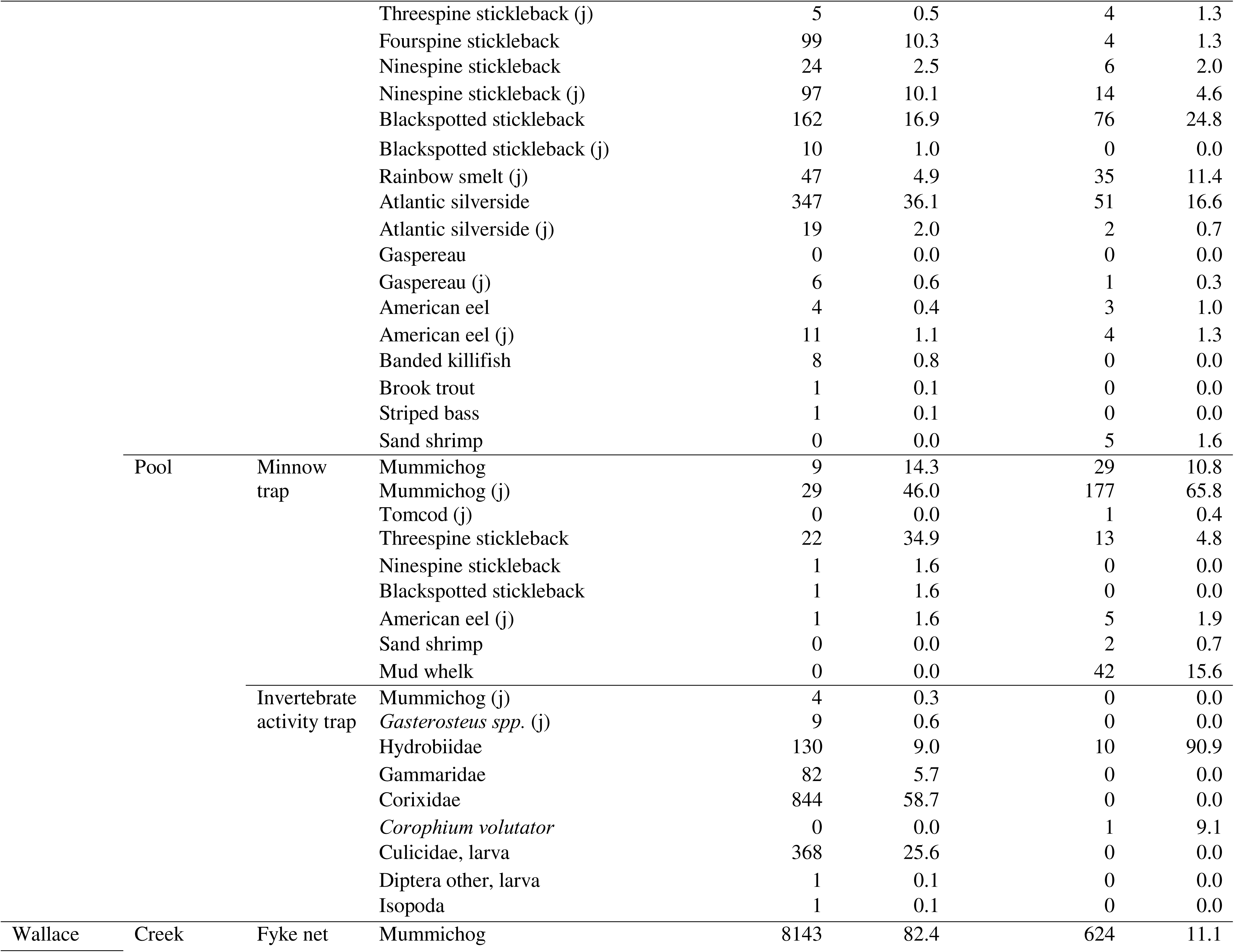
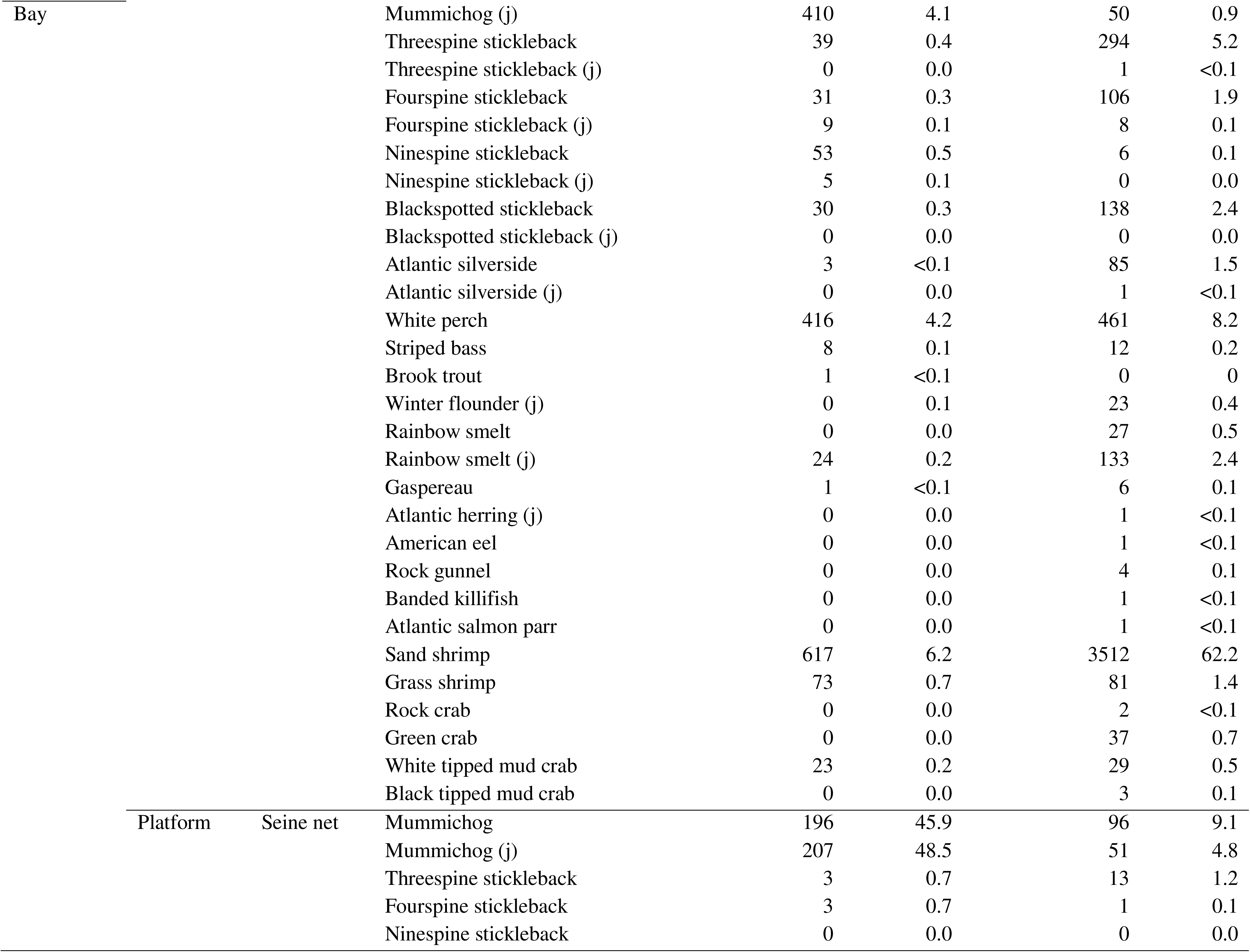
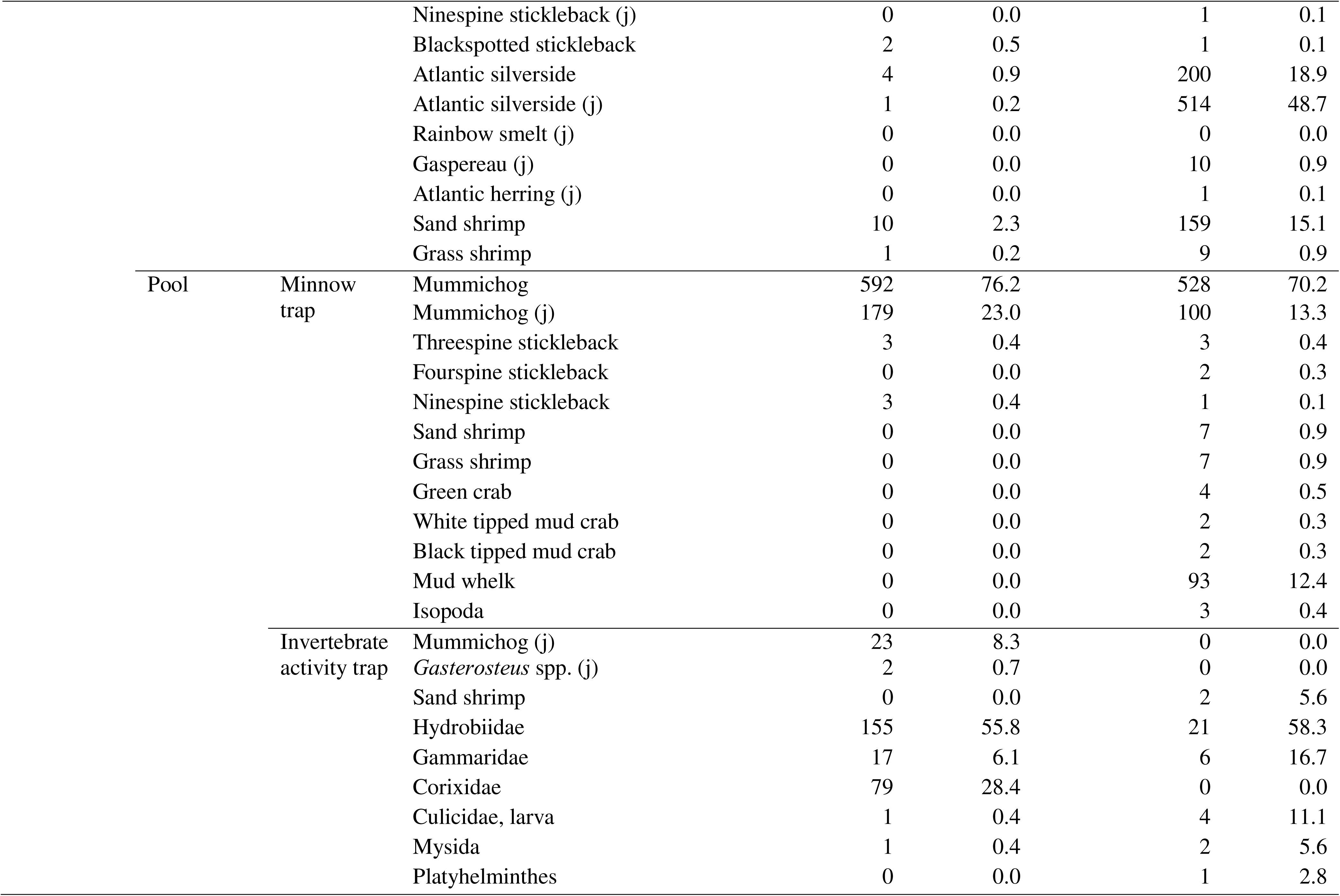
Fish and invertebrate taxa, abundance, and percentage of total catch from fyke nets set in intertidal creeks, seine hauls on the marsh platform, and minnow and invertebrate activity traps deployed in salt pools in reference and restoring salt marshes in Aulac and Wallace Bay in May–July in 2022–2023. Data pooled over all sampling rounds (see Table S1). A “j” following a fish species name indicates juveniles and those without are adults (see Table S2.1). Invertebrates were not differentiated by life stage. Gaspereau are alewives (*Alosa aestivalis*) and blueback herring (*A. pseudoharengus*). *Gasterosteus* spp. in invertebrate activity traps are threespine (*G. aculeatus*) and/or blackspotted (*G. wheatlandi*) sticklebacks.

#### 2.3.1. Gut content analyses

Retained fish from fyke nets facing ebb tides (to reduce issues relating to gut evacuation rates and passive capture methods; Berg 1979) and seine hauls (Table S4.1) were euthanized (clove oil and ethanol solution, University of New Brunswick Animal Care Protocols 22001 and 23001) and then frozen until further processing. Prior to dissections, each fish was thawed, measured for total length (nearest 0.1 cm), and weighed (nearest 0.01 gram). We excised the stomach of each fish, weighed it full and then again when all contents were removed; the difference gave the wet weight of gut contents. While tomcods have a true stomach, mummichogs do not, requiring cuts at the esophagus and the second 180° turn of the digestive tract (James-Pirri et al. 2001). We did not retain the posterior section of mummichog guts because contents are less intact and more difficult to recognize than those in the first sections (Berg 1979). We calculated a standardized gut fullness index as the ratio of the gut content wet weight to the intact wet weight of the fish (Berg 1979). We identified gut items to the lowest taxon possible (see Table S4.2 for faunal prey) and enumerated under a dissecting microscope.

### 2.4. Statistical analysis

For each of Aulac and Wallace Bay, we used PRIMER with PERMANOVA (Permutation Multivariate Analysis of Variance) add-on (version 6; Anderson et al. 2008; Clarke and Warwick 2001) to evaluate differences between reference and restoring sites for: nekton communities in creeks and on platforms; abiotic conditions, aquatic vegetative community, and small-bodied faunal communities in pools; and fish prey assemblages. We constructed resemblance matrices using Bray-Curtis similarity for non-transformed gut fullness indices and vegetation percent cover, 4^th^ root transformed faunal densities, presence-absence for prey assemblages (as well as 4^th^-root transformed prey assemblages; in Electronic Supplement 4), and Euclidean distance for normalized abiotic variables. We transformed data to better assess contributions of all taxa to community patterns (4^th^ root), to include vegetal items (presence-absence) in gut analyses, or to account for different units (normalized) for abiotic analyses. A dummy variable (value = 0.01) was added to deal with zero densities for organismal community datasets (Clarke & Gorley 2015). Site type (Reference, Restoring), Year (2022 and 2023), and Month (May, June, July) were fixed factors. For nekton density in creeks and on platforms, Net (2 levels per Site type) and Haul (3 levels per Site type), respectively, were the base unit of replication. For creeks, deployed nets sampled a different mass of water each round and so were treated as independent. Since the same pools were repeatedly sampled among rounds within a year, the random factor Pool was orthogonal to Month and nested in Year and Site type. When the interaction Site type*Year*Month was significant, we analyzed Site type patterns by year.

To complement nekton community multivariate analyses, we conducted univariate analyses on species richness, diversity (Inverse Simpson Index), evenness (Simpson Evenness), and total nekton abundance (Lande 1996; Bittinger 2020) separately for creek, platform, and pool communities, using the same linear model described above. Note that PERMANOVA is appropriate for both univariate and multivariate analyses (Anderson et al. 2008). For invertebrate activity traps in Aulac, they were mostly empty in the restoring site (Table 1); therefore, no univariate analyses were conducted. In Wallace Bay, invertebrate activity traps in restoring pools were empty in June 2022 and July 2023, and so those dates were omitted from analysis.

For PERMANOVA analyses of gut fullness indices and prey assemblage, we recoded Year and Month into year-month combinations termed Round, because we did not always have fish for all sampled months in each year (see Table S4.1). Site type and Round (up to 6 levels of year-month combinations) were fixed factors. Fish individuals, the base unit of replication, were treated as independent because they had time to disperse throughout each site before being captured. If a significant interaction Site type*Round occurred, we conducted pairwise comparisons between Site type levels for each round. Given that fish size can influence consumed prey, we conducted PERMANOVAs on fish length to ensure there was no confounding effect of fish size on gut fullness or prey assemblage results (Tables S4.3, S4.4, and S4.6). Based on the number of fish retained (Table S4.1), we adjusted the number of rounds in each analysis to exclude those where fish were missing from one or both site types. No tomcods were captured in Wallace Bay and very few were caught on the platform in Aulac; therefore, we only analyzed creek tomcods from Aulac.

For all analyses, we used an α = 0.05 significance level. To complement PERMANOVAs, we conducted Permutational Dispersion (PERMDISP) tests on significant fixed effects to assess if detected patterns were due to differences in the means (centroids) and/or variances (dispersion) of the response variables (Anderson 2006). Components of variation were calculated for both fixed and random effects to examine the relative importance of our various spatial and temporal scales. Similarity Percentage (SIMPER) tests were conducted to assess the taxon, prey item, or abiotic variable contributing most as well as contributing consistently (based on the average dissimilarity divided by the standard deviation of dissimilarities) to significant site type patterns.

## 3. RESULTS

### 3.1. The visiting nekton community in Aulac

Over all sampling rounds in Aulac, fyke nets in reference creeks caught 2,654 individuals and 13 species of nekton, while those in restoring creeks captured 1,430 individuals and 12 species (Table 1). Seine hauls on reference platforms caught 961 individuals and 13 species, while those on restoring platforms captured 307 individuals and 11 species. Reference creeks were dominated by tomcods and threespine sticklebacks (*Gasterosteus aculeatus*), while reference platforms were dominated by Atlantic silversides (*Menidia menidia*). Tomcods accounted for almost half of the nekton individuals in restoring creeks and were most abundant on restoring platforms alongside Atlantic silversides and blackspotted sticklebacks (*G. wheatlandi*). Below, we also assessed temporal patterns.

#### 3.1.1. Creek communities in Aulac

Species richness (grand mean: ∼6.2), diversity (∼2.6), evenness (∼0.18, indicating unequal species distribution), and total number of nekton individuals did not differ significantly between reference and restoring creeks in Aulac (Tables 1, S5.1 and S5.2). However, creek nekton communities differed yearly by site type (significant Site type*Year), as well as among months (PERMANOVA, Table 2, Figs. 2 and 3a). If we do not indicate a life stage before a fish species name in the following sections, we are referring to the adult form. Despite small yearly differences, species contributing most to the overall site type community dissimilarity (44%) were juvenile and adult ninespine sticklebacks (*Pungitius pungitius*), threespine sticklebacks, blackspotted sticklebacks, and Atlantic silversides being more abundant in reference creeks, while juvenile and adult American eels (*Anguilla rostrata*), mummichogs, tomcods, and sand shrimps (*Crangon septemspinosa*) being more common in restoring creeks (SIMPER, Table S6.1, Fig. 2).

**Fig. 2.**
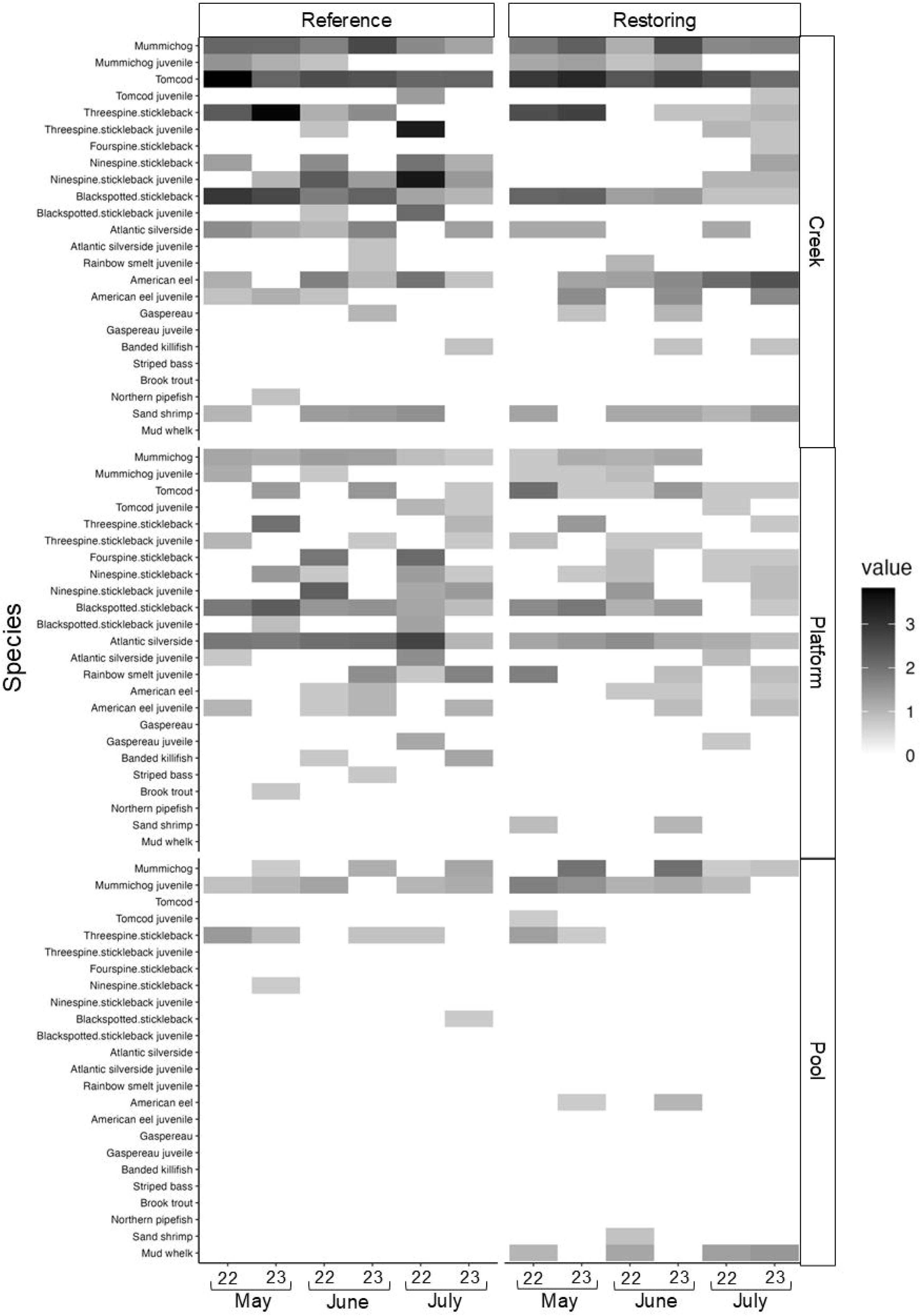
Heatmap showing average densities of nekton taxa (per net or trap; 4^th^ root transformed data) captured in three subhabitats in the Aulac reference and restoring salt marshes for three months (May–July) in 2022–2023. Averaged over 2 fyke nets (creeks), 3 seine hauls (platform), or 4 minnow traps (salt pools) per sampling round per site type (*n* = 2–4). A “J” following a fish species name indicates juveniles and those without are adults. Invertebrates were not divided by life stage. Gaspereau are alewives (*Alosa aestivalis*) and blueback herring (*A. pseudoharengus*).

**Fig. 3.**
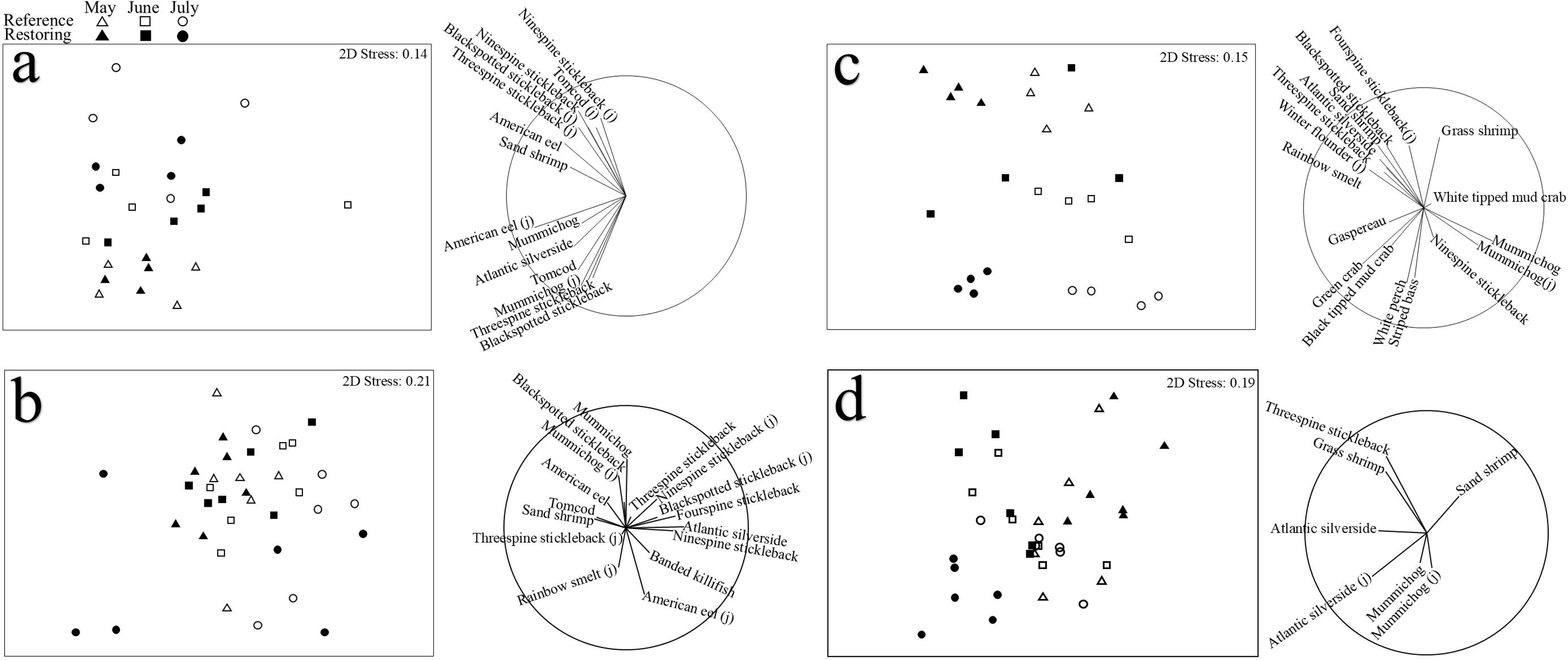
nMDS graphs for (a,c) captured nekton communities in creeks and on (b,d) platforms in reference and restoring salt marshes in (a,b) Aulac and (c,d) Wallace Bay for 3 months (May–July) in 2022–2023. A symbol represents community composition (4^th^ root transformed) averaged per Year-Month-Site type combinations. Juvenile life stage of some fish species indicated by “(j)”, and fish species without are adults. Gaspereau are alewives (*Alosa aestivalis*) and blueback herring (*A. pseudoharengus*). Adjacent vector diagrams represent Pearson correlations (r) between a taxon and nMDS axes; a given taxon vector shows direction of increased density across the corresponding nMDS graph. Circle surrounding vectors indicates maximum vector length (r=1 if parallel to one of the nMDS axes); only correlations with r > 0.2 with at least one nMDS axis shown. Stress values were ∼0.2 or less, indicating that the 2-dimensional representations were adequate for the multi-dimensional situations (Clarke & Warwick 2001).

**Table 2.**
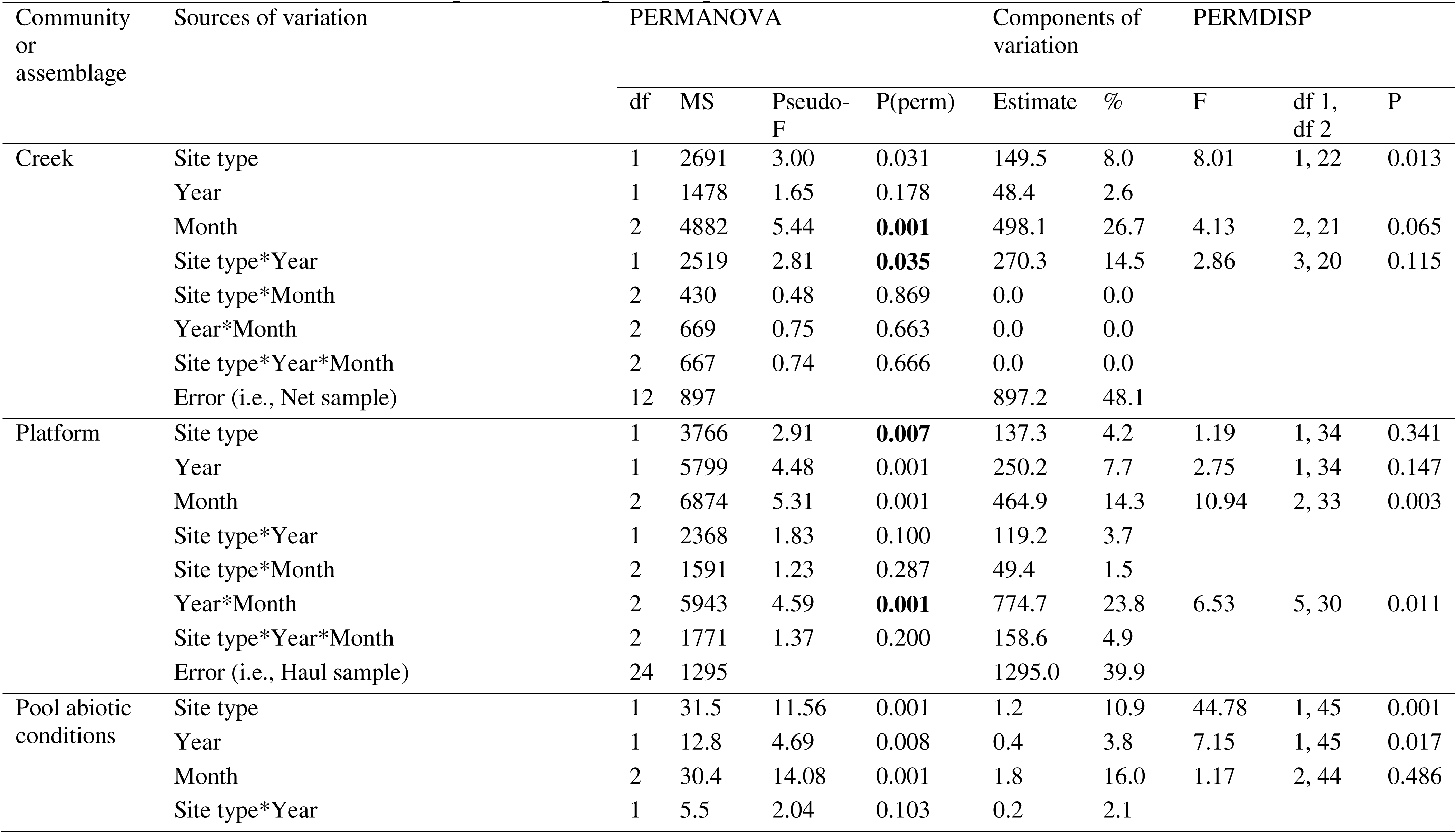
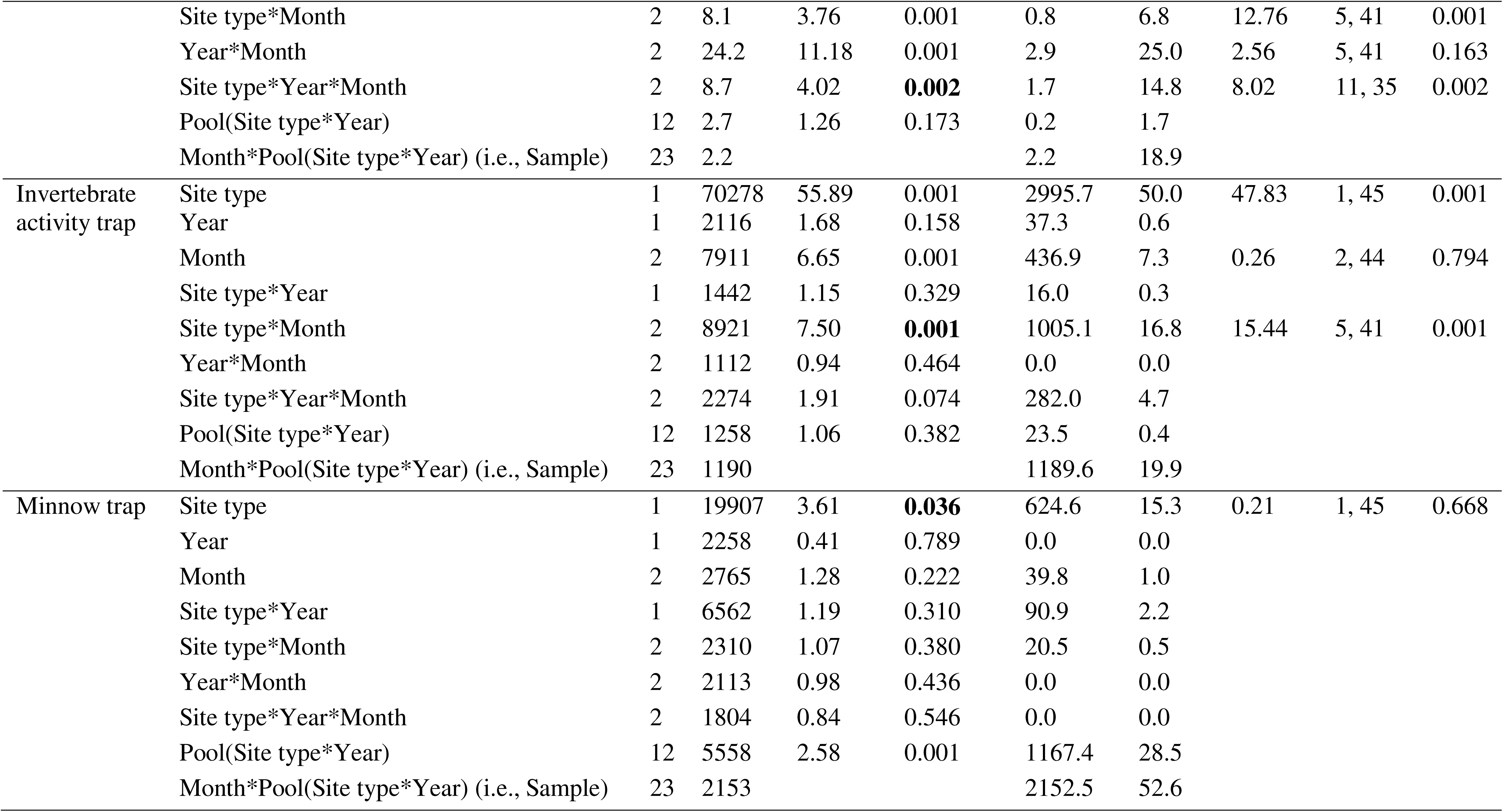
PERMANOVA results comparing fish and invertebrate communities in Aulac reference and restoring salt marshes in May–July 2022–2023; analyses done separately for creek, marsh platform, and salt pool subhabitats. See Table S1 for sampling dates, Table 1 for taxa included in each biotic community, and Table S8.1 for abiotic variables measured in salt pools. Site type, Year and Month are fixed factors, and Pool is a random factor. Taxa densities 4th root transformed and abiotic variables normalized prior to analysis. P-values bolded for significant and interpretable fixed effects; 994–999 unique permutations. PERMDISP tests conducted for significant fixed effects to assess amount of multivariate dispersion among groups; df1 and df2 represent the numerator and denominator degrees of freedom for the F-ratio, respectively. Analysis of components of variation conducted for both fixed and random effects to evaluate relative importance of spatiotemporal scales.

| Community or assemblage | Sources of variation | PERMANOVA |  |  |  | Components of variation |  | PERMDISP |  |  |
| --- | --- | --- | --- | --- | --- | --- | --- | --- | --- | --- |
|  |  | df | MS | Pseudo-F | P(perm) | Estimate | % | F | df 1, df 2 | P |
| Creek | Site type | 1 | 2691 | 3.00 | 0.031 | 149.5 | 8.0 | 8.01 | 1, 22 | 0.013 |
|  | Year | 1 | 1478 | 1.65 | 0.178 | 48.4 | 2.6 |  |  |  |
|  | Month | 2 | 4882 | 5.44 | **0.001** | 498.1 | 26.7 | 4.13 | 2, 21 | 0.065 |
|  | Site type*Year | 1 | 2519 | 2.81 | **0.035** | 270.3 | 14.5 | 2.86 | 3, 20 | 0.115 |
|  | Site type*Month | 2 | 430 | 0.48 | 0.869 | 0.0 | 0.0 |  |  |  |
|  | Year*Month | 2 | 669 | 0.75 | 0.663 | 0.0 | 0.0 |  |  |  |
|  | Site type*Year*Month | 2 | 667 | 0.74 | 0.666 | 0.0 | 0.0 |  |  |  |
|  | Error (i.e., Net sample) | 12 | 897 |  |  | 897.2 | 48.1 |  |  |  |
| Platform | Site type | 1 | 3766 | 2.91 | **0.007** | 137.3 | 4.2 | 1.19 | 1, 34 | 0.341 |
|  | Year | 1 | 5799 | 4.48 | 0.001 | 250.2 | 7.7 | 2.75 | 1, 34 | 0.147 |
|  | Month | 2 | 6874 | 5.31 | 0.001 | 464.9 | 14.3 | 10.94 | 2, 33 | 0.003 |
|  | Site type*Year | 1 | 2368 | 1.83 | 0.100 | 119.2 | 3.7 |  |  |  |
|  | Site type*Month | 2 | 1591 | 1.23 | 0.287 | 49.4 | 1.5 |  |  |  |
|  | Year*Month | 2 | 5943 | 4.59 | **0.001** | 774.7 | 23.8 | 6.53 | 5, 30 | 0.011 |
|  | Site type*Year*Month | 2 | 1771 | 1.37 | 0.200 | 158.6 | 4.9 |  |  |  |
|  | Error (i.e., Haul sample) | 24 | 1295 |  |  | 1295.0 | 39.9 |  |  |  |
| Pool abiotic conditions | Site type | 1 | 31.5 | 11.56 | 0.001 | 1.2 | 10.9 | 44.78 | 1, 45 | 0.001 |
|  | Year | 1 | 12.8 | 4.69 | 0.008 | 0.4 | 3.8 | 7.15 | 1, 45 | 0.017 |
|  | Month | 2 | 30.4 | 14.08 | 0.001 | 1.8 | 16.0 | 1.17 | 2, 44 | 0.486 |
|  | Site type*Year | 1 | 5.5 | 2.04 | 0.103 | 0.2 | 2.1 |  |  |  |
|  | Site type*Month | 2 | 8.1 | 3.76 | 0.001 | 0.8 | 6.8 | 12.76 | 5, 41 | 0.001 |
|  | Year*Month | 2 | 24.2 | 11.18 | 0.001 | 2.9 | 25.0 | 2.56 | 5, 41 | 0.163 |
|  | Site type*Year*Month | 2 | 8.7 | 4.02 | **0.002** | 1.7 | 14.8 | 8.02 | 11, 35 | 0.002 |
|  | Pool(Site type*Year) | 12 | 2.7 | 1.26 | 0.173 | 0.2 | 1.7 |  |  |  |
|  | Month*Pool(Site type*Year) (i.e., Sample) | 23 | 2.2 |  |  | 2.2 | 18.9 |  |  |  |
| Invertebrate activity trap | Site type | 1 | 70278 | 55.89 | 0.001 | 2995.7 | 50.0 | 47.83 | 1, 45 | 0.001 |
|  | Year | 1 | 2116 | 1.68 | 0.158 | 37.3 | 0.6 |  |  |  |
|  | Month | 2 | 7911 | 6.65 | 0.001 | 436.9 | 7.3 | 0.26 | 2, 44 | 0.794 |
|  | Site type*Year | 1 | 1442 | 1.15 | 0.329 | 16.0 | 0.3 |  |  |  |
|  | Site type*Month | 2 | 8921 | 7.50 | **0.001** | 1005.1 | 16.8 | 15.44 | 5, 41 | 0.001 |
|  | Year*Month | 2 | 1112 | 0.94 | 0.464 | 0.0 | 0.0 |  |  |  |
|  | Site type*Year*Month | 2 | 2274 | 1.91 | 0.074 | 282.0 | 4.7 |  |  |  |
|  | Pool(Site type*Year) | 12 | 1258 | 1.06 | 0.382 | 23.5 | 0.4 |  |  |  |
|  | Month*Pool(Site type*Year) (i.e., Sample) | 23 | 1190 |  |  | 1189.6 | 19.9 |  |  |  |
| Minnow trap | Site type | 1 | 19907 | 3.61 | **0.036** | 624.6 | 15.3 | 0.21 | 1, 45 | 0.668 |
|  | Year | 1 | 2258 | 0.41 | 0.789 | 0.0 | 0.0 |  |  |  |
|  | Month | 2 | 2765 | 1.28 | 0.222 | 39.8 | 1.0 |  |  |  |
|  | Site type*Year | 1 | 6562 | 1.19 | 0.310 | 90.9 | 2.2 |  |  |  |
|  | Site type*Month | 2 | 2310 | 1.07 | 0.380 | 20.5 | 0.5 |  |  |  |
|  | Year*Month | 2 | 2113 | 0.98 | 0.436 | 0.0 | 0.0 |  |  |  |
|  | Site type*Year*Month | 2 | 1804 | 0.84 | 0.546 | 0.0 | 0.0 |  |  |  |
|  | Pool(Site type*Year) | 12 | 5558 | 2.58 | 0.001 | 1167.4 | 28.5 |  |  |  |
|  | Month*Pool(Site type*Year) (i.e., Sample) | 23 | 2153 |  |  | 2152.5 | 52.6 |  |  |  |

#### 3.1.2. Marsh platform communities in Aulac

Species diversity (∼3.0) and evenness (∼0.21) did not differ significantly between reference and restoring platforms in Aulac. However, species richness differed by site type depending on month, while total number of nekton individuals differed by site type depending on year and month (Tables S5.1 and S5.2). Generally, reference platforms supported higher species richness (∼5.3 vs. ∼4.6; Table S5.1) and nekton numbers (Table 1). Nekton community composition differed by site type independent of years and months, as well as showed temporal (Year*Month) variation (PERMANOVA, Table 2, Figs. 2 and 3b). Both differences in centroids and dispersion in community composition contributed to the temporal pattern (PERMDISP, Table 2); the restoring community showed high dispersion in July (Fig. 3b). Species contributing most to site type community dissimilarity (52%) were a relatively wide set, including consistently silversides and blackspotted sticklebacks, as well as fourspine (*Apeltes quadracus*), ninespine sticklebacks, juvenile rainbow smelts (*Osmerus mordax*), mummichogs, and juvenile eels, which were more abundant on reference platforms (SIMPER, Table S6.1, Fig. 2). Tomcods were more abundant on restoring platforms.

#### 3.1.3. Spatiotemporal variation in the visiting nekton community in Aulac

The Site type effect (summing components of variation for the main effect and its interaction with Year and/or Month) accounted for considerable (23% for creeks) or moderate (14% for platforms) amount of the nekton community variation in Aulac (Table 2). Temporal variation in nekton community composition was also considerable to moderate due to Month (27% for creeks and 14% for platform), and moderate to low due to Year (3□8%); variation accounted by the Year*Month interaction was non-existent (0%) for creek communities, but considerable (24%) for platform communities. Our smallest spatiotemporal scale (replicate nets and hauls) accounted for sizeable amount of variation (40□48%) in creek and platform nekton communities.

### 3.2. Environmental conditions and fauna in Aulac pools

#### 3.2.1. Abiotic conditions and aquatic vegetation in Aulac pools

The abiotic conditions in Aulac pools differed between the reference and restoring sites depending on month and year (significant Site type*Year*Month interaction, PERMANOVA, Table 2), due to both differences in centroids and dispersion (PERMDISP, Table 2, Fig. 4a). For both 2022 and 2023, the conditions differed by site type depending on month, again due to both differences in centroids and dispersion (significant Site*Month interaction, PERMANOVA and PERMDISP, Table S7.1). For the main contributing variables, water depth (mean: 29 vs 18 cm) and pH (7.9 vs. 7.7) were greater, while sediment penetrability (24 vs 36 cm) and water salinity (17.7 vs. 18.3 ppt) were lower in reference pools compared to restoring pools (SIMPER, Tables S6.1 and S8.1). Water dissolved oxygen concentration (6.7 vs. 6.5 mg/L) showed much variability between sampling events, and similar overall averages in reference and restoring pools.

**Fig. 4.**
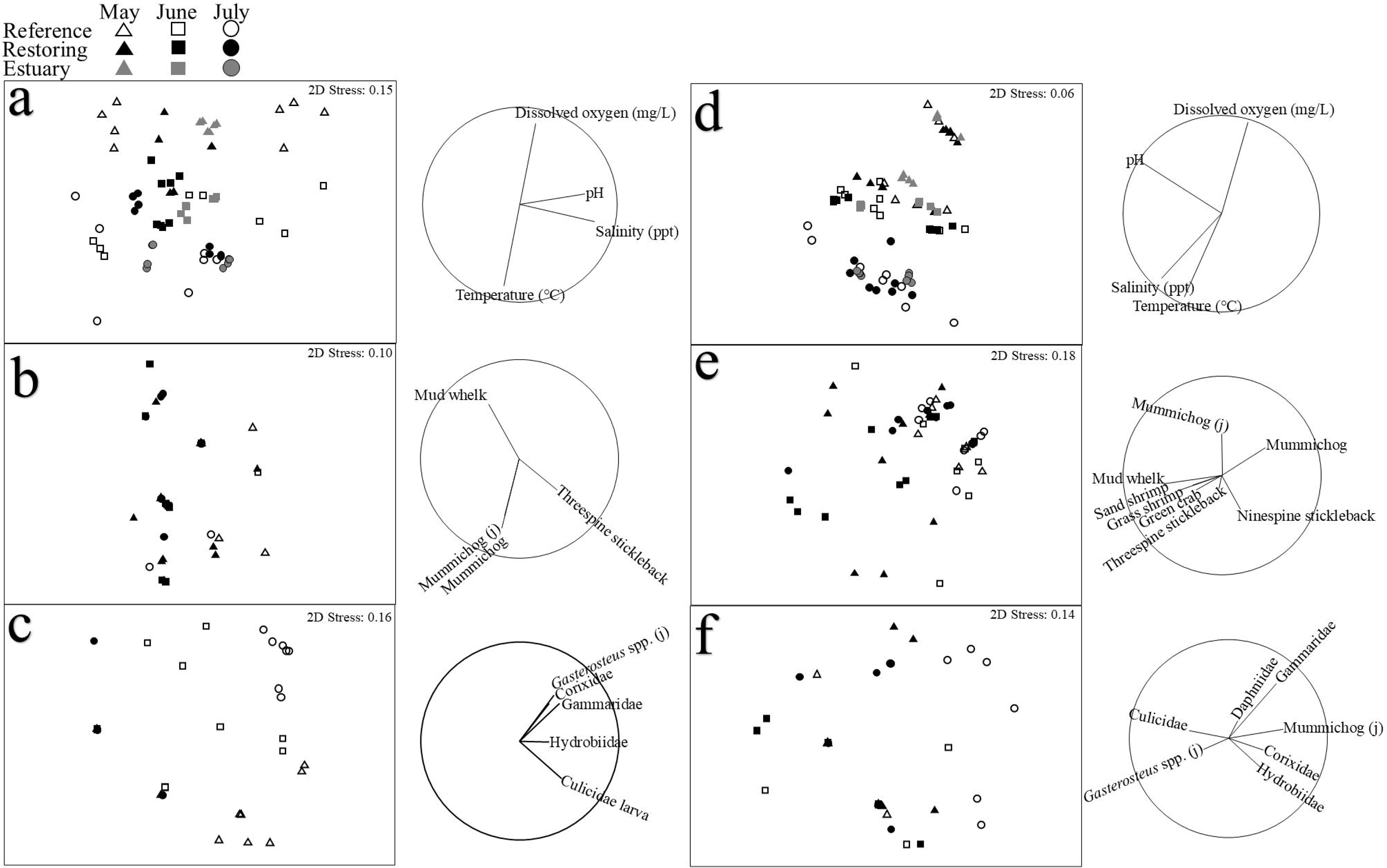
nMDS graphs for salt pools: (a,d) abiotic conditions (water temperature, salinity, pH, and dissolved oxygen) and biotic communities captured in (b,e) minnow traps and (c,f) invertebrate activity traps in reference and restoring salt marshes in (a–c) Aulac and (d–f) Wallace Bay for 3 months (May–July) in 2022–2023. A symbol represents the abiotic conditions (normalized) or community composition (4^th^ root transformed) averaged per Year-Month-Site type combinations. Juvenile life stage of fish species indicated by “(j)”. *Gasterosteus* spp. are threespine (*G. aculeatus*) and blackspotted (*G. wheatlandi*) sticklebacks. See Fig. 3 caption for explanation of vector diagrams and stress values.

Percent cover of aquatic vegetation in June and July was significantly higher in reference pools (37% and 24% for *Ruppia* and macroalgae, respectively) than in restoring pools (0% and 1%), with some yearly variation (Tables S8.3 and S8.4).

#### 3.2.2. Invertebrate activity trap fauna in Aulac pools

The composition of invertebrate activity trap communities in Aulac differed by site type depending on month, due to differences in centroids and dispersion (significant Site type*Month interaction, PERMANOVA and PERMDISP, Table 2, Fig. 4c), with an overall community dissimilarity of 100% (SIMPER, Table S6.1). Traps from restoring pools were mostly empty (except for a few hydrobiid snails and *Corophium* amphipods), while those in reference pools were typically dominated by water boatmen (Corixidae), mosquito larvae (Culicidae), hydrobiid snails, gammarid amphipods, and also had small juvenile *Gasterosteus* spp. and mummichogs (Tables 1 and S9.1).

#### 3.2.3. Minnow trap fauna in Aulac pools

Species richness (∼1.0), diversity (∼0.4), and evenness (∼0.04) and the total number of nekton individuals captured in minnow traps did not differ significantly between reference and restoring pools in Aulac (Tables 1, S5.1 and S5.2, Fig. 2). However, composition of minnow trap communities differed significantly by site type independent of months and years (PERMANOVA, Table 2, Fig. 4b). Species contributing most to community dissimilarity between site types (87%) were threespine sticklebacks being more abundant in reference pools, while large juvenile and adult mummichogs, and mud whelks (*Tritia obsoleta*) being more so in restoring pools (SIMPER, Table S6.1, Fig. 2). Adult mummichogs were also consistent in this pattern (based on average dissimilarity/SD of dissimilarities > 1; Table S6.1).

#### 3.2.4. Spatiotemporal variation in environmental conditions and small-bodied faunal communities in Aulac pools

The Site type effect (main effect and its interaction with Year and/or Month) accounted for a considerable amount of the variation in pools (35% for abiotic conditions, 63% for aquatic vegetation percent cover, 72% invertebrate activity trap communities, and 18% for minnow trap communities in Aulac (Tables 2 and S8.4). The smaller spatial scale of Pool accounted for considerable variation for minnow traps (28%) and moderate variation for aquatic vegetation (8%), but very low variation (<2%) for abiotic conditions and invertebrate activity traps. Yearly variation was low for most responses (0 4%) except aquatic vegetation (13%). Other temporal variation differed substantially depending on the response (0 25%); in particular, the Year*Month interaction accounted for considerable variation (25%) in pool abiotic conditions. Our smallest spatiotemporal scale (sample) accounted for moderate to considerable variation (14 20%) in abiotic conditions, aquatic vegetation and invertebrate activity traps, and for most of the minnow trap variation (53%).

### 3.3. The visiting nekton community in Wallace Bay

Over all sampling rounds in Wallace Bay, fyke nets in reference creeks caught 10,564 individuals and 16 species of nekton, while those in restoring creeks captured 4,969 individuals and 23 species (Table 1). Seine hauls captured 427 individuals and 7 species on reference platforms, while those on restoring platforms caught 1056 individuals and 10 species. Reference creeks and platforms were dominated by mummichogs, while sand shrimps and Atlantic silversides were most abundant in restoring creeks and on restoring platforms, respectively.

#### 3.3.1. Creek communities in Wallace Bay

Nekton diversity (∼2.2), evenness (∼0.10), and total number of individuals did not differ significantly between reference and restoring creeks in Wallace Bay. However, richness differed by site type depending on month and year (Tables 1, S5.1 andS5.2, Fig. 5); generally, richness was higher in restoring creeks (∼9.3 vs. ∼7.0; except for May 2022). The composition of creek communities also differed by site type depending on month (significant Site type*Month, PERMANOVA, Table 3, Fig. 3c). Species contributing most to community dissimilarity between site types (43%) were juvenile and adult mummichogs being consistently most abundant in reference creeks, while sand shrimps, grass shrimps (*Palaemon paludosus*), green crabs (*Carcinus maenas*), white perch (*Morone americana*), and white tipped mud crabs (*Rhithropanopeus harrisii*) being more abundant in restoring creeks (SIMPER, Table S6.1).

**Fig. 5.**
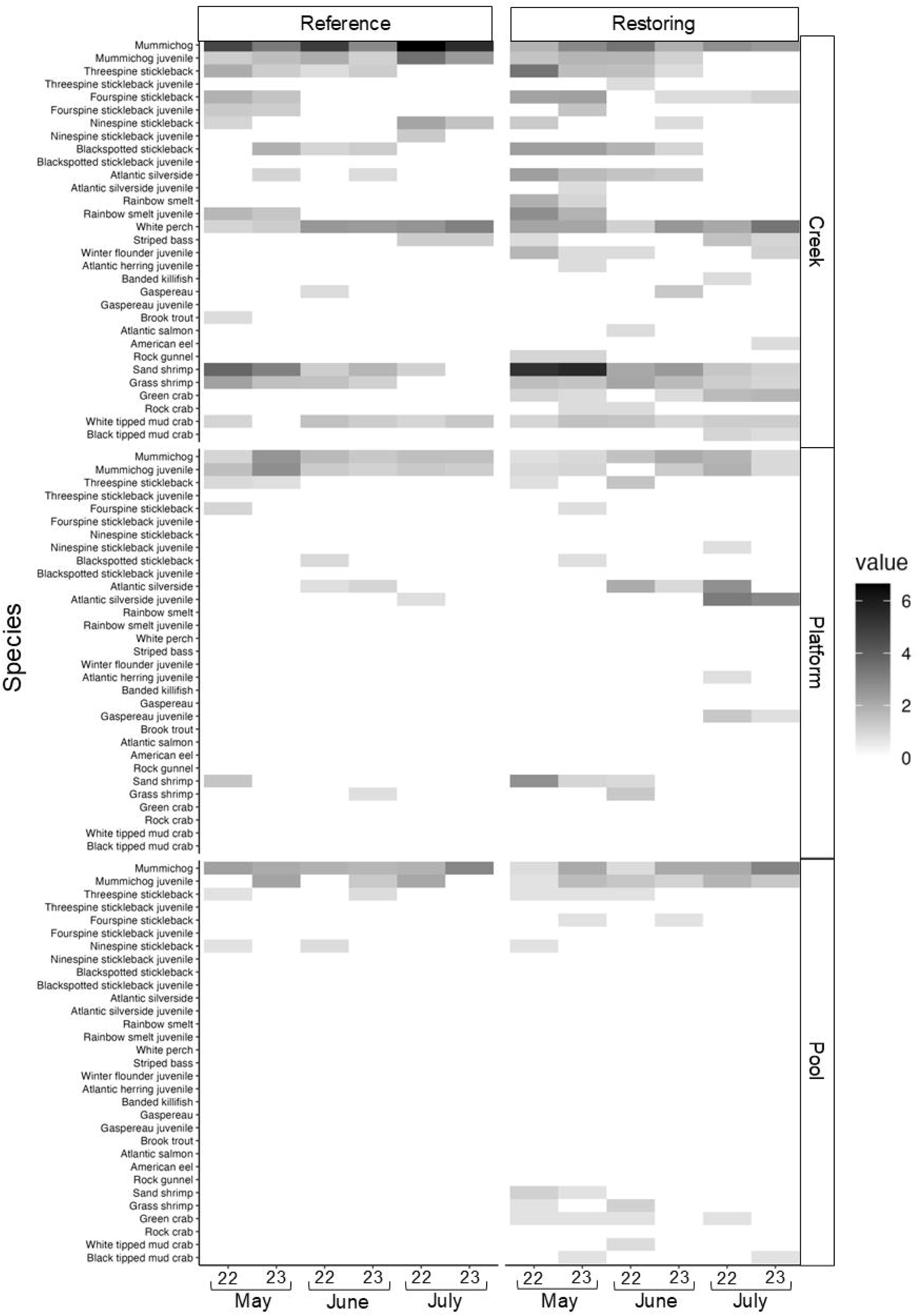
Heatmap showing average densities of nekton taxa (per net or trap; 4^th^ root transformed data) captured in three subhabitats in the Wallace Bay reference and restoring salt marshes for three months (May–July) in 2022–2023. Averaged over 2 fyke nets (creeks), 3 seine hauls (platform), or 4 minnow traps (salt pools) per sampling round per site type (*n* = 2–4). A “J” following a fish species name indicates juveniles and those without are adults. Invertebrates were not divided by life stage. Gaspereau are alewives (*Alosa aestivalis*) and blueback herring (*A. pseudoharengus*).

**Table 3.** PERMANOVA results comparing fish and invertebrate communities in Wallace Bay reference and restoring salt marshes in May–July 2022–2023; analyses done separately for creek, marsh platform, and salt pool subhabitats. See Table S1 for dates of sampling rounds, Table 1 for taxa included in each community, and Table S8.1 for abiotic variables measured in salt pools. Site type, Year and Month are fixed factors, and Pool is a random factor. Taxa densities 4th root transformed and abiotic variables normalized prior to analysis. P-values bolded for significant and interpretable fixed effects; 997–999 unique permutations. PERMDISP tests conducted for significant fixed effects to assess amount of multivariate dispersion among groups; df1 and df2 represent the numerator and denominator degrees of freedom for the F-ratio, respectively. Analysis of components of variation conducted for both fixed and random effects to evaluate relative importance of spatiotemporal scales.

| Community or assemblage | Sources of variation | PERMANOVA |  |  |  | Components of variation |  | PERMDISP |  |  |
| --- | --- | --- | --- | --- | --- | --- | --- | --- | --- | --- |
|  |  | df | MS | Pseudo-F | P(perm) | Estimate | % | F | df 1, df 2 | P |
| Creek | Site type | 1 | 4534 | 9.28 | 0.002 | 337.1 | 17.3 | 1.68 | 1, 22 | 0.164 |
|  | Year | 1 | 880 | 1.80 | 0.115 | 32.7 | 1.7 |  |  |  |
|  | Month | 2 | 6355 | 13.01 | 0.001 | 733.3 | 37.6 | 1.06 | 2, 21 | 0.413 |
|  | Site type*Year | 1 | 532 | 1.09 | 0.364 | 7.3 | 0.4 |  |  |  |
|  | Site type*Month | 2 | 1402 | 2.87 | **0.005** | 228.4 | 11.7 | 4.05 | 5, 18 | 0.057 |
|  | Year*Month | 2 | 742 | 1.52 | 0.121 | 63.5 | 3.3 |  |  |  |
|  | Site type*Year*Month | 2 | 610 | 1.25 | 0.272 | 60.7 | 3.1 |  |  |  |
|  | Error (i.e., Net sample) | 12 | 488 |  |  | 488.4 | 25.0 |  |  |  |
| Platform | Site type | 1 | 7298 | 6.59 | 0.002 | 343.9 | 8.2 | 7.07 | 1, 34 | 0.028 |
|  | Year | 1 | 2478 | 2.24 | 0.119 | 76.1 | 1.8 |  |  |  |
|  | Month | 2 | 11179 | 10.09 | 0.001 | 839.3 | 20.0 | 0.28 | 2, 33 | 0.789 |
|  | Site type*Year | 1 | 3545 | 3.20 | 0.031 | 270.8 | 6.4 | 3.91 | 3, 32 | 0.046 |
|  | Site type*Month | 2 | 5136 | 4.64 | 0.001 | 671.3 | 16.0 | 0.72 | 5, 30 | 0.767 |
|  | Year*Month | 2 | 1054 | 0.95 | 0.477 | 0.0 | 0.0 |  |  |  |
|  | Site type*Year*Month | 2 | 3794 | 3.42 | **0.013** | 895.4 | 21.3 | 3.87 | 11, 24 | 0.075 |
|  | Error (i.e., Haul sample) | 24 | 1108 |  |  | 1108.0 | 26.4 |  |  |  |
| Pool abiotic conditions | Site type | 1 | 1.7 | 0.56 | 0.591 | 0.0 | 0.0 |  |  |  |
|  | Year | 1 | 36.5 | 11.90 | 0.001 | 1.4 | 15.2 | 10.01 | 1, 46 | 0.005 |
|  | Month | 2 | 49.3 | 20.95 | 0.001 | 2.9 | 31.9 | 0.05 | 2, 45 | 0.957 |
|  | Site type*Year | 1 | 5.3 | 1.71 | 0.177 | 0.2 | 2.0 |  |  |  |
|  | Site type*Month | 2 | 7.6 | 3.24 | **0.005** | 0.7 | 7.2 | 1.45 | 5, 42 | 0.255 |
|  | Year*Month | 2 | 13.8 | 5.85 | **0.001** | 1.4 | 15.5 | 1.76 | 5, 42 | 0.243 |
|  | Site type*Year*Month | 2 | 2.1 | 0.88 | 0.539 | 0.0 | 0.0 |  |  |  |
|  | Pool(Site type*Year) | 12 | 3.1 | 1.30 | 0.168 | 0.2 | 2.6 |  |  |  |
|  | Month*Pool(Site type*Year) (i.e., Sample) | 24 | 2.4 |  |  | 2.4 | 25.6 |  |  |  |
| Invertebrate activity trap | Site type | 1 | 4509 | 2.32 | 0.054 | 106.9 | 1.8 | 0.23 | 1, 46 | 0.634 |
|  | Year | 1 | 10763 | 5.54 | 0.003 | 367.5 | 6.3 | 2.24 | 1, 46 | 0.159 |
|  | Month | 2 | 6677 | 2.02 | 0.044 | 211.2 | 3.6 | 1.04 | 2, 45 | 0.034 |
|  | Site type*Year | 1 | 3260 | 1.68 | 0.153 | 109.7 | 1.9 |  |  |  |
|  | Site type*Month | 2 | 6128 | 1.86 | 0.097 | 353.7 | 6.1 |  |  |  |
|  | Year*Month | 2 | 5971 | 1.81 | 0.090 | 334.2 | 5.7 |  |  |  |
|  | Site type*Year*Month | 2 | 7543 | 2.29 | **0.039** | 1061.2 | 18.2 | 3.70 | 11, 36 | 0.024 |
|  | Pool(Site type*Year) | 12 | 1943 | 0.59 | 0.972 | 0.0 | 0.0 |  |  |  |
|  | Month*Pool(Site type*Year) (i.e., Sample) | 24 | 3298 |  |  | 3297.6 | 56.4 |  |  |  |
| Minnow trap | Site type | 1 | 13643 | 13.48 | 0.001 | 526.3 | 12.9 | 9.05 | 1, 46 | 0.025 |
|  | Year | 1 | 3560 | 3.52 | 0.023 | 106.2 | 2.6 | 2.24 | 1, 46 | 0.273 |
|  | Month | 2 | 4347 | 3.14 | 0.009 | 185.1 | 4.5 | 5.22 | 2, 45 | 0.053 |
|  | Site type*Year | 1 | 5509 | 5.44 | 0.003 | 374.8 | 9.2 | 5.64 | 3, 44 | 0.013 |
|  | Site type*Month | 2 | 3485 | 2.52 | 0.013 | 262.5 | 6.4 | 3.18 | 5, 42 | 0.071 |
|  | Year*Month | 2 | 5311 | 3.83 | 0.003 | 490.6 | 12.0 | 5.66 | 5, 42 | 0.006 |
|  | Site type*Year*Month | 2 | 4421 | 3.19 | **0.006** | 758.8 | 18.6 | 3.81 | 11, 36 | 0.058 |
|  | Pool(Site type*Year) | 12 | 1012 | 0.73 | 0.883 | 0.0 | 0.0 |  |  |  |
|  | Month*Pool(Site type*Year) (i.e., Sample) | 24 | 1386 |  |  | 1385.5 | 33.9 |  |  |  |

#### 3.3.2. Platform nekton communities in Wallace Bay

Nekton diversity (∼1.6 in reference vs ∼2.4 in restoring), richness (∼1.3 vs ∼1.6), evenness (∼0.13 vs ∼0.16), and total number of individuals differed by site type depending on month and year and were generally higher on restoring platforms in Wallace Bay (Table 1, S5.1 and S5.2, Fig. 5). The platform community composition differed by site type depending on year and month (significant Site type*Year*Month, PERMANOVA, Table 3, Fig. 3d). For both 2022 and 2023, platform communities still differed by site type depending on month (Table S7.1). The species contributing most to community dissimilarity between site types (64%) were juvenile and adult mummichogs being consistently more abundant on reference platforms, while juvenile and adult silversides, threespine sticklebacks, sand shrimps, and grass shrimps being more numerous on restoring platforms (SIMPER, Table S6.1).

#### 3.3.3. Spatiotemporal variation in the visiting nekton community in Wallace Bay

The Site type effect (main effect and its interaction with Year and/or Month) in Wallace Bay accounted for a considerable amount of nekton community variation (33% for creeks and 52% for platform; Table 3). Temporal variation was also considerable for Month (38% for creeks and 20% for platforms), but low for Year and Year-Month (≤3%). Our smallest spatiotemporal scale (replicate nets and hauls) accounted for sizable community variation in creeks (25%) and on platforms (26%).

### 3.4. Environmental conditions and fauna in Wallace Bay salt pools

#### 3.4.1. Abiotic conditions and aquatic vegetation in Wallace Bay pools

Abiotic conditions in salt pools differed by site type in Wallace Bay depending on month (significant Site type*Month interaction, PERMANOVA, Table 3, Fig. 4d). Water depth (mean: 27 vs. 18 cm) and sediment penetrability (27 vs. 25 cm) were higher in reference compared to restoring pools, and contributed most to abiotic differences (SIMPER, Tables S6.1 and S8.1). Water pH (∼7.8), salinity (16.0 vs 16.7 ppt) and dissolved oxygen concentration (7.3 vs. 6.8 mg/L) showed variability among sampling events, and so were on average more similar for reference and restoring pools (Table S8.1).

Percent cover of aquatic vegetation in June and July was significantly higher in reference pools (40% and 31% for *Ruppia* and macroalgae, respectively) than in restoring pools (6% and 8%), with some temporal (monthly and yearly) variation (Tables S8.3 and S8.4).

#### 3.4.2. Invertebrate activity trap fauna in Wallace Bay pools

Taxon richness (∼1.33 in reference vs ∼1.23 in restoring), diversity (∼1.17 vs ∼1.23), and evenness (∼0.13 vs ∼0.14) in invertebrate activity trap communities differed between reference and restoring pools depending on month in Wallace Bay (Tables 1, S5.1, S5.3, and S9.1). More individuals were captured in reference invertebrate activity traps overall (Table 1); however, the total number of individuals did not differ significantly between site types in Wallace Bay (Table S5.3). The indices were higher in restoring traps in some rounds (e.g., May 2022) and lower in other rounds (e.g., July 2022), and total number of individuals was generally lower in restoring traps. Community composition differed by site type depending on year and month, due to differences in both centroids and dispersion (significant Site type*Year*Month interaction, PERMANOVA and PERMDISP, Table 3, Fig. 4f). In 2022, communities differed by site type depending on month, and in 2023, they differed by site type only (Table S7.1). Taxa contributing most to overall community dissimilarity (82%) were hydrobiid snails, gammarid amphipods, water boatmen, and small juvenile mummichogs and sticklebacks, being more abundant in reference pools, and mosquito (Culicidae) larvae being more so in restoring pools (SIMPER, Table S6.1, Table 1).

#### 3.4.3. Minnow trap fauna in Wallace Bay pools

Species diversity (∼1.7) and total number of individuals in minnow trap communities did not differ significantly between site types in Wallace Bay; however, species richness (∼1.6 in restoration vs ∼2.6 in restoring) and evenness (∼0.11 vs ∼0.17) were higher in restoring pools (Table 1, S5.1 and S5.2). The composition of minnow trap communities differed by site type depending on year and month (significant Site type*Year*Month interaction, PERMANOVA, Table 3, Figs.2 and 4e), due to differences in community centroids and dispersion (PERMDISP, Table 2). In 2022, minnow trap communities differed by site type depending on month, and in 2023, they differed by site type only (Table S7.1). The main species contributing to community dissimilarity between site types (61%) were adult mummichogs consistently being more abundant in reference pools, and large juvenile mummichogs, mud whelks, sand shrimps, and threespine sticklebacks being more abundant in restoring pools (SIMPER, Table S6.1).

#### 3.4.4. Spatiotemporal variation in pool environmental conditions and small-bodied fauna in Wallace Bay

The Site type effect (main effect and its interaction with Year and/or Month) in Wallace Bay accounted for a moderate to sizeable amount of the variation in pools (9% for abiotic conditions, 67% for aquatic vegetation percent cover, 28% for invertebrate activity trap communities, and 47% for minnow trap communities; Tables 3 and S8.4). The spatial scale of pool accounted for low variation (<3%) in abiotic conditions, aquatic vegetation, invertebrate activity traps, and minnow traps. Temporal variation (yearly, monthly and for the Year-Month interaction) was considerable for abiotic conditions (15 32%), and low to moderate for aquatic vegetation (2 9%), invertebrate activity traps (4 6%) and minnow traps (3 12%). Our smallest spatiotemporal scale (sample) accounted for sizable variation in abiotic conditions (26%), aquatic vegetation (16%), invertebrate activity traps (56%), and minnow traps (34%).

### 3.5. Fish foraging patterns in Aulac and Wallace Bay salt marshes

#### 3.5.1. Tomcod and mummichog gut fullness and prey assemblage in Aulac

For both tomcods (captured in creeks) and mummichogs (captured in creeks and platforms), gut fullness did not differ significantly by site type in Aulac (PERMANOVA, Table 4, Fig. 6). Gut fullness index measured as weight of ingested food relative to body weight of intact fish averaged 3.2 ± 0.2% (± SE) for tomcods, 3.3 ± 0.2% for creek mummichogs, and 3.7 ± 0.3% for platform mummichogs, reflecting partially full guts (Table S4.7). Only one fish (a tomcod in the restoring site in July) appeared to have an empty gut.

**Fig. 6.**
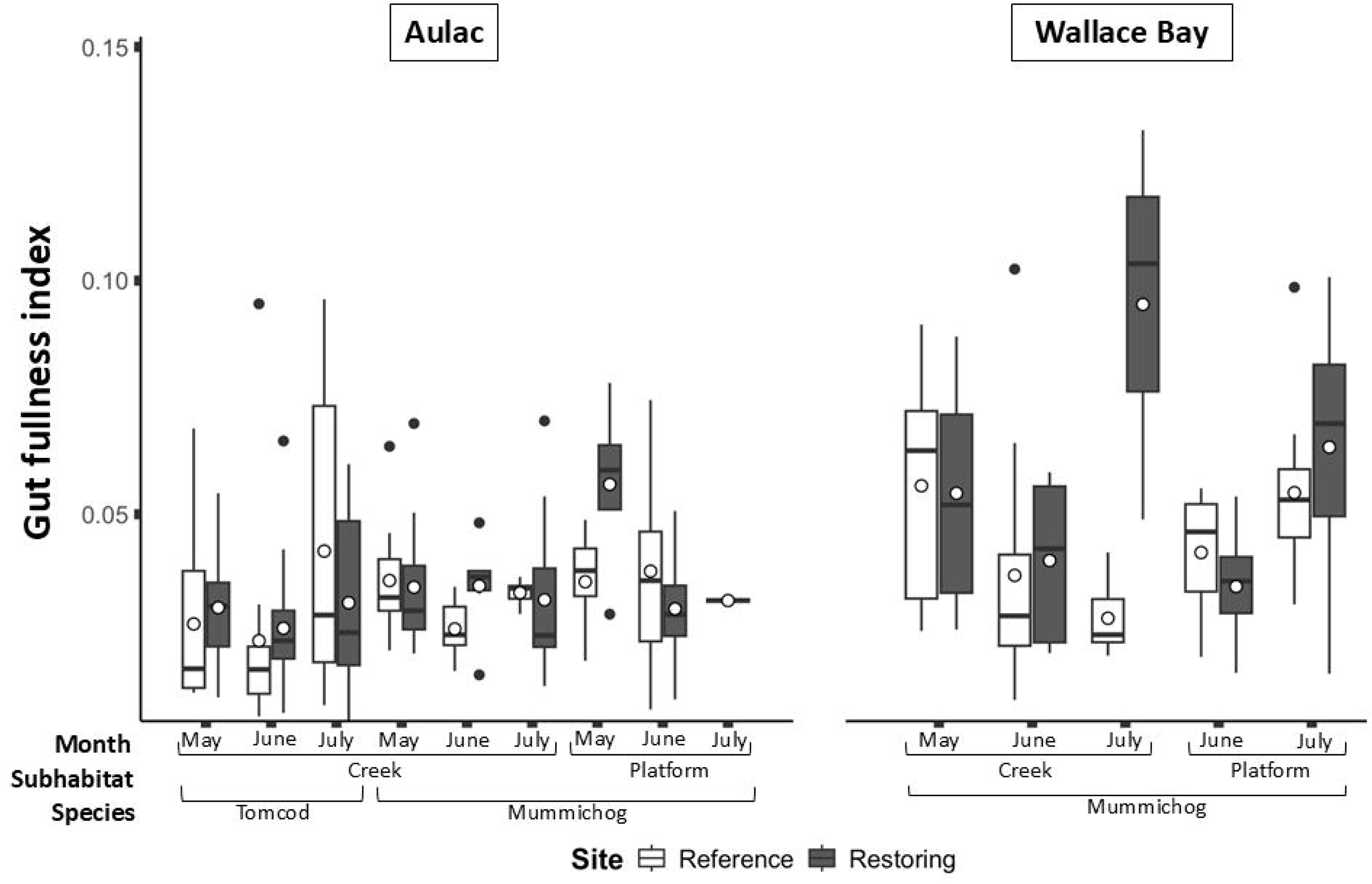
Gut fullness indices (ratio of gut content wet weight to intact fish wet weight) for tomcods (*Microgadus tomcod*) and mummichogs (*Fundulus heteroclitus*) captured in creeks and platforms in reference and restoring salt marshes in Aulac and Wallace Bay. Fish were captured in May–July in 2022–2023, and are graphed pooling over years. We aimed to retain the same number of fish for both site types and subhabitats during each round; however, this was not possible (*n =* 1–10 fish depending on Year-Month-Subhabitat-Species combination; Table S4.1). Tomcods were only retained from creeks in Aulac; no tomcods were caught in Wallace Bay. Only 1 mummichog was retained from the reference platform in Aulac in July (over both years), and no platform mummichogs were captured from either site type in Wallace Bay in May of both years. In box plots, white circle and thick horizontal bar are mean and median, respectively; lower and upper box edges are 25^th^ and 75^th^ quartiles, respectively; whiskers are 1.5 x interquartile range, and black dots are values outside this range.

**Table 4.**
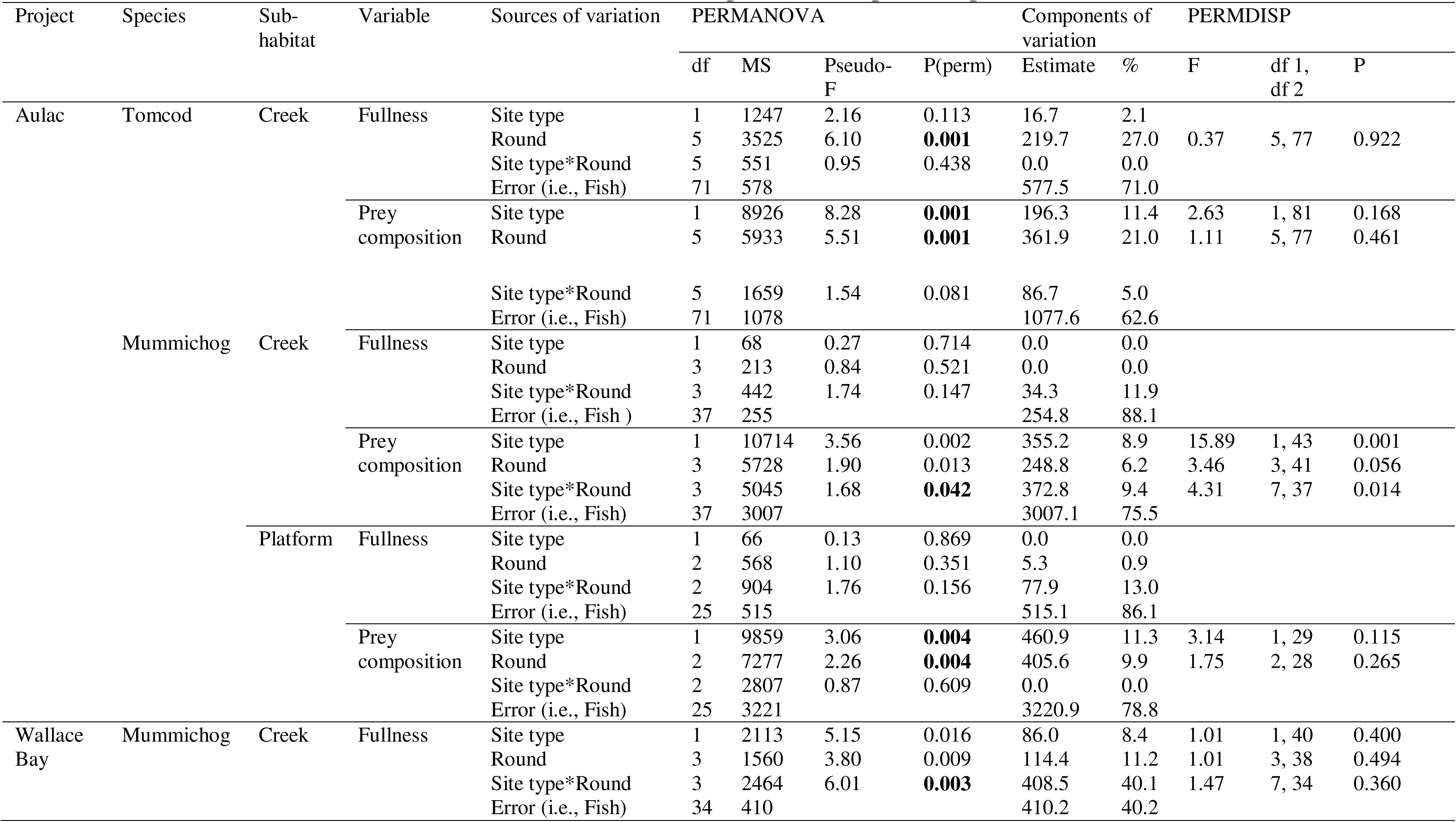
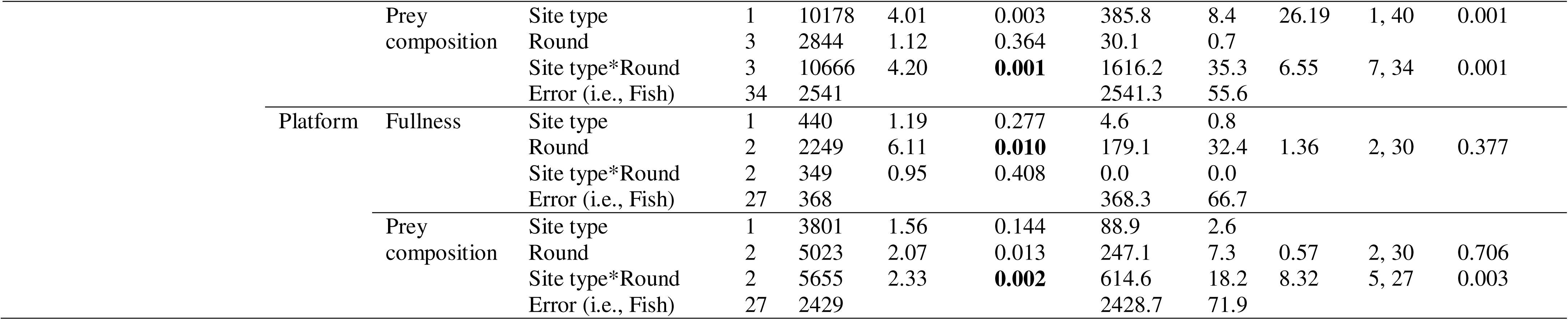
PERMANOVA results for gut fullness and prey composition for tomcods (*Microgadus tomcod*) and mummichogs (*Fundulus heteroclitus*) in Aulac and Wallace Bay reference and restoring salt marsh creeks and platforms in 2022–2023; analyses done separately for each species and each subhabitat. Round indicates a year-month combination; see Table S1 for dates of sampling rounds and Table S4.1 for number of fish retained from each round. Site type and Round are fixed factors. Taxa densities converted to presence-absence prior to analysis. P-values bolded for significant and interpretable fixed effects; 998–999 unique permutations. PERMDISP tests conducted for significant fixed effects to assess amount of multivariate dispersion among groups; df1 and df2 represent the numerator and denominator degrees of freedom for the F-ratio, respectively. Analysis of components of variation conducted for both fixed and random effects to evaluate relative importance of spatiotemporal scales.

Prey assemblage for tomcods captured in creeks differed by site type (PERMANOVA, Table 4). *Corophium volutator* amphipods were the most prevalent prey in tomcods from both site types (Fig. 7). Prey taxa contributing most to the moderate compositional dissimilarity (50%) between site types were gammarid amphipods, nereid polychaetes, and vegetal detritus being more consistently present in reference creek tomcods, while sand shrimps being more so in restoring creek tomcods (SIMPER, Table S6.2). Note that prey composition transformed to presence-absence (to include vegetal matter) or using 4^th^-root transformation of fauna prey counts (excluding vegetal matter) showed similar patterns (Tables 4 and S4.5).

**Fig. 7.**
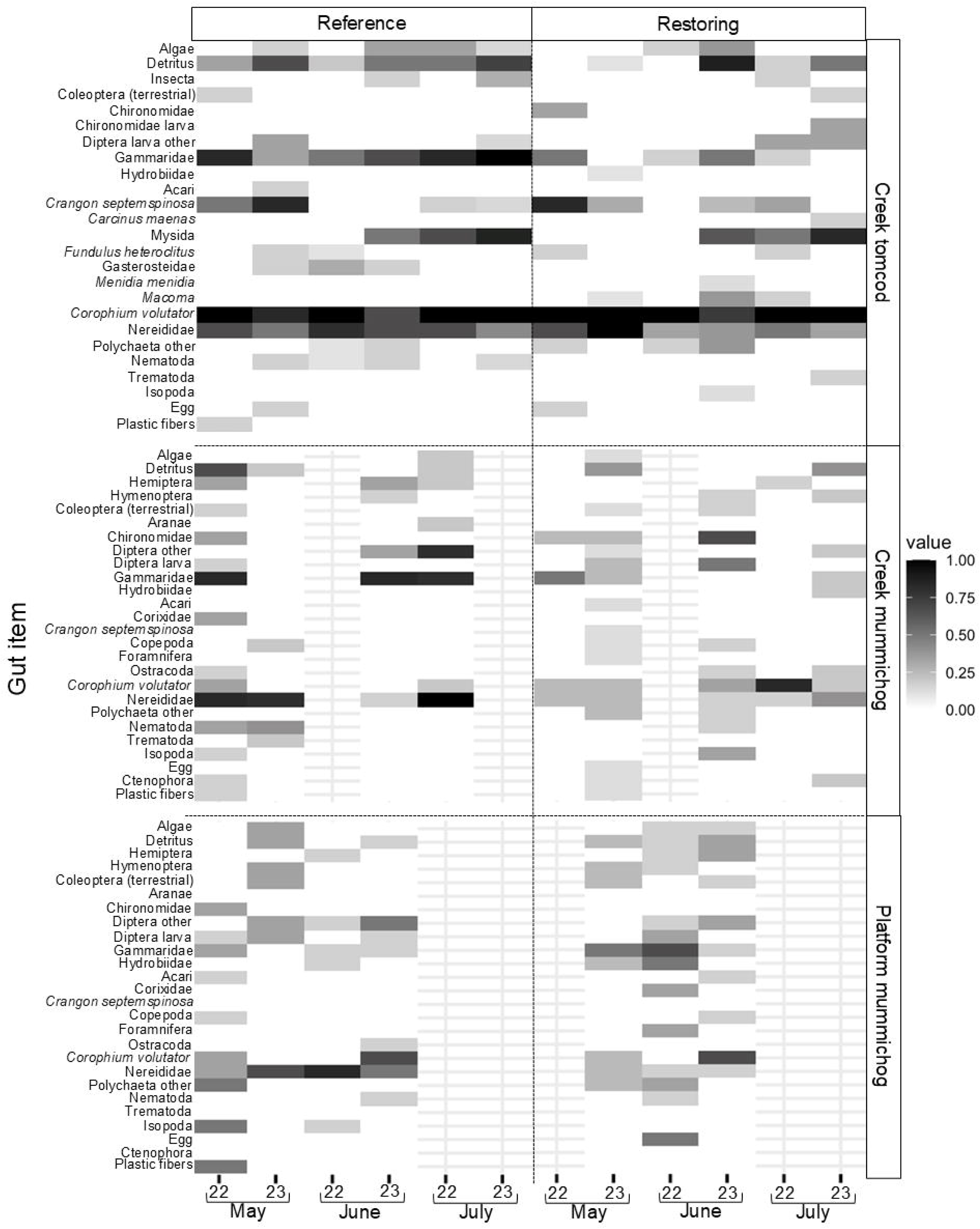
Heatmap showing average presence-absence of gut items in tomcods (*Microgadus tomcod*) and mummichogs (*Fundulus heteroclitus*) captured in creeks and on platforms in Aulac reference and restoring salt marshes for three months (May–July) in 2022–2023. A value of 1 (black) represents consistently present, 0 (white) consistently absent, and shades of grey are in between. Averaged over fish captured in the ebb fyke net (creeks) and in seine hauls (platform) per sampling round per site type (*n* = 3–10, Table S4.3). An “A” or an “L” following a taxa name indicates an adult or larval life stage. Areas with gridlines represent rounds when no fish were captured. The order of prey listed on a heat map reflects nekton prey first, then invertebrate prey typical of salt marshes followed by those typical of mudflats, and then other prey items.

Prey composition of mummichogs captured in creeks differed by site type depending on round (specifically, May and June 2023), due to both differences on centroids and dispersion (PERMANOVA, PERMDISP, and paired comparisons Tables 4 and S4.6). Mummichog gut content was varied; no specific prey item was particularly prevalent in all mummichogs (Fig. 7). Prey taxa contributing most to the high compositional dissimilarity (88%) between site types were gammarid amphipods nereid polychaetes, certain insects (hemipterans and adult dipterans), and vegetal detritus being more consistently present in reference creek mummichogs, while adult chironomids, some larval dipterans, and *Corophium* amphipods being more so in restoring creek mummichogs (SIMPER, Table S6.2).

Prey compositions in mummichogs captured on marsh platforms differed by site type (PERMANOVA, Table 4), with a high dissimilarity (82%, SIMPER, Table S6.2, Fig. 7). Main contributing prey taxa were nereid polychaetes and adult dipterans being more consistently present in reference platform mummichogs, while gammarid, *Corophium* amphipods and hemipterans being more so in restoring platform mummichogs (Table S6.2).

#### 3.5.2. Spatiotemporal variation in tomcod and mummichog gut fullness and prey assemblage in Aulac

The Site type effect (main effect and its interaction with Round) in Aulac accounted for little (2%) of the gut fullness variation for tomcods (in creeks), whereas it accounted for a moderate amount of variation (12-13%) for mummichogs (in creeks and on platforms) (Table 4). However, this Site type effect accounted for moderate to considerable variation (12-18%) in prey composition for tomcods and mummichogs (Tables 4 and S4.5). Temporal variation (Round alone) differed depending on the response: for gut fullness, it was considerable (27%) for creek tomcods, and very low (<1%) for creek and platform mummichogs. For prey composition, variation attributed to sampling round was also considerable (21-26%) for tomcods, but moderate (6-11%) for creek and platform mummichogs. For all analyses, individual fish (our base unit of replication) accounted for the largest amount of variation (57-88%).

#### 3.5.3. Mummichog gut fullness and prey assemblage in Wallace Bay

For creeks, mummichog gut fullness differed by site type depending on round in Wallace Bay (PERMANOVA, Table 4, Fig. 6); average weight of ingested food relative to intact body weight varied among sampling rounds from 2.6-7.6% and 4.0-9.5% for the reference and restoring sites, respectively (Table S4.7). For most sampling rounds, gut indices were comparable between fish captured in reference and restoring creeks. However, two sampling rounds had large differences: average gut fullness was 4.5 times greater in reference than restoring creek fish in May 2022, and ∼2.5 times greater in restoring than reference creek fish in July 2023 (Table S4.6).

Prey composition in creek mummichogs differed by site type depending on round, due to differences in centroids and dispersion (PERMANOVA and PERMDISP, Table 4, Fig. 8). Differences occurred in mummichogs captured in May 2022, and in June and July 2023 (Pairwise comparisons, Table S4.6). Prey taxa contributing most to the large gut assemblage dissimilarity (87%) between site types were nereid polychaetes being more consistently present in reference creek mummichogs, while mysids, sand shrimps, vegetal detritus and macroalgae were more common in restoring creek mummichog (SIMPER, Table S6.2).

**Fig. 8.**
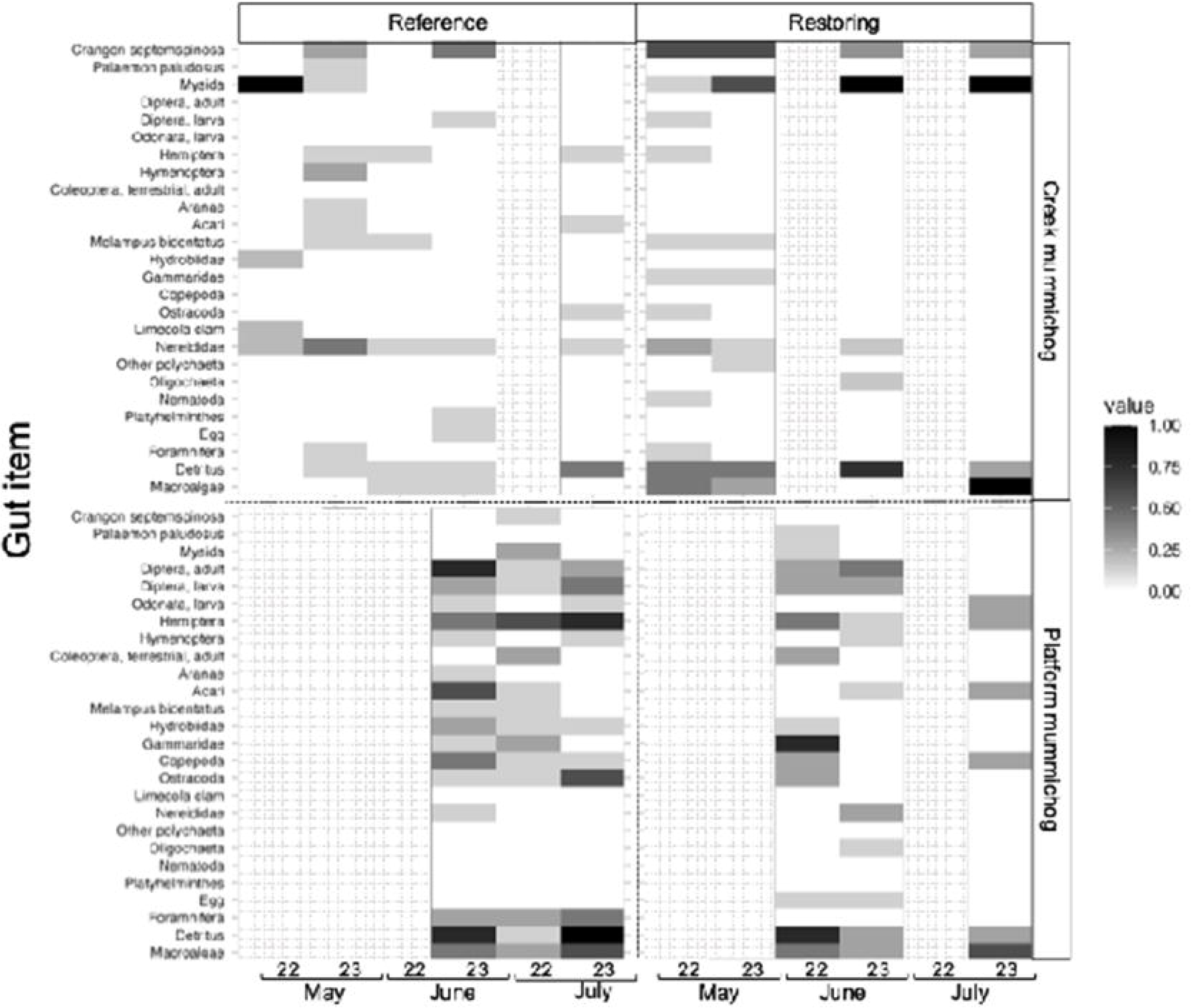
Heatmap showing average presence-absence of gut items in mummichogs (*Fundulus heteroclitus*) captured in creeks and on platforms in Wallace Bay reference and restoring salt marshes for three months (May–July) in 2022–2023. A value of 1 (black) represents consistently present, 0 (white) consistently absent, and shades of grey are in between. Averaged over fish captured in the ebb fyke net (creeks) and in seine hauls per sampling round per site type (*n* = 3–6, Table S4.1). An “A” or an “L” following a taxa name indicates an adult or larval life stage. Areas with gridlines represent rounds when no fish were captured. The order of prey listed on a heat map reflects nekton prey first, then invertebrate prey typical of salt marshes followed by those typical of mudflats, and then other prey items.

For marsh platforms, mummichog gut fullness did not differ significantly, averaging 4.8 ± 0.3% for ingested food relative to body weight (PERMANOVA, Table 4, Fig. 6). However, prey composition for platform mummichogs differed by site type depending on round, due to differences in centroids and dispersion (PERMANOVA and PERMDISP, Table 4, Fig. 8); differences occurred in July of both years (Pairwise comparisons, Table S4.6). Prey taxa were quite diverse; the main ones contributing to the large compositional dissimilarity (87%) between site types were vegetal detritus, insects (hemipterans, dipterans), hydrobiid snails, and various very small organisms (foraminifera, mites, and ostracods) being more consistently present in reference platform mummichogs, while macroalgae, copepods, and polychaetes (nereids, spionids and capitellids) being more so in restoring platform mummichogs (SIMPER, Table S6.2).

#### 3.5.4. Spatiotemporal variation in mummichog gut fullness and prey assemblage in Wallace Bay

The Site type effect (main effect and its interaction with Round) in Wallace Bay accounted for more of the gut fullness variation (49% vs 1%) and prey assemblage variation (36-44% vs 12-21%) for creek mummichogs than platform mummichogs (Tables 4 and S4.5). In comparison, temporal variation (Round alone) was lower for creek mummichogs than platform mummichogs: gut fullness (11% vs 32%) and prey assemblage that included vegetal matter (1% vs 7%) (Table 4). Prey assemblage without vegetal matter had similar temporal variation (13%) for creek and platform mummichogs (Table S4.5). For all analyses, individual fish accounted for a sizeable amount of variation (40-75%).

## 4. DISCUSSION

We measured multiple responses of nekton to salt marsh restoration in two very different case studies. The Aulac project is in a megatidal environment with its restoring site, a former pastureland, being over a decade old, whereas the Wallace Bay project is in a microtidal environment with its restoring site, previously a freshwater impoundment, being a couple of years old. These large situational differences provide insight into what nekton restoring dynamics can be. Any comparison between projects must be done cautiously because we cannot directly address the influence of tidal regime, pre-restoration state, or age. In both projects, we found that visiting nekton species were similar between reference and restoring sites, but community structure differed between these site types; salt marsh pools provided nekton support, but this support appeared different between site types (significantly different environmental conditions, potential prey communities, and nekton communities); and fish gut fullness was similar, but prey composition differed between site types. Dissimilarities in composition for taxa-related measures ranged from 43-100% between site types.

Our application of a multifaceted approach to monitoring a specific ecosystem function (i.e., nekton support) following restoration allows for a holistic understanding of its response. Our approach included structural and functional metrics that together highlighted nuances in restoration progress through the lens of nekton. The visiting nekton community is a structural metric that captures the diversity of species that use salt marshes to varying degrees. Accounting for life stage of the visiting nekton begins to give insights into the habitat support being provided to each species. Being isolated at low tide, the environmental conditions and faunal communities in salt pools provide an understanding of the quality and functional support that is available to nekton when other areas of the marsh are inaccessible. Foraging dynamics by fishes demonstrate whether an ecological function is being fulfilled and to what degree by a restoring site. The insights gained by close examination of each of our metrics is discussed below along with their hypotheses for future research.

### 4.1. The visiting nekton community in creeks and on platforms in two distinct salt marsh restoration case studies

The visiting nekton communities in reference and restoring marshes differed considerably (14 52% of the variation) in both our older megatidal (Aulac) and young microtidal (Wallace Bay) projects. For Aulac, although species richness and total nekton numbers in creeks were similar, reference creeks supported higher numbers of various sticklebacks and silversides, while juvenile and adult eels, mummichogs, tomcods, and sand shrimps were more abundant in restoring creeks. For the Aulac reference platform, species richness and total nekton numbers were generally higher compared to the restoring platform, but the array of species underlying site type variation was similar to that in creeks. For Wallace Bay, species richness was higher in the restoring site (creeks and platform) than in the reference site; the restoring site supported various fish and invertebrates (e.g., sand and grass shrimps, silversides, white perch, and white tipped mud crabs), while the reference site was dominated by juvenile and adult mummichogs. Variations in the visiting nekton communities between reference and restoring marshes may be attributed to differences in platform function between site types in both Aulac and Wallace Bay.

Estuarine nekton can rapidly access new areas made available by salt marsh restoration, given their high mobility and resilience (Raposa 2002, Able et al. 2004, Lechene et al. 2018). Our results in the young Wallace Bay restoration and nekton monitoring in the first years of the Aulac restoration (Brylinsky 2012) support this rapid response. Our assessment of Aulac over a decade later, shows similar species between site types, but still a difference in community structure, highlighting the time it may take for full recovery. Comparisons of nekton communities between reference and restoring marshes have yielded conflicting results with some identifying higher abundances and species richness in reference marshes (Able et al. 2000, Able et al. 2008, Raposa 2008), while others report the opposite (Roman et al. 2002, Hampel et al. 2003). Yet others have found nekton abundances and assemblages to be similar between reference and restoring marshes (Burdick et al. 1997, Raposa & Talley 2012). These studies occurred in various regions, assessed recovering marshes at varying ages, and involved different restoration techniques and sampling strategies. Therefore, the inconsistent patterns in the literature are not surprising. Furthermore, there is a lack of long-term monitoring data (more than a decade) comparing reference and restoring salt marsh nekton, resulting in many unknowns surrounding whether it is feasible to expect that restoring communities will eventually mirror those in reference sites (Raposa & Talley 2012). Our Aulac project indicates that it can take more than a decade for a megatidal environment. In the following, we explore site-specific aspects that may have contributed to site type differences in creek and platform nekton communities.

#### 4.1.1. Potential contributors to variation in nekton community composition between reference and restoring marshes

Hydrology and platform elevation are key aspects in salt marsh restoration trajectories as they can facilitate colonization and reestablishment of natural plant and faunal communities (Able et al. 2008, Bowron et al. 2011). Both aspects regulate nekton access to the marsh platform, a subhabitat that is particularly important for resident fish species (McIvor & Odum 1988, Kneib 1997). In some cases, resident species make up a larger proportion of the salt marsh nekton community in reference marshes compared to those undergoing restoration (Kimball & Able 2007a). While we observed this in Wallace Bay with mummichogs being dominant in reference creeks and on platforms, we observed the reverse in Aulac where mummichogs were more common in restoring creeks and platforms.

Coinciding with the high tidal range, Bay of Fundy salt marshes occur at a high elevation relative to mean sea level, reducing platform access to nekton (Bateman 2021, Endresz et al. 2026). For many decades (since the 1950s), the large agricultural dike completely eliminated tidal flow and associated sedimentation from the Aulac restoring site resulting in its platform being considerably lower in elevation than the reference platform (Virgin et al. 2020, Rudin 2021). At the time of sampling (∼2022), the restoring platform was on average ∼94 cm lower than the reference platform and colonized by *S. alterniflora,* creating low marsh habitat, which is more accessible to nekton. In contrast, the reference platform in Aulac has a large high marsh zone (dominated by *S. patens*), which is flooded infrequently and thus less accessible to nekton, and low marsh zones restricted to the edges of creeks and pools. Higher numbers of mummichogs, tomcods, eels, and sand shrimps could be indicative of the value of the accessible low marsh habitat currently being provided by the restoring marsh in Aulac. The higher numbers of sticklebacks, silversides, and juvenile smelts in the reference marsh suggests the importance of access to a vegetated high marsh platform that provides rich food resources and refuge at high tide for some species. In an upper Bay of Fundy salt marsh restoration involving a culvert expansion, Bowron et al. (2011) also recorded higher numbers of mummichogs and eels in the restoring site and more silversides in the reference site. This study revealed some differences to ours (e.g., more sticklebacks in the restoring site); however, such variations may be associated with differences in age, location, tidal range, and pre-restoration state.

In Wallace Bay, at approximately the time of sampling, the restoring platform was at a lower elevation (on average ∼25 cm lower) than the reference platform, and functioning entirely as low marsh habitat. Unlike in Aulac, this elevational difference is likely not contributing as greatly to nekton community differences due to the microtidal regime of the Northumberland Strait. Salt marshes in this region are at a lower elevation relative to mean sea level, and more equally partitioned into low marsh and high marsh zones, reflecting more frequent tidal inundation. Thus, the reference site in Wallace Bay supports a considerable low marsh area that is available to nekton. Mummichogs dominate the Wallace Bay reference site (86-94% of nekton individuals, Table 1), but not the restoring site (12-14%). Rather, various crustaceans (e.g., sand and grass shrimps, mud crabs), white perch (in creeks) and silversides (on the platform) made up a larger proportion of the restoring community. Given the young age of the Wallace Bay restoration, the restoring site was undergoing a transition that results in an abundance of decaying plant matter as the freshwater vegetation dies off from tidal inundation (Smith & Warren 2012). During this transitionary stage, benthic algal production rapidly increases on the platform (Zheng et al. 2004, Able et al. 2008). At the time of sampling, the restoring platform was composed of bare areas covered with benthic green algae, dotted with patches of *S. alterniflora,* and surrounded by dying freshwater plants (*Typha* cattails) at the edge of the site (Dickinson 2024). The abundance of decaying plant matter and increased levels of benthic algal production may be drawing shrimp into the restoring site, as these represent important dietary sources (Robertson & Weis 2007).

In addition, the lack of a densely vegetated marsh platform in the Wallace Bay restoring site may have led to mummichog and other small nekton being more vulnerable to predation, potentially increasing white perch presence. White perch are known to consume mummichogs and small crustaceans including shrimps (Weis 2005). Moreover, the high numbers of mummichogs in the reference site may be keeping crustacean numbers lower as these fish consume small crustaceans including shrimps and juvenile crabs (Dibble et al. 2014, this study) and have demonstrated top-down control on shrimp populations in other geographic locations (Kneib 1986).

Our examination of the visiting nekton community allowed us to identify the species present in and potentially using our marshes. In summary, similar species were observed in reference and restoring sites in both our projects supporting a rapid response by nekton to salt marsh restoration. However, differences in community structure may reflect variations in subhabitat function between site types. In our decade-old Aulac restoration, a considerable elevational difference still exists between site types and could be indicative of a slower recovery trajectory in megatidal estuaries. Nekton community variations between reference and restoring sites in Aulac may reflect the importance of low vs. high marsh habitat for certain species. In our young Wallace Bay project, elevational differences between site types are relatively small and nekton community variations are likely attributed to the transitionary stage of the restoring site. The lack of a densely vegetated marsh platform in the Wallace Bay restoring site appears to support less suitable habitat for the resident mummichog. However, it may provide enhanced feeding opportunities for crustaceans and transient fish (e.g., white perch). The visiting nekton community represented our structural metric for assessing salt marsh restoration success and provides complementary insights gained from our two functional metrics (below).

### 4.2. Pool recovery dynamics in two distinct salt marsh restoration case studies

Pool environmental conditions (9 35% of the abiotic variation, 63-67% of the aquatic vegetation variation) and faunal communities (18 72% of the variation) differed between reference and restoring sites in Aulac and Wallace Bay, indicating possible differences in function. In Aulac, differential site type patterns are likely attributed to our comparison of somewhat different subhabitats: true salt pools versus creek pools. The development of true salt pools in a megatidal salt marsh like Aulac may only begin once high marsh conditions are established in the restoring site (Virgin et al. 2020). Salt pools have been found in higher densities in high marsh zones than in low marsh zones in mesotidal salt marshes (e.g., Plum Island, Massachusetts; Millette et al. 2010, Wilson et al. 2014). Based on our cursory observations for over a decade, we suspect that any area that is depressed in the Aulac restoring site gets filled in through sedimentation, delaying salt pool development. Sediment deposition can be very high in regularly inundated areas, since the upper Bay of Fundy tidal waters carry a very high suspended sediment load (0.12–12.67 g/L; Amos & Tee 1989, van Proosdij et al. 2006).

The frequency of tidal flooding leads to inherent abiotic differences between salt pools and pools within the creek system. Creek pools are tidally flooded more frequently and so are more connected to the surrounding estuarine environment than salt pools; consequently, creek pool water conditions tend to be similar to tidal waters (Allen et al. 2017, Kimball et al. 2023). Salt pools are flooded less frequently and typically experience higher salinity and temperature fluctuations (Adamowicz & Roman 2005). Our observations in Aulac supported the above: restoring creek pools generally had water conditions (salinity, pH, dissolved oxygen concentration) more similar to the surrounding Bay waters as well as firmer bottom sediment (indicative of relatively high water flow rates) (Electronic Supplement 8). In comparison, Aulac reference salt pools had more dissimilar water conditions, as well as softer bottom sediment. We observed a different pattern in the microtidal Wallace Bay likely because the restoring pools were located on the marsh platform and resembled early-stage salt pools (Fig. S3.2) with shallow depths and unstable bottom sediment. Thus, here, the variations in abiotic conditions between reference and restoring pools were likely influenced by site type (rather than being different subhabitats). Reference pools in Wallace Bay were deeper, and had higher pH levels (Koop-Jakobsen & Gutbrod 2019) and firmer bottom sediment than restoring pools.

The biotic communities in pools complemented our abiotic measures, and also showed potential differences in habitat provisioning between reference and restoring pools. Salt pools support the growth of submerged aquatic vegetation like ditch grass (*R. maritima*) and macroalgae providing aquatic invertebrates and small-bodied fish with food and refuge (Spivak et al. 2017, Litvin et al. 2018). In Aulac and Wallace Bay, summer aquatic vegetation cover was considerable in reference salt pools and minimal in creek pools and early-stage salt pools in restoring sites. We suspect that lack of this feature in restoring pools in both projects contribute to site type differences in faunal communities (Virgin et al. 2020). Invertebrates also contribute to the provisioning status of pools for fish. In both projects, the reference site had the usual salt pool taxa, dominated by hydrobiid snails and gammarid amphipods (Virgin et al. 2020, Noel et al. 2023), whereas the restoring site pools had mud whelks (captured in minnow traps). The content of the invertebrate activity traps in restoring pools aligned with our abiotic observations; these traps were almost empty in Aulac creek pools, and contained low densities of sand shrimp, gammarid amphipods, and hydrobiid snails in the early-stage salt pools in Wallace Bay. The pool community of fish in our reference sites was similar to that of other established salt marshes (Nixon & Oviatt 1973, Poulin & FitzGerald 1989, Endresz et al. 2026), with mummichogs and sticklebacks dominating. Although restoring pools also had mummichogs and sticklebacks, transient species and invertebrate nekton were present, including juvenile tomcod, American eels, and sand shrimp in Aulac, and sand and grass shrimp, mud crabs, and green crabs in Wallace Bay. Our faunal observations indicate differences in the habitat provisioning of pools in the restoring sites compared to the reference sites.

The presence of juvenile mummichogs in restoring pools does not necessarily indicate that these pools are serving the same nursery function as reference pools. In both our projects, we only captured very small juvenile mummichogs and sticklebacks (< 3 cm length, in invertebrate activity traps) in reference salt pools, whereas large juvenile mummichogs (3-5 cm length, in minnow traps) were captured in greater numbers in restoring pools than reference salt pools. Minnow and invertebrate activity traps are size selective, and the community captured in the latter is likely a better reflection of nursery function. Nursery habitats support higher densities, and growth and survival rates of juvenile nekton (Beck et al. 2001). Our reference salt pools appear to meet these nursery criteria particularly for mummichogs and sticklebacks. They generally had higher summer temperatures and abundance of food (i.e., algae, invertebrate taxa), both of which contribute metabolic advantages to juvenile nekton (Spivak et al. 2017, Campana & Hurley 1989). Ditch grass and macroalgae offer enhanced refuge while providing food to the invertebrates that are preyed on by juvenile fish. Our restoring pools do not appear to offer the same metabolic support and protection provided by reference salt pools, likely indicating reduced nursery function.

In summary, assessing the abiotic conditions and vegetative, invertebrate and fish communities in pools allowed us to look at the quality and function of the essential salt pool habitat. A primary function of salt pools is to provide low tide refuge, which given the presence of nekton in reference and restoring pools, appears to be fulfilled in both projects. The differences in pool abiotic conditions between site types reflect variation in the degree of isolation from tidal flooding. This is most pronounced in Aulac where the restoring site pools are within the intertidal creek network and are tidally flooded consistently, whereas the reference pools can be isolated for up to a few weeks. The high frequency of tidal flooding that creek pools experience limits other habitat support that is provided by salt pools. Salt pools can also support rich feeding and nursery grounds for certain nekton species; based on our examination of the invertebrate activity trap contents and vegetation cover, this enhanced habitat support only seems to be present in reference pools. In both projects, salt pool development appears to take time and may occur alongside the establishment of the high marsh zone. The state of salt pools represented a functional metric in our assessment of salt marsh restoration success, and we observed that restoration pools have yet to reach the heightened habitat support of reference salt pools.

### 4.3. Foraging patterns of dominant fishes in two distinct salt marsh restoration case studies

A key nekton function of salt marshes is foraging support (Rountree & Able 2007), which based on gut fullness indices, appears to be met in our Aulac and Wallace Bay projects. Average gut fullness values for Site type-Subhabitat-Round combinations ranged from 3% to 9.5% of the intact fish weight for mummichogs and tomcods, and did not differ significantly by site type, except for mummichogs in Wallace Bay creeks (where in July restoring fish had relatively high fullness values, ranging from ∼5-13%, Fig. 6). Other studies that evaluated dietary support of restoring salt marshes also observed similar gut fullness between fish captured in reference and restoring marshes (Kimball & Able 2007b, Stamp et al. 2022). Together, this suggests that foraging support at a basic level returns with relative ease in salt marshes undergoing restoration. However, assessing gut item assemblages provides enhanced understanding of this ecological function in restoring salt marshes.

Tomcods and mummichogs are omnivorous and opportunistic feeders; therefore, their diets tend to be highly diverse and largely influenced by the availability of food resources (Kneib 1986, Howe 1971). In both our projects, for all species-subhabitat combinations, the most variation in prey assemblage (40-88%) occurred between individual fish, reflecting the flexible nature of tomcod and mummichog food selection. The prey assemblage in both species was primarily composed of a variety of faunal taxa including polychaetes, crustaceans (amphipods, shrimp, copepods, ostracods), insects, and nematodes which is consistent with other studies (Kneib & Stiven 1978, Joyce & Weisberg 1986, Carroll et al. 2024). For example, in the Hudson River estuary (New York and New Jersey), tomcods fed on amphipods, mysids, isopods, sand shrimps, small molluscs, squids, and fish larvae (Grabe 1978, Grabe 1980). In Montsweag Bay, (Maine), polychaetes were of primary importance in tomcod diets (Alexander 1971). In a region of the Bay of Fundy different than our study (Avon River estuary in Minas Basin), *C. volutator* amphipods and sand shrimps dominated tomcod gut contents (Carroll et al. 2024). In Atlantic USA salt marshes, mummichogs fed on small crustaceans, insects, annelids, gastropods, nematodes, and detritus (Weisberg et al. 1981, James-Pirri et al. 2001). Detritus and macroalgae were recurring gut items in tomcods and mummichogs in Aulac and Wallace Bay. These vegetal items are thought to offer little nutritional value to the fish, and their consumption may occur incidentally while foraging for fauna (Prinslow et al. 1974, Kneib et al. 1980).

The prey assemblage of tomcods and mummichogs captured in creeks and of mummichogs on platforms differed by site type (12-44% of the variation) in both our case studies. In Aulac, although tomcods captured in both reference and restoring sites predominantly consumed *C. volutator* amphipods and nereid polychaetes, reference tomcods consumed salt marsh-related items such as gammarids, juvenile sticklebacks, hemipterans and detritus more consistently. Similarly, Aulac mummichogs, independent of site type and subhabitats, consumed fair amounts of nereid polychaetes, gammarid and corophiid amphipods, insects, and detritus.

Main contributors to site type diet differences in mummichogs were more feeding on nereids, gammarids, hemipterans and detritus in reference creeks, and nereids and certain adult dipteran taxa in reference platforms compared to the restoration site. Gut items for restoration site mummichogs also showed a mixed bag, with more feeding on chironomids, larval flies, and *C. volutator* in restoring creeks, and gammarids, *C. volutator*, hydrobiid snails, hemipterans and detritus on restoring platforms than at the reference site. The prey item assemblage in Aulac reference and restoring tomcods and mummichogs demonstrate strong linkages with the adjacent mudflat ecosystem. Polychaete worms (including Nereididae), *C. volutator* and *Limecola* clams are common mudflat fauna (Gerwing et al. 2015) and appear to be important prey items for tomcods and mummichogs in Aulac.

Mummichogs in Wallace Bay, across site types and subhabitats, generally consumed most commonly nekton crustaceans (mysids, sand shrimps), hemipterans, detritus and green macroalgae. Main contributors to site type diet differences were more nereids in reference creeks and terrestrial arthropods (Hemiptera, Hymenoptera, Diptera, Coleoptera, Acariformes) and snails (Hydrobiidae and *Melampus*) on reference platforms, whereas more feeding on mysids and sand shrimps in restoring creeks, and copepods and polychaetes (including nereids, spionids and capitellids) on restoring platforms Given that restoring creek nekton communities in Wallace Bay supported high numbers of mysids and sand shrimps, it is not surprising that they were a common prey item in restoring site mummichogs.

Differences in tomcod and mummichog prey assemblages between site types may reflect the transitionary stage of restoring sites in both of our projects. In particular, although consumed insects were present in all combinations of fish species, site types and subhabitats, their presence in restoring platform mummichogs in Wallace Bay was low, likely reflecting a very young restoration site with a mainly muddy platform environment. The other sites had various insect taxa contributing appreciably to fish diets, as well as had extensive vegetation, with Wallace Bay reference having both a substantial low zone (*S. alterniflora*) and high zone (*S. patens*), and Aulac reference and restoration dominated by high marsh zone and low marsh zone, respectively. Indeed, recovery of the typical marsh platform invertebrate community (including insects and arachnids) lags behind that of the vegetation community (Craft 2000, Petillon et al. 2014, Virgin et al. 2020). Thus, the gut contents in generalist fish like tomcods and mummichogs likely reflect the invertebrate community present in the reference and restoring marshes, including the recovery trajectory in the latter.

In summary, our assessment of gut fullness and contents in tomcods and mummichogs from Aulac and Wallace Bay give insights into the foraging support provided by salt marshes to nekton. Gut fullness results indicated that the restoring salt marshes in both projects fulfilled the foraging function at a basic level. The omnivorous and opportunistic feeding behaviour of tomcods and mummichogs appears to allow these species to shift their diet. Site type patterns in prey assemblages of tomcods and mummichogs from both projects may reflect differences in prey availability between reference and restoring marshes. Our restoring sites are in transition (being entirely low marsh in Aulac, or transitioning from mudflat to low marsh in Wallace Bay) and are more influenced by tides, whereas our reference sites are mature salt marshes (with a high marsh zone, associated with certain terrestrial arthropod taxa). As with our other metrics, we learned that ecologically, the foraging function for nekton has yet to fully recover; however, there are indications that conditions mirroring reference marshes are not needed to provide valuable habitat to tomcods and mummichogs (Stamp et al. 2022).

## 5. CONCLUSIONS

Understanding the response of species and biotic communities to salt marsh restoration is fundamental to evaluating the effectiveness of these actions and improving outcomes. Measuring response using multiple and integrated approaches provides a well-rounded view. Both our case studies (Aulac and Wallace Bay) showed similar nekton species in reference and restoring sites. However, the community structure (e.g., the dominating species) differed by site type, even in our decade-old Aulac project. The restoring pools offer low tide refuge, but do not support to the same degree other nekton functions (foraging and nursery grounds) provided by reference pools. Restoring marsh platforms and creeks in both projects appear to fulfill foraging function for tomcods and mummichogs, since their gut fullness values were similar between site types, despite differences in gut content assemblages. Thus, the restorations in both projects are providing valuable habitat to nekton and can be viewed as being successful from the point of view of nekton. Our study demonstrates the effectiveness of using metrics like community structure and usage patterns to evaluate the progress of salt marsh restoration projects. It also reinforces the importance of using standardized and consistent methodologies in reference and restoring marshes over multiple years to identify and understand restoration patterns and trajectories. Although we did not directly compare the two projects for obvious reasons, we have provided insights into the potential implications of tidal regime on salt marsh restoration trajectories.

## Acknowledgements

Our study was supported by a MITACS ACCELERATE Doctoral Fellowship (IT30551, to KCE and MAB) and three MITACS ACCELERATE internships (IT25528, IT30590, IT46603) in partnership with Ducks Unlimited Canada; New Brunswick Wildlife Trust Fund (grants # BE23-13, BE22-17, and B201-216 to MAB and KCE); New Brunswick Innovation Fund (Climate Impact Research Fund grant # CIF_2022_011 to MAB); Natural Sciences and Engineering Research Council of Canada (Discovery grant # RGPIN-2020–04106 to MAB); Clean Tech (Environment and Climate Change Canada) and Natural Resources Internships (Natural Resources Canada) administered by the Career Launcher Program; Canada Summer Jobs Program (Employment and Social Development Canada); Future NB via UNB Engineering and Science Co-op Program; and the University of New Brunswick. We thank James Curtis, John Linihan, Romeo Naeder, Greg Norris, Anajose Reyes Guevara, Alexa Stack Mills, and Krish Thapar for field assistance; James Curtis and Hilary MacLean for assistance with fish dissections; Jeff Ollerhead for marsh platform elevation data; Olivier Grimard and Nicole Endresz for assisting with figure execution in R; and John Brazner, Tommi Linnansaari, and Liz Bateman for feedback and insightful comments on all aspects of the project.

