## Supplemental information for "Integrating nekton communities, salt pool dynamics, and fish foraging patterns to assess salt marsh restoration success in Maritime Canada"

Electronic supplementary material for:

Authors: Kiana C. Endresz^1^*, Nic R. McLellan^2^ and Myriam A. Barbeau^1^

Addresses:

^1^Department of Biology, University of New Brunswick, Fredericton, New Brunswick, Canada, E3B 5A3

^2^Ducks Unlimited Canada, 64 Hwy 6, Amherst, Nova Scotia, Canada, B4H 3Z5

Date: 25 May 2026

**Supplement S1: Overall sampling program for salt marsh nekton in Aulac and Wallace Bay, Maritime Canada**

**Table S1** Dates for all nekton and salt pool sampling conducted in the reference and restoring sites for the salt marsh restoration projects in Aulac, NB and Wallace Bay, NS in 2022‒2023. Aulac: 45°51.07’N, 64°17.64’W and 45°51.38’N, 64°18.08’W for the restoring site, and 45°50.69’N, 64°17.24’W for the reference site; Wallace Bay: 45°49.65’N, 63°32.63’W for the restoring site, and 45°50.22’N, 63°33.36’W for the reference site. During each sampling round, we used fyke nets to sample the nekton communities in intertidal creeks over a nighttime high tide; a seine net to sample nekton communities on the marsh platform during a daytime high tide. We sampled salt pools or creek pools using a paired minnow trap and invertebrate activity trap, deployed over a nighttime high tide. We also recorded abiotic measurements (sediment penetrability, and water depth, pH, temperature, salinity, and dissolved oxygen concentration) in the evening at low tide in our selected pools.

| Project | Year | Month | Dates |
| --- | --- | --- | --- |
| Aulac | 2022 | May | 15‒17 |
|  |  | June | 14‒16 |
|  |  | July | 15‒17 |
|  | 2023 | May | 17‒19 |
|  |  | June | 15‒17 |
|  |  | July | 6‒8 |
| Wallace Bay | 2022 | May | 2‒4 |
|  |  | June | 1‒3 |
|  |  | July | 17‒19 |
|  | 2023 | May | 8‒10 |
|  |  | June | 7‒9 |
|  |  | July | 18‒20 |

**Supplement S2: Body size of nekton in Aulac and Wallace Bay, Maritime Canada**

**Table S2.1** Mean (± SD) and range of total body length for fish and shrimp, and carapace width for crabs, in the nekton communities caught in fyke and seine nets in the reference and restoring site in Aulac and Wallace Bay in 2022–2023. n is the number of individuals measured.

|  |  |  | Aulac | | | Wallace Bay | | |
| --- | --- | --- | --- | --- | --- | --- | --- | --- |
| Group | Common name | Scientific name | Mean±SD | Range (cm) | n | Mean±SD | Range | n |
| Fish | Mummichog | *Fundulus heteroclitus* | 7.0 ± 1.6 | 3.6–11.2 | 359 | 7.5 ± 2.2 | 1.4–17.5 | 872 |
|  | Banded killifish | *Fundulus diaphanus* | 6.6 ± 1.4 | 4.6–8.0 | 5 | 7.4 | n/a | 1 |
|  | Atlantic silverside | *Menidia menidia* | 8.0 ± 1.4 | 3.2–11.9 | 314 | 7.4 ± 3.5 | 1.2–11.9 | 202 |
|  | Tomcod | *Microgadus tomcod* | 15.5 ± 3.2 | 3.2–26.1 | 751 | n/a | n/a |  |
|  | Threespine stickleback | *Gasterosteus aculeatus* | 4.9 ± 2.1 | 0.8–8.9 | 543 | 5.3 ± 1.1 | 1.7–7.5 | 129 |
|  | Blackspotted stickleback | *Gasterosteus wheatlandi* | 3.6 ± 0.9 | 1.0–4.9 | 373 | 4.4 ± 0.6 | 3.5–5.3 | 107 |
|  | Fourspine stickleback | *Apeltes quadracus* | 3.5 | n/a | 1 | 3.7 ± 0.7 | 2.2–7.6 | 146 |
|  | Ninespine stickleback | *Pungitius pungitius* | 2.8 ± 0.8 | 1.0–6.2 | 243 | 3.6 ± 0.9 | 2.5–6.2 | 54 |
|  | Striped bass | *Morone saxatilis* | 35.4 | n/a | 1 | 25.0 ± 6.0 | 13.7–33.5 | 33 |
|  | White perch | *Morone americana* | n/a | n/a | 0 | 20.7 ± 3.1 | 10.4–33.8 | 447 |
|  | Winter flounder | *Pseudopleuronectes americanus* | n/a | n/a | 0 | 9.2 ± 2.9 | 6.3–17.5 | 23 |
|  | American eel | *Anguilla rostrata* | 24.7 ± 11.2 | 9.2–65.0 | 205 | 47.5 | n/a | 1 |
|  | Rock gunnel | *Pholis gunnellus* | n/a | n/a | 0 | 11.2 ± 1.4 | 10.1–13.2 | 5 |
|  | Atlantic herring | *Clupea harengus* | n/a | n/a | 0 | 3.4 ± 1.6 | 2.2–4.5 | 2 |
|  | Gaspereau | *Alosa pseudoharengus, A. aestivalis* | 11.7 ± 9.9 | 4.1–26.2 | 11 | 14.2 ± 11.7 | 3.4–28.9 | 18 |
|  | Brook trout | *Salvelinus fontinalis* | 17.4 | n/a | 1 | 21.8 | n/a | 1 |
|  | Atlantic salmon | *Salmo salar* | n/a | n/a | 0 | 8.4 | n/a | 1 |
|  | Rainbow smelt | *Osmerus mordax* | 7.1 ± 1.0 | 4.4–9.8 | 70 | 10.4 ± 3.2 | 3.9–19.2 | 107 |
|  | Northern pipefish | *Syngnathus fuscus* | 15.1 | n/a | 1 | n/a | n/a | 0 |
| Invertebrate | Sand shrimp | *Crangon septemspinosa* | 5.4 ± 0.8 | 3.6–8.1 | 61 | 3.6 ± 0.9 | 1.1–7.0 | 358 |
|  | Grass shrimp | *Palaemon paludosus* | n/a | n/a | 0 | 3.2 ± 0.6 | 0.8–5.2 | 131 |
|  | Green crab | *Carcinus maenas* | 6.7 | n/a | 1 | 5.1 ± 1.5 | 1.2–8.9 | 38 |
|  | Rock crab | *Cancer irroratus* | n/a | n/a | 0 | 7.8 ± 1.1 | 6.5–9.0 | 5 |
|  | Black tipped mud crab | *Panopeus herbstii* | n/a | n/a | 0 | 2.3 ± 0.3 | 2.0–2.5 | 3 |
|  | White tipped mud crab | *Rhithropanopeus harrisi* | n/a | n/a | 0 | 1.9 ± 0.6 | 1.0–4.5 | 51 |

**Table S2.2** Lengths used to differentiate juvenile from adult fishes that were captured in fyke nets, seine nets, and minnow traps in reference and restoring salt marshes in Aulac and Wallace Bay in 2022–2023. Fish captured in invertebrate activity traps were all small juveniles.

| Species | Length of juveniles | Reference |
| --- | --- | --- |
| Mummichog | < 5 cm | Able 1990 |
| Tomcod | < 13 cm | Grabe 1980 |
| Threespine stickleback | < 3 cm | Coad & Power 1973 |
| Blackspotted stickleback | < 3 cm | Poulin & FitzGerald 1987 |
| Ninespine stickleback | < 4 cm | Poulin & FitzGerald 1987 |
| Rainbow smelt | < 12 cm | Collette & Klein-Macphee 2002 |
| Atlantic silverside | < 5 cm | Bengtson 1984 |
| American eel | < 15 cm | Hildebrand & Schroeder 1928 |
| Winter flounder | < 20 cm | Perlmutter 1947 |
| Gaspereau | < 20 cm | DFO 2022 |
| Atlantic herring | < 25 cm | O’Brien et al. 1993 |

**Supplement 3. Photographs of salt pools, creek pools, and invertebrate activity traps in Aulac and Wallace Bay, Maritime Canada**

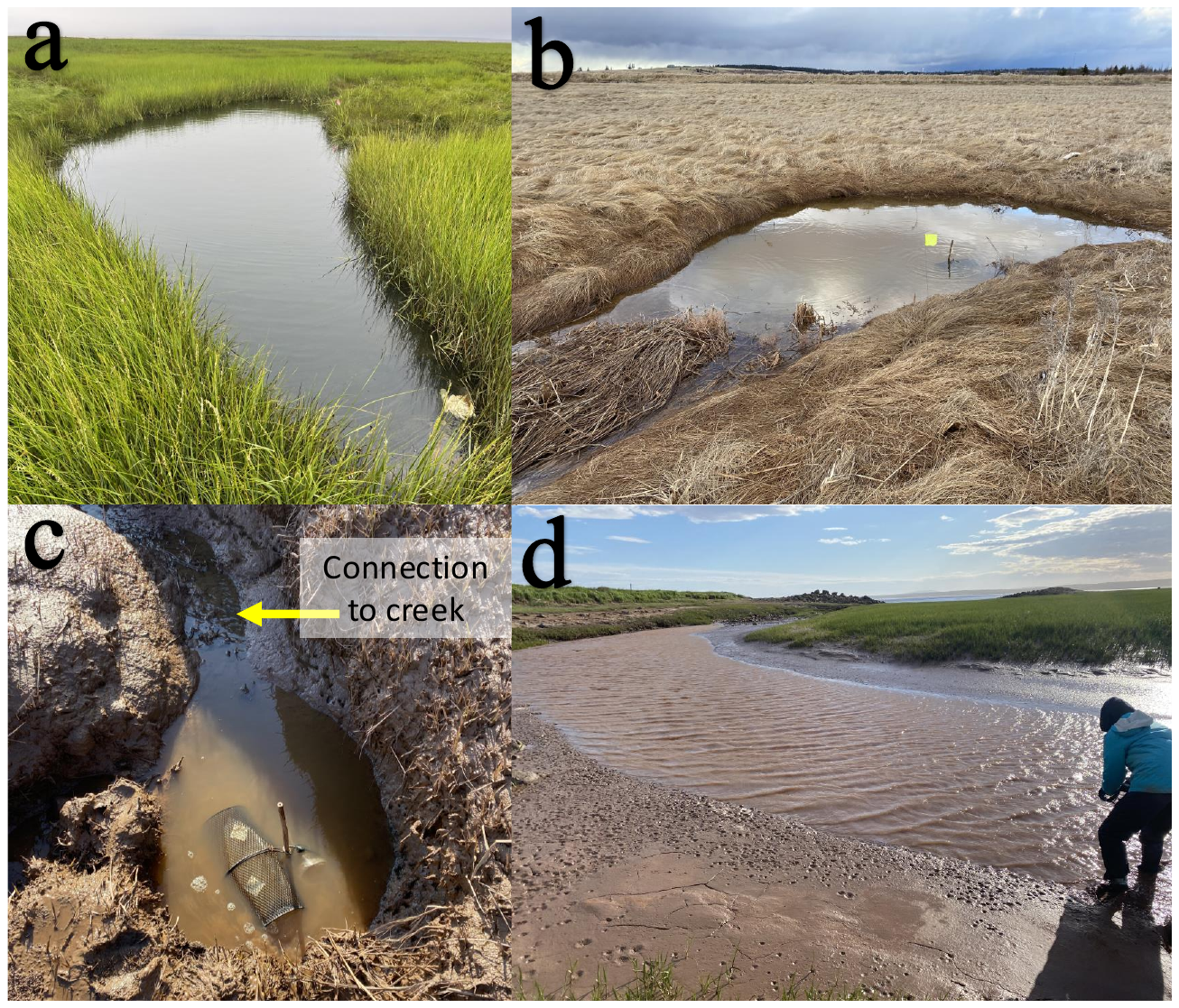

**Fig S3.1.** Photographs of the reference salt pools (a,b) and the restoring creek pools (c,d) sampled in Aulac in 2022‒2023. a) Reference pool in July 2023, b) reference pool in May 2022, c) restoring creek pool with minnow and invertebrate activity trap deployed in May 2022, and d) restoring creek pool in June 2022. Photographs by K.C. Endresz.

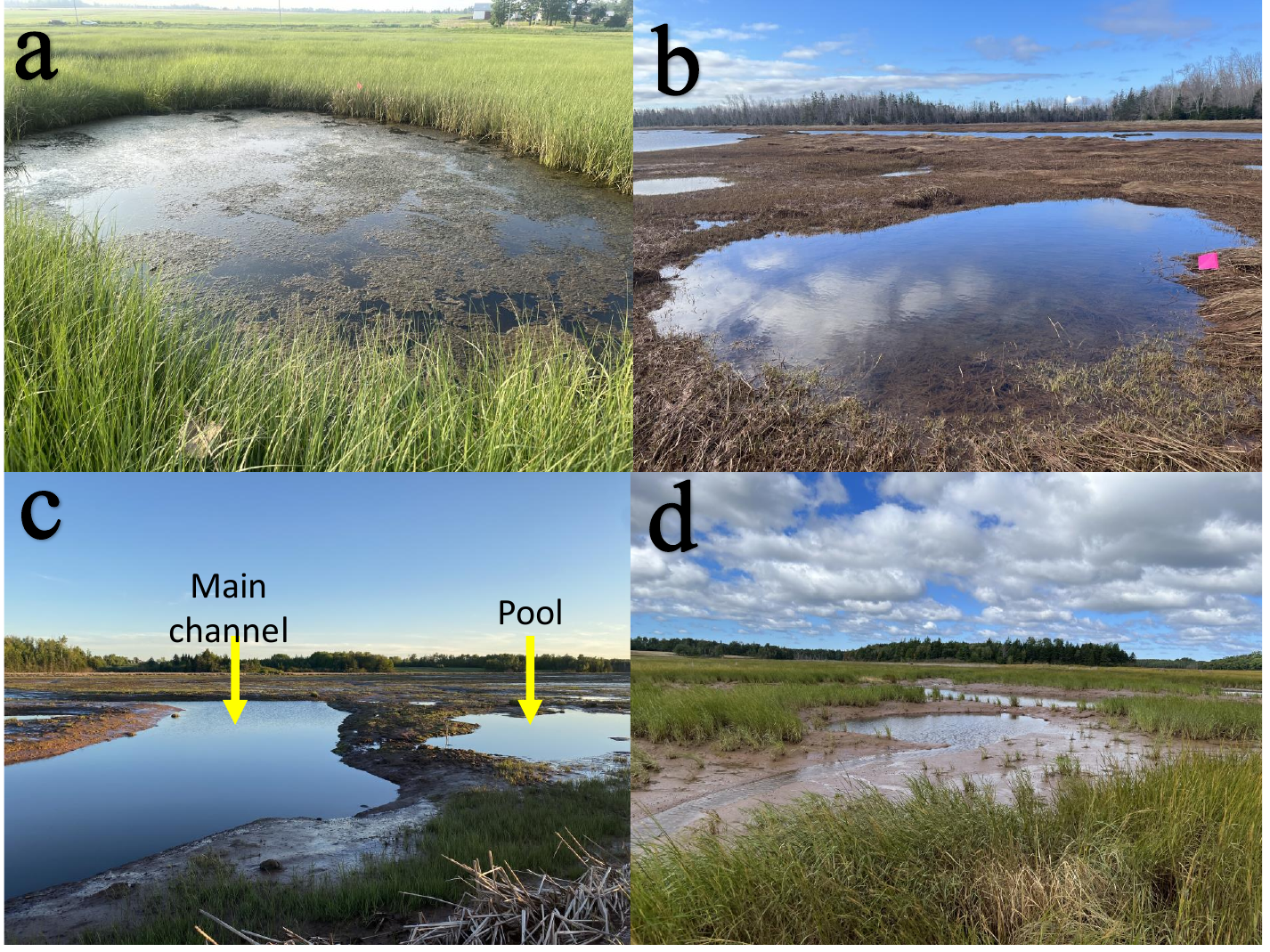

**Fig S3.2.** Photographs of the reference (a,b) and restoring pools (c,d) sampled in Wallace Bay in 2022‒2023. a) reference salt pool in July 2023, b) reference salt pool in May 2023, c) restoring pool with location of the main channel indicated in June 2022, and d) restoring pool in July 2023. Photographs by K.C. Endresz.

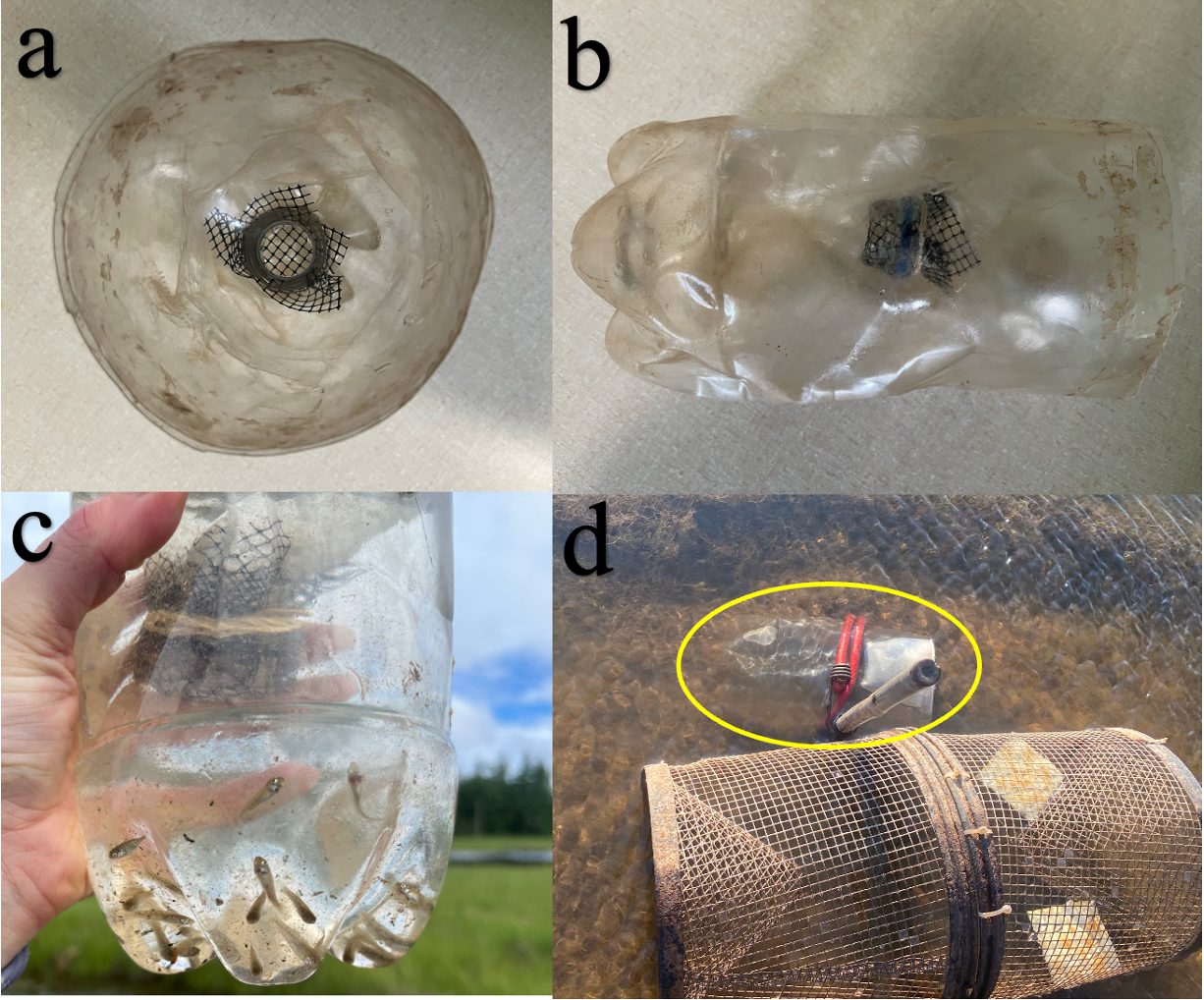
**Fig S3.3.** Photographs of the invertebrate activity trap used to sample faunal communities in reference and restoring salt marsh pools in Aulac and Wallace Bay in 2022‒2023. a) View of the mouth of the trap with mesh secured around the opening, b) side view of the trap, c) juvenile mummichogs and sticklebacks captured in the trap in a reference marsh in July 2023, and d) an invertebrate activity trap (indicated by the yellow oval) attached to a bamboo stake and paired with a minnow trap about to be deployed in a reference salt pool in June 2023. Photographs by K.C. Endresz.

**Supplement 4. Additional information and results of gut content analyses in Aulac and Wallace, Maritime Canada**

**Table S4.1** Number of tomcods (*Microgadus tomcod)* and mummichogs (*Fundulus heteroclitus*) retained in fyke nets (facing the ebb tide) and seine hauls from Aulac and Wallace Bay reference and restoring salt marshes for gut content analysis in 2022‒2023. No tomcods were captured in Wallace Bay**.** See Table S1 for dates of sampling rounds.

| Project | Site type | Year | Month | Net type | Species | Number |
| --- | --- | --- | --- | --- | --- | --- |
| Aulac | Reference | 2022 | May | Fyke net | Tomcod | 6 |
|  |  |  |  | Fyke net | Mummichog | 6 |
|  |  |  | June | Fyke net | Tomcod | 6 |
|  |  |  |  | Seine net | Mummichog | 6 |
|  |  |  | July | Fyke net | Tomcod | 10 |
|  |  | 2023 | May | Fyke net | Tomcod | 6 |
|  |  |  |  | Fyke net | Mummichog | 5 |
|  |  |  | June | Fyke net | Tomcod | 6 |
|  |  |  |  | Fyke net | Mummichog | 6 |
|  |  |  |  | Seine net | Mummichog | 6 |
|  |  |  | July | Fyke net | Tomcod | 7 |
|  |  |  |  | Fyke net | Mummichog | 5 |
|  |  |  |  | Seine net | Mummichog | 1 |
|  | Restoring | 2022 | May | Fyke net | Tomcod | 6 |
|  |  |  |  | Fyke net | Mummichog | 4 |
|  |  |  | June | Fyke net | Tomcod | 10 |
|  |  |  |  | Seine net | Mummichog | 6 |
|  |  |  | July | Fyke net | Tomcod | 6 |
|  |  | 2023 | May | Fyke net | Tomcod | 8 |
|  |  |  |  | Fyke net | Mummichog | 8 |
|  |  |  | June | Fyke net | Tomcod | 6 |
|  |  |  |  | Fyke net | Mummichog | 6 |
|  |  |  |  | Seine net | Mummichog | 6 |
|  |  |  | July | Fyke net | Tomcod | 6 |
|  |  |  |  | Fyke net | Mummichog | 5 |
| Wallace Bay | Reference | 2022 | May | Fyke net | Mummichog | 4 |
|  |  | 2023 |  | Fyke net | Mummichog | 6 |
|  |  | 2023 | June | Fyke net | Mummichog | 6 |
|  |  |  |  | Seine net | Mummichog | 6 |
|  |  | 2022 | July | Seine net | Mummichog | 6 |
|  |  | 2023 |  | Fyke net | Mummichog | 6 |
|  |  |  |  | Seine net | Mummichog | 6 |
|  | Restoring | 2022 | May | Fyke net | Mummichog | 6 |
|  |  | 2023 |  | Fyke net | Mummichog | 6 |
|  |  | 2023 | June | Fyke net | Mummichog | 6 |
|  |  |  |  | Seine net | Mummichog | 6 |
|  |  | 2022 | July | Seine net | Mummichog | 6 |
|  |  | 2023 |  | Fyke net | Mummichog | 3 |
|  |  |  |  | Seine net | Mummichog | 3 |

**Table S4.2** Reference table of prey taxa observed in gut contents of tomcods (*Microgadus tomcod)* and mummichogs (*Fundulus heteroclitus*), and the corresponding coarse taxonomic categories used in subsequent dietary analyses. The tomcods and mummichogs were captured in creeks and on platforms in reference and restoring salt marshes in Aulac and Wallace Bay in May–July in 2022–2023.

| **Group of gut items** | **Taxa** | **Level of Identification** | **Taxonomic names used** |
| --- | --- | --- | --- |
| Fish | Atlantic silverside | Species | *Menidia menidia* |
|  | Mummichog | Species | *Fundulus heteroclitus* |
|  | Stickleback | Family | Gasterosteidae |
| Crustacea, Malacostraca | Green crab | Species | *Carcinus maenas* |
|  | Grass shrimp | Species | *Palaemon paludosus* |
|  | Sand shrimp | Species | *Crangon septemspinosa* |
|  | Mysida | Order | Mysida |
|  | Amphipoda | Species, Family | *Corophium volutator,* Gammaridae |
|  | Isopoda | Order | Isopoda |
| Crustacea, Other | Copepoda | Subclass | Copepoda |
|  | Ostracoda | Class | Ostracoda |
| Insecta | Coleoptera | Family, Order | Corixidae, other Coleoptera |
|  | Diptera | Family, Order | Chironomidae, other Diptera |
|  | Hemiptera | Order | Hemiptera |
|  | Hymenoptera | Order | Hymenoptera |
|  | Odonata | Order | Odonata |
| Arachnida | Acariformes | Superorder | Acariformes |
|  | Araneae | Order | Araneae |
| Mollusca | Gastropoda | Species, Family | *Melampus bidentatus*, *Tritia (Ilyanassa) obsoleta*, Hydrobiidae, |
|  | Bivalvia | Genus | *Limecola* |
| Annelida | Polychaeta | Family, Class | Nereididae, other Polychaeta |
|  | Oligochaeta | Subclass | Oligochaeta |
| Other, worms | Nematoda | Phylum | Nematoda |
|  | Platyhelminthes | Class | Trematoda |
| Other, Rhizaria | Foraminifera | Class | Foraminifera |

**Table S4.3** Means ± SE total body length of tomcod (*Microgadus tomcod)* and mummichogs (*Fundulus heteroclitus*) retained for gut content analysis, in reference and restoring creeks and platforms in May‒July 2022‒2023 in Aulac and Wallace Bay. Tomcods were only captured in reference and restoring creeks in Aulac, while mummichogs were captured in creeks and on platforms in both site types in Aulac and Wallace Bay. See Table S4.1 for the number of each species caught and retained, and Table S1 for dates of sampling rounds.

| Project | Species | Subhabitat | Site type | Mean length ± SE (cm) |
| --- | --- | --- | --- | --- |
| Aulac | Tomcod | Creek | Reference | 14.6±0.6 |
|  |  |  | Restoring | 15.1±0.3 |
|  | Mummichog | Creek | Reference | 7.9±0.3 |
|  |  |  | Restoring | 7.8±0.3 |
|  |  | Platform | Reference | 6.7±0.3 |
|  |  |  | Restoring | 7.8±0.4 |
| Wallace Bay | Mummichog | Creek | Reference | 8.0±0.3 |
|  |  |  | Restoring | 8.4±0.3 |
|  |  | Platform | Reference | 7.0±0.2 |
|  |  |  | Restoring | 7.3±0.3 |

**Table S4.4** Permutational ANOVA results for total length (0.1 cm accuracy) for fish used in gut content analyses, as well as for percent vegetal matter in gut contents of tomcods (*Microgadus tomcod*) and mummichogs (*Fundulus heteroclitus*) in Aulac and Wallace Bay reference and restoring creeks and platforms in 2022–2023; analyses done separately for each species and each subhabitat. Univariate analyses conducted using PERMANOVA, with Bray-Curtis coefficient (Anderson et al., 2008). Site type and Round are fixed factors. See Table S1 for dates of sampling rounds and Table S4.1 for the number of fish retained from each round. P-values bolded for significant and interpretable fixed effects; 998–999 unique permutations. Analysis of components of variation conducted for both fixed and random effects to evaluate relative importance of spatiotemporal scales.

| Response variable | Project | Species | Subhabitat | PERMANOVA | | | | | Components of variation | |
| --- | --- | --- | --- | --- | --- | --- | --- | --- | --- | --- |
|  |  |  |  | Sources of variation | df | MS | Pseudo-F | P(perm) | Estimate | % |
| Total fish length | Aulac | Tomcod | Creek | Site type | 1 | 131 | 1.15 | 0.302 | 0.4 | 0.27 |
|  |  |  |  | Round | 5 | 313 | 2.75 | 0.024 | 14.8 | 9.10 |
|  |  |  |  | Site type*Round | 5 | 344 | 3.03 | **0.010** | 34.3 | 21.04 |
|  |  |  |  | Error (i.e., Fish) | 71 | 113 |  |  | 113.5 | 69.60 |
|  |  | Mummichog | Creek | Site type | 1 | 27 | 0.45 | 0.511 | 0.0 | 0.00 |
|  |  |  |  | Round | 3 | 242 | 4.04 | **0.014** | 16.6 | 21.73 |
|  |  |  |  | Site type*Round | 3 | 20 | 0.34 | 0.812 | 0.0 | 0.00 |
|  |  |  |  | Error (i.e., Fish) | 37 | 60 |  |  | 60.0 | 78.27 |
|  |  |  | Platform | Site type | 1 | 235 | 2.64 | 0.121 | 10.2 | 8.96 |
|  |  |  |  | Round | 2 | 12 | 0.13 | 0.898 | 0.0 | 0.00 |
|  |  |  |  | Site type*Round | 2 | 159 | 1.78 | 0.181 | 14.0 | 12.31 |
|  |  |  |  | Error (i.e., Fish) | 25 | 89 |  |  | 89.2 | 78.73 |
|  | Wallace Bay | Mummichog | Creek | Site type | 1 | 82 | 1.00 | 0.321 | 0.0 | 0.02 |
|  |  |  |  | Round | 3 | 32 | 0.39 | 0.764 | 0.0 | 0.00 |
|  |  |  |  | Site type*Round | 3 | 98 | 1.20 | 0.344 | 3.2 | 3.79 |
|  |  |  |  | Error (i.e., Fish) | 34 | 82 |  |  | 81.9 | 96.19 |
|  |  |  | Platform | Site type | 1 | 6 | 0.09 | 0.812 | 0.0 | 0.00 |
|  |  |  |  | Round | 2 | 90 | 1.32 | 0.278 | 2.1 | 2.69 |
|  |  |  |  | Site type*Round | 2 | 100 | 1.47 | 0.260 | 6.1 | 7.97 |
|  |  |  |  | Error (i.e., Fish) | 27 | 68 |  |  | 68.0 | 89.34 |
| % vegetal matter | Aulac | Tomcod | Creek | Site type | 1 | 11862 | 6.00 | 0.011 | 247.2 | 7.82 |
|  |  |  |  | Round | 5 | 8929 | 4.51 | 0.001 | 528.0 | 16.70 |
|  |  |  |  | Site type*Round | 5 | 4721 | 2.39 | **0.024** | 408.8 | 12.93 |
|  |  |  |  | Error (i.e., Fish) | 71 | 1979 |  |  | 1978.5 | 62.56 |
|  |  | Mummichog | Creek | Site | 1 | 3683 | 1.80 | 0.163 | 75.3 | 3.25 |
|  |  |  |  | Round | 3 | 4043 | 1.97 | 0.102 | 182.2 | 7.86 |
|  |  |  |  | Site type*Round | 3 | 2113 | 1.03 | 0.397 | 11.5 | 0.49 |
|  |  |  |  | Error (i.e., Fish) | 37 | 2050 |  |  | 2050.1 | 88.40 |
|  |  |  | Platform | Site type | 1 | 865 | 0.59 | 0.483 | 0.0 | 0.00 |
|  |  |  |  | Round | 2 | 5031 | 3.46 | **0.030** | 357.5 | 19.18 |
|  |  |  |  | Site type*Round | 2 | 1709 | 1.17 | 0.321 | 50.6 | 2.71 |
|  |  |  |  | Error (i.e., Fish) | 25 | 1456 |  |  | 1455.8 | 78.11 |
|  | Wallace Bay | Mummichog | Creek | Site type | 1 | 30635 | 13.64 | **0.001** | 1434.2 | 37.83 |
|  |  |  |  | Round | 3 | 3363 | 1.50 | 0.206 | 111.1 | 2.93 |
|  |  |  |  | Site type*Round | 3 | 1066 | 0.47 | 0.835 | 0.0 | 0.00 |
|  |  |  |  | Error (i.e., Fish) | 34 | 2246 |  |  | 2246.0 | 59.24 |
|  |  |  | Platform | Site type | 1 | 3622 | 1.67 | 0.195 | 94.3 | 2.67 |
|  |  |  |  | Round | 2 | 2338 | 1.08 | 0.355 | 16.3 | 0.46 |
|  |  |  |  | Site type*Round | 2 | 8734 | 4.03 | **0.011** | 1251.0 | 35.46 |
|  |  |  |  | Error (i.e., Fish) | 27 | 2167 |  |  | 2166.8 | 61.41 |

**Table S4.5** PERMANOVA results for prey assemblage (4^th^-root transformed) for tomcods (*Microgadus tomcod*) and mummichogs (*Fundulus heteroclitus*) in Aulac and Wallace Bay reference and restoring salt marsh creeks and platforms in 2022–2023; analyses done separately for each species and each subhabitat. See Table S1 for dates of sampling rounds, Table S4.1 for the number of fish retained from each round, and Table S4.2 for prey assemblage. Site type and Round are fixed factors. This table complements Table 4 to assess if the patterns detected differ between an analysis on taxa densities 4^th^ root transformed and vegetal sources (detritus and macroalgae) omitted versus an analysis on taxa densities converted to presence-absence with vegetal sources included. P-values bolded for significant and interpretable fixed effects; 997–999 unique permutations. PERMDISP tests conducted for significant fixed effects to assess amount of multivariate dispersion among groups; df1 and df2 represent the numerator and denominator degrees of freedom for the F-ratio, respectively. Analysis of components of variation conducted for both fixed and random effects to evaluate relative importance of spatiotemporal scales.

|  |  |  | PERMANOVA | | | | | Components of variation | | PERMDISP | | |
| --- | --- | --- | --- | --- | --- | --- | --- | --- | --- | --- | --- | --- |
| Project | Species | Subhabitat | Sources of variation | df | MS | Pseudo-F | P (perm) | Estimate | % | F | df 1, df 2 | P |
| Aulac | Tomcod | Creek | Site type | 1 | 9066 | 8.73 | 0.001 | 200 | 11.1 | 0.24 | 1, 81 | 0.654 |
|  |  |  | Round | 5 | 7331 | 7.06 | 0.001 | 469 | 25.9 | 4.55 | 5, 77 | 0.003 |
|  |  |  | Site type*Round | 5 | 1722 | 1.66 | **0.027** | 102 | 5.6 | 6.09 | 11, 71 | 0.001 |
|  |  |  | Error (i.e., Fish) | 71 | 1038 |  |  | 1038 | 57.4 |  |  |  |
|  | Mummichog | Creek | Site type | 1 | 1118 | 3.57 | **0.001** | 371 | 9.2 | 16.83 | 1, 43 | 0.002 |
|  |  |  | Round | 3 | 5553 | 1.77 | **0.020** | 221 | 5.5 | 3.72 | 3, 41 | 0.052 |
|  |  |  | Site type*Round | 3 | 4763 | 1.52 | 0.067 | 298 | 7.4 |  |  |  |
|  |  |  | Error (i.e., Fish) | 37 | 3134 |  |  | 3134 | 77.9 |  |  |  |
|  |  | Platform | Site type | 1 | 10712 | 3.29 | **0.004** | 518 | 12.2 | 3.54 | 1, 29 | 0.090 |
|  |  |  | Round | 2 | 7902 | 2.43 | **0.005** | 465 | 11.0 | 1.88 | 2, 28 | 0.228 |
|  |  |  | Site type*Round | 2 | 3008 | 0.92 | 0.536 | 0 | 0.0 |  |  |  |
|  |  |  | Error (i.e., Fish ) | 25 | 3257 |  |  | 3257 | 76.8 |  |  |  |
| Wallace Bay | Mummichog | Creek | Site type | 1 | 19310 | 9.03 | 0.001 | 867.5 | 20.8 | 1.24 | 1, 40 | 0.335 |
|  |  |  | Round | 3 | 7476 | 3.49 | 0.001 | 530.8 | 12.7 | 0.12 | 3, 38 | 0.962 |
|  |  |  | Site type*Round | 3 | 5338 | 2.50 | **0.005** | 636.3 | 15.2 | 4.40 | 7, 34 | 0.019 |
|  |  |  | Error (i.e., Fish ) | 34 | 2139 |  |  | 2139.2 | 51.3 |  |  |  |
|  |  | Platform | Site type | 1 | 5539 | 2.56 | **0.027** | 218.7 | 7.5 | 0.83 | 1, 31 | 0.419 |
|  |  |  | Round | 2 | 6262 | 2.89 | **0.004** | 390.2 | 13.4 | 2.71 | 2, 30 | 0.112 |
|  |  |  | Site type*Round | 2 | 2849 | 1.32 | 0.217 | 130.2 | 4.5 |  |  |  |
|  |  |  | Error (i.e., Fish) | 27 | 2165 |  |  | 2165.0 | 74.6 |  |  |  |

**Table S4.6** Pairwise comparisons examining a significant Site type*Round interaction and focussing on the site type effect (reference vs. restoring) for gut fullness, prey assemblage, and total fish length of tomcods (*Microgadus tomcod*) and mummichogs (*Fundulus heteroclitus*) captured in two subhabitats in salt marshes in Aulac and Wallace Bay in 2022‒2023. P-values bolded for significant comparisons. When the number of unique permutations is low (< 100), P-values obtained by Monte Carlo simulations (P(MC)) should be used. See Table 4 for main PERMANOVAs, and Table S4.4 for supplementary permutational ANOVAs. See also Table S4.1 for number of fish included during each round (and reflecting the degrees of freedom).

| Project | Species | Subhabitat | Response variable | Source | Round | df | t | P(perm) | Unique permutations | P(MC) |
| --- | --- | --- | --- | --- | --- | --- | --- | --- | --- | --- |
| Aulac | Tomcod | Creek | Prey assemblage | Site type*Round | May 2022 | 10 | 1.08 | 0.356 | 267 | 0.330 |
|  |  |  |  |  | June 2022 | 14 | 2.76 | **0.001** | 439 | 0.002 |
|  |  |  |  |  | July 2022 | 14 | 1.61 | 0.069 | 246 | 0.087 |
|  |  |  |  |  | May 2023 | 12 | 0.99 | 0.446 | 852 | 0.418 |
|  |  |  |  |  | June 2023 | 10 | 1.21 | 0.245 | 308 | 0.251 |
|  |  |  |  |  | July 2023 | 11 | 2.09 | **0.005** | 767 | 0.013 |
|  | Mummichog | Creek | Prey assemblage | Site type*Round | May 2022 | 8 | 1.03 | 0.378 | 206 | 0.396 |
|  |  |  |  |  | May 2023 | 11 | 1.53 | **0.033** | 687 | 0.055 |
|  |  |  |  |  | June 2023 | 10 | 2.06 | **0.005** | 236 | 0.010 |
|  |  |  |  |  | July 2023 | 8 | 1.05 | 0.483 | 91 | 0.367 |
| Wallace Bay | Mummichog | Creek | Gut fullness | Site type*Round | May 2022 | 8 | 2.60 | **0.003** | 153 | 0.008 |
|  |  |  |  |  | May 2023 | 10 | 1.33 | 0.149 | 410 | 0.162 |
|  |  |  |  |  | June 2023 | 9 | 2.08 | **0.003** | 202 | 0.013 |
|  |  |  |  |  | July 2023 | 7 | 2.23 | 0.030 | 31 | **0.025** |
|  |  | Creek | Prey assemblage | Site type*Round | May 2022 | 8 | 1.72 | **0.017** | 113 | 0.062 |
|  |  |  |  |  | May 2023 | 10 | 1.33 | 0.150 | 404 | 0.166 |
|  |  |  |  |  | June 2023 | 9 | 2.08 | **0.011** | 205 | 0.014 |
|  |  |  |  |  | July 2023 | 7 | 2.23 | 0.032 | 31 | **0.019** |
|  |  | Platform | Prey assemblage | Site type*Round | June 2023 | 10 | 1.00 | 0.461 | 407 | 0.408 |
|  |  |  |  |  | July 2022 | 10 | 1.81 | **0.002** | 410 | 0.022 |
|  |  |  |  |  | July 2023 | 7 | 1.54 | 0.034 | 63 | 0.093 |
| Aulac | Tomcod | Creek | Total fish length | Site type*Round | May 2022 | 10 | 1.73 | 0.124 | 312 | 0.115 |
|  |  |  |  |  | June 2022 | 14 | 0.96 | 0.391 | 915 | 0.343 |
|  |  |  |  |  | July 2022 | 14 | 1.54 | 0.173 | 931 | 0.149 |
|  |  |  |  |  | May 2023 | 12 | 1.42 | 0.164 | 823 | 0.168 |
|  |  |  |  |  | June 2023 | 10 | 0.30 | 0.821 | 403 | 0.787 |
|  |  |  |  |  | July 2023 | 11 | 5.07 | **0.002** | 668 | 0.001 |

**Table S4.7 SEE ATTACHED EXCEL FILE – TABLE TOO LARGE TO BE INCLUDED IN WORD DOCUMENT.** Mean ± SE of gut items found in tomcods (*Microgadus tomcod*) and mummichogs (*Fundulus heteroclitus*) in creeks and on platforms in reference and restoring salt marshes in Aulac and Wallace Bay. Fish were captured in fyke nets (creek) and seine hauls (platform) and retained from 3 rounds (May, June, and July) in 2022–2023. Data averaged for each Species-Subhabitat-Site type-Round combination. Vegetal items (macroalgae and detritus) are omitted since they were recorded as either present or absent.

**Supplement 5. Diversity analyses for the faunal communities in intertidal creeks, on marsh platforms, and in salt pools in Aulac and Wallace Bay, Maritime Canada**

**Table S5.1** Mean ± SE species richness, diversity (Inverse Simpson Index), and evenness (Simpson Evenness) of faunal communities per net or trap in intertidal creeks, marsh platforms, and salt pools in reference and restoring salt marshes in Aulac and Wallace Bay. Sampling occurred over 3 rounds (May, June, and July) in 2022–2023. Data pooled over fyke nets, seine hauls, minnow traps, and invertebrate activity traps per site type and round. Aulac invertebrate activity traps deployed in restoring pools were mostly empty over the rounds (Table 1); therefore, this capture method is omitted from the Aulac analysis. Wallace Bay invertebrate activity traps in restoring pools for June 2022 and July 2023 were empty, and so omitted from analysis. Species richness is the number of taxa in a sample. Inverse Simpson Index was calculated as the reciprocal of the summed squared proportions of species densities in a sample, and ranges from a minimum of 1 to a maximum equal to the species richness of the sample when species are equally frequent. Simpson Evenness is calculated as the Inverse Simpson Index divided by the maximum species richness for a community, and ranges between very small values when a species dominates to a maximum of 1 when the species of equal in abundance (Lande 1996, Bittinger 2020). Maximum species richness used for fyke nets were 14 and 23, for seine nets 14 and 10, and for minnow traps 9 and 12 for Aulac and Wallace Bay, respectively. For Wallace Bay invertebrate activity traps, the maximum species richness used was 9.

| Project | Subhabitat | Capture method | Year | Month | Site type | Species richness | Species diversity | Species evenness |
| --- | --- | --- | --- | --- | --- | --- | --- | --- |
| Aulac | Creek | Fyke net | 2022 | May | Reference | 6.50±1.50 | 2.37±0.07 | 0.17±0.00 |
|  |  |  |  |  | Restoring | 5.00±0.00 | 2.77±0.02 | 0.20±0.00 |
|  |  |  |  | June | Reference | 7.00±0.00 | 3.61±0.43 | 0.26±0.03 |
|  |  |  |  |  | Restoring | 6.00±0.00 | 1.83±0.39 | 0.13±0.03 |
|  |  |  |  | July | Reference | 6.50±0.50 | 2.57±0.34 | 0.17±0.05 |
|  |  |  |  |  | Restoring | 6.50±0.50 | 2.57±0.34 | 0.18±0.02 |
|  |  |  | 2023 | May | Reference | 6.00±2.00 | 2.24±0.36 | 0.16±0.03 |
|  |  |  |  |  | Restoring | 6.00±0.00 | 3.16±0.01 | 0.23±0.00 |
|  |  |  |  | June | Reference | 5.50±3.50 | 2.64±1.42 | 0.19±0.10 |
|  |  |  |  |  | Restoring | 6.00±1.00 | 2.05±0.77 | 0.15±0.06 |
|  |  |  |  | July | Reference | 5.50±0.50 | 2.07±0.36 | 0.15±0.03 |
|  |  |  |  |  | Restoring | 7.50±0.50 | 2.72±0.27 | 0.19±0.02 |
|  | Marsh platform | Seine net | 2022 | May | Reference | 3.00±1.00 | 2.24±0.34 | 0.16±0.02 |
|  |  |  |  |  | Restoring | 5.00±0.58 | 2.35±0.49 | 0.17±0.04 |
|  |  |  |  | June | Reference | 6.00±0.00 | 3.06±0.23 | 0.22±0.02 |
|  |  |  |  |  | Restoring | 6.00±0.58 | 4.66±0.29 | 0.33±0.02 |
|  |  |  |  | July | Reference | 5.33±0.33 | 3.31±0.39 | 0.24±0.03 |
|  |  |  |  |  | Restoring | 4.33±1.45 | 3.99±1.29 | 0.29±0.09 |
|  |  |  | 2023 | May | Reference | 5.00±1.00 | 3.12±0.27 | 0.22±0.02 |
|  |  |  |  |  | Restoring | 4.33±0.88 | 2.92±0.40 | 0.21±0.03 |
|  |  |  |  | June | Reference | 6.00±0.58 | 3.28±0.43 | 0.23±0.03 |
|  |  |  |  |  | Restoring | 5.33±0.67 | 2.86±0.45 | 0.20±0.03 |
|  |  |  |  | July | Reference | 6.33±0.88 | 2.07±0.30 | 0.15±0.02 |
|  |  |  |  |  | Restoring | 2.33±0.88 | 1.70±0.37 | 0.12±0.03 |
|  | Salt pool | Minnow trap | 2022 | May | Reference | 0.75±0.48 | 0.26±0.15 | 0.03±0.02 |
|  |  |  |  |  | Restoring | 1.75±0.25 | 0.59±0.03 | 0.07±0.00 |
|  |  |  |  | June | Reference | 0.50±0.29 | 0.25±0.14 | 0.03±0.02 |
|  |  |  |  |  | Restoring | 1.00±0.41 | 0.40±0.13 | 0.04±0.01 |
|  |  |  |  | July | Reference | 0.75±0.48 | 0.29±0.17 | 0.03±0.02 |
|  |  |  |  |  | Restoring | 0.75±0.25 | 0.38±0.13 | 0.04±0.01 |
|  |  |  | 2023 | May | Reference | 0.75±0.48 | 0.29±0.17 | 0.03±0.02 |
|  |  |  |  |  | Restoring | 1.25±0.25 | 0.50±0.00 | 0.06±0.00 |
|  |  |  |  | June | Reference | 0.50±0.29 | 0.25±0.14 | 0.03±0.02 |
|  |  |  |  |  | Restoring | 1.75±0.48 | 0.52±0.01 | 0.06±0.00 |
|  |  |  |  | July | Reference | 0.67±0.67 | 0.18±0.18 | 0.02±0.02 |
|  |  |  |  |  | Restoring | 1.00±0.00 | 0.50±0.00 | 0.06±0.00 |
| Wallace Bay | Creek | Fyke net | 2022 | May | Reference | 13.50±0.50 | 1.93±0.21 | 0.08±0.01 |
|  |  |  |  |  | Restoring | 8.50±0.50 | 1.84±0.65 | 0.08±0.03 |
|  |  |  |  | June | Reference | 6.00±1.00 | 1.28±0.20 | 0.06±0.01 |
|  |  |  |  |  | Restoring | 8.50±1.50 | 3.06±1.87 | 0.13±0.08 |
|  |  |  |  | July | Reference | 5.00±1.00 | 1.50±0.48 | 0.07±0.02 |
|  |  |  |  |  | Restoring | 8.50±1.50 | 3.13±1.27 | 0.14±0.06 |
|  |  |  | 2023 | May | Reference | 8.00±0.00 | 1.97±0.46 | 0.09±0.02 |
|  |  |  |  |  | Restoring | 12.00±1.00 | 1.61±0.33 | 0.07±0.01 |
|  |  |  |  | June | Reference | 6.50±0.50 | 2.25±0.72 | 0.10±0.03 |
|  |  |  |  |  | Restoring | 8.50±0.50 | 4.06±0.86 | 0.18±0.04 |
|  |  |  |  | July | Reference | 3.00±0.00 | 1.72±0.58 | 0.07±0.03 |
|  |  |  |  |  | Restoring | 9.50±1.50 | 1.88±0.11 | 0.08±0.00 |
|  | Marsh platform | Seine net | 2022 | May | Reference | 2.33±0.88 | 1.95±0.53 | 0.19±0.05 |
|  |  |  |  |  | Restoring | 2.00±0.00 | 1.29±0.25 | 0.13±0.03 |
|  |  |  |  | June | Reference | 1.67±0.33 | 1.15±0.08 | 0.11±0.01 |
|  |  |  |  |  | Restoring | 3.33±0.33 | 2.54±0.30 | 0.25±0.03 |
|  |  |  |  | July | Reference | 1.33±0.33 | 1.09±0.09 | 0.11±0.01 |
|  |  |  |  |  | Restoring | 3.33±0.88 | 1.41±0.20 | 0.14±0.02 |
|  |  |  | 2023 | May | Reference | 1.33±0.33 | 1.03±0.03 | 0.10±0.00 |
|  |  |  |  |  | Restoring | 2.33±0.33 | 2.16±0.45 | 0.22±0.04 |
|  |  |  |  | June | Reference | 1.67±0.33 | 1.76±0.49 | 0.18±0.05 |
|  |  |  |  |  | Restoring | 1.33±0.33 | 1.23±0.23 | 0.12±0.02 |
|  |  |  |  | July | Reference | 1.00±0.00 | 1.00±0.00 | 0.10±0.00 |
|  |  |  |  |  | Restoring | 2.00±0.00 | 1.06±0.02 | 0.11±0.00 |
|  | Salt pool | Minnow trap | 2022 | May | Reference | 1.50±0.50 | 1.03±0.03 | 0.09±0.00 |
|  |  |  |  |  | Restoring | 2.75±0.75 | 2.47±0.70 | 0.21±0.06 |
|  |  |  |  | June | Reference | 1.50±0.29 | 1.22±0.19 | 0.10±0.22 |
|  |  |  |  |  | Restoring | 3.25±0.48 | 2.04±0.30 | 0.17±0.02 |
|  |  |  |  | July | Reference | 1.75±0.25 | 1.55±0.21 | 0.13±0.02 |
|  |  |  |  |  | Restoring | 2.50±0.29 | 2.14±0.30 | 0.18±0.02 |
|  |  |  | 2023 | May | Reference | 2.00±0.00 | 1.86±0.03 | 0.16±0.00 |
|  |  |  |  |  | Restoring | 3.50±0.29 | 2.34±0.59 | 0.20±0.05 |
|  |  |  |  | June | Reference | 2.00±0.58 | 1.47±0.26 | 0.12±0.02 |
|  |  |  |  |  | Restoring | 2.00±0.41 | 1.77±0.28 | 0.15±0.02 |
|  |  |  |  | July | Reference | 1.00±0.00 | 1.00±0.00 | 0.08±0.00 |
|  |  |  |  |  | Restoring | 1.75±0.25 | 1.07±0.03 | 0.09±0.00 |
|  |  | Invertebrate activity trap | 2022 | May | Reference | 1.00±0.00 | 1.00±0.00 | 0.11±0.00 |
|  |  |  |  |  | Restoring | 1.75±0.25 | 1.75±0.25 | 0.19±0.03 |
|  |  |  |  | June | Reference | 1.00±0.00 | 1.00±0.00 | 0.11±0.00 |
|  |  |  |  | July | Reference | 1.75±0.48 | 1.48±0.28 | 0.16±0.03 |
|  |  |  |  |  | Restoring | 1.00±0.00 | 1.00±0.00 | 0.11±0.00 |
|  |  |  | 2023 | May | Reference | 1.00±0.00 | 1.00±0.00 | 0.11±0.00 |
|  |  |  |  |  | Restoring | 1.00±0.00 | 1.00±0.00 | 0.11±0.00 |
|  |  |  |  | June | Reference | 1.25±0.25 | 0.88±0.12 | 0.10±0.01 |
|  |  |  |  |  | Restoring | 1.25±0.25 | 1.15±0.15 | 0.13±0.02 |
|  |  |  |  | July | Reference | 2.00±0.00 | 1.66±0.16 | 0.18±0.02 |

**Table S5.2** Permutational ANOVA results for differences in species richness, diversity (Inverse Simpson Index), evenness (Simpson Evenness), and total number of individuals for nekton communities in intertidal creeks, on marsh platforms, and in salt pools in Aulac reference and restoring salt marshes in 2022–2023. Univariate analyses conducted using PERMANOVA, with Bray-Curtis coefficient (Anderson et al., 2008). Site type, Year and Month are fixed factors, and Pool is a random factor. See Table S1 for dates of sampling rounds. P-values bolded for significant and interpretable fixed effects; 170–999 unique permutations. Analysis of components of variation conducted for both fixed and random effects to evaluate relative importance of spatiotemporal scales. Note that for minnow traps in salt pools, the error term is actually estimated by Year*Month*Pool(Site type).

|  |  | PERMANOVA | |  |  |  | Components of variation | |
| --- | --- | --- | --- | --- | --- | --- | --- | --- |
| Subhabitat | Community | Sources of variation | df | MS | Pseudo-F | P(perm) | Estimate | % |
| Creek | Species richness | Site type | 1 | 83 | 0.31 | 0.658 | 0.0 | 0.0 |
|  |  | Year | 1 | 104 | 0.38 | 0.577 | 0.0 | 0.0 |
|  |  | Month | 2 | 119 | 0.44 | 0.699 | 0.0 | 0.0 |
|  |  | Site type*Year | 1 | 425 | 1.57 | 0.226 | 25.7 | 8.7 |
|  |  | Site type*Month | 2 | 122 | 0.45 | 0.683 | 0.0 | 0.0 |
|  |  | Year*Month | 2 | 105 | 0.39 | 0.757 | 0.0 | 0.0 |
|  |  | Site type*Year*Month | 2 | 52 | 0.19 | 0.890 | 0.0 | 0.0 |
|  |  | Error (i.e., Net) | 12 | 271 |  |  | 270.6 | 91.3 |
|  | Species diversity | Site type | 1 | 15 | 0.05 | 0.863 | 0.0 | 0.0 |
|  |  | Year | 1 | 60 | 0.22 | 0.662 | 0.0 | 0.0 |
|  |  | Month | 2 | 159 | 0.58 | 0.596 | 0.0 | 0.0 |
|  |  | Site type*Year | 1 | 302 | 1.09 | 0.296 | 4.2 | 1.1 |
|  |  | Site type*Month | 2 | 655 | 2.37 | 0.135 | 94.7 | 25.2 |
|  |  | Year*Month | 2 | 70 | 0.25 | 0.805 | 0.0 | 0.0 |
|  |  | Site type*Year*Month | 2 | 50 | 0.18 | 0.850 | 0.0 | 0.0 |
|  |  | Error (i.e., Net) | 12 | 276 |  |  | 276.3 | 73.6 |
|  | Species evenness | Site type | 1 | 15 | 0.05 | 0.868 | 0.0 | 0.0 |
|  |  | Year | 1 | 60 | 0.22 | 0.664 | 0.0 | 0.0 |
|  |  | Month | 2 | 159 | 0.58 | 0.574 | 0.0 | 0.0 |
|  |  | Site type*Year | 1 | 302 | 1.09 | 0.317 | 4.2 | 1.1 |
|  |  | Site type*Month | 2 | 655 | 2.37 | 0.142 | 94.7 | 25.2 |
|  |  | Year*Month | 2 | 70 | 0.25 | 0.784 | 0.0 | 0.0 |
|  |  | Site type*Year*Month | 2 | 50 | 0.18 | 0.883 | 0.0 | 0.0 |
|  |  | Error (i.e., Net) | 12 | 276 |  |  | 276.3 | 73.6 |
|  | Total number of individuals | Site type | 1 | 1631 | 0.90 | 0.411 | 0.0 | 0.0 |
|  |  | Year | 1 | 965 | 0.54 | 0.624 | 0.0 | 0.0 |
|  |  | Month | 2 | 2127 | 1.18 | 0.325 | 40.4 | 1.9 |
|  |  | Site type*Year | 1 | 3539 | 1.96 | 0.171 | 289.2 | 13.6 |
|  |  | Site type*Month | 2 | 287 | 0.16 | 0.981 | 0.0 | 0.0 |
|  |  | Year*Month | 2 | 1180 | 0.65 | 0.629 | 0.0 | 0.0 |
|  |  | Site type*Year*Month | 2 | 781 | 0.43 | 0.802 | 0.0 | 0.0 |
|  |  | Error (i.e., Net) | 12 | 1804 |  |  | 1803.6 | 84.5 |
| Marsh platform | Species richness | Site type | 1 | 534 | 1.84 | 0.151 | 13.6 | 1.9 |
|  |  | Year | 1 | 473 | 1.63 | 0.193 | 10.2 | 1.4 |
|  |  | Month | 2 | 1069 | 3.69 | 0.024 | 64.9 | 9.1 |
|  |  | Site type*Year | 1 | 19 | 0.06 | 0.959 | 0.0 | 0.0 |
|  |  | Site type*Month | 2 | 1326 | 4.58 | **0.009** | 172.7 | 24.3 |
|  |  | Year*Month | 2 | 80 | 0.28 | 0.872 | 0.0 | 0.0 |
|  |  | Site type*Year*Month | 2 | 766 | 2.64 | 0.059 | 158.7 | 22.4 |
|  |  | Error (i.e., Haul) | 24 | 290 |  |  | 289.9 | 40.8 |
|  | Species diversity | Site type | 1 | 148 | 0.70 | 0.461 | 0.0 | 0.0 |
|  |  | Year | 1 | 2664 | 12.56 | **0.001** | 136.2 | 33.4 |
|  |  | Month | 2 | 731 | 3.45 | **0.038** | 43.3 | 10.6 |
|  |  | Site type*Year | 1 | 249 | 1.17 | 0.290 | 4.1 | 1.0 |
|  |  | Site type*Month | 2 | 91 | 0.43 | 0.743 | 0.0 | 0.0 |
|  |  | Year*Month | 2 | 284 | 1.34 | 0.283 | 11.9 | 2.9 |
|  |  | Site type*Year*Month | 2 | 211 | 1.00 | 0.393 | 0.0 | 0.0 |
|  |  | Error (i.e., Haul) | 24 | 212 |  |  | 212.2 | 52.0 |
|  | Species evenness | Site type | 1 | 148 | 0.70 | 0.481 | 0.0 | 0.0 |
|  |  | Year | 1 | 2664 | 12.56 | **0.002** | 136.2 | 33.4 |
|  |  | Month | 2 | 731 | 3.45 | **0.030** | 43.3 | 10.6 |
|  |  | Site type*Year | 1 | 249 | 1.17 | 0.297 | 4.1 | 1.0 |
|  |  | Site type*Month | 2 | 91 | 0.43 | 0.739 | 0.0 | 0.0 |
|  |  | Year*Month | 2 | 284 | 1.34 | 0.253 | 11.9 | 2.9 |
|  |  | Site type*Year*Month | 2 | 211 | 1.00 | 0.383 | 0.0 | 0.0 |
|  |  | Error (i.e., Haul) | 24 | 212 |  |  | 212.2 | 52.0 |
|  | Total number of individuals | Site type | 1 | 14761 | 20.23 | 0.001 | 779.5 | 27.2 |
|  |  | Year | 1 | 1793 | 2.46 | 0.068 | 59.1 | 2.1 |
|  |  | Month | 2 | 4226 | 5.79 | 0.001 | 291.4 | 10.2 |
|  |  | Site type*Year | 1 | 518 | 0.71 | 0.534 | 0.0 | 0.0 |
|  |  | Site type*Month | 2 | 2531 | 3.47 | 0.003 | 300.2 | 10.5 |
|  |  | Year*Month | 2 | 1373 | 1.88 | 0.095 | 107.2 | 3.7 |
|  |  | Site type*Year*Month | 2 | 2529 | 3.47 | **0.012** | 599.7 | 20.9 |
|  |  | Error (i.e., Haul) | 24 | 730 |  |  | 729.7 | 25.5 |
| Salt pool | Species richness | Site type | 1 | 218 | 0.59 | 0.501 | 0.0 | 0.0 |
| (Minnow | | Year | 1 | 102 | 0.40 | 0.590 | 0.0 | 0.0 |
| traps) |  | Month | 2 | 469 | 1.64 | 0.233 | 31.9 | 4.9 |
|  |  | Site type*Year | 1 | 95 | 0.37 | 0.582 | 0.0 | 0.0 |
|  |  | Site type*Month | 2 | 736 | 2.57 | 0.169 | 157.5 | 24.3 |
|  |  | Year*Month | 2 | 426 | 0.93 | 0.437 | 0.0 | 0.0 |
|  |  | Site type*Year*Month | 2 | 14 | 0.03 | 0.966 | 0.0 | 0.0 |
|  |  | Pool(Site type) | 5 | 218 | 0.48 | 0.796 | 0.0 | 0.0 |
|  |  | Year*Pool(Site type) | 4 | 116 | 0.25 | 0.879 | 0.0 | 0.0 |
|  |  | Month*Pool(Site type) | 8 | 218 | 0.48 | 0.829 | 0.0 | 0.0 |
|  |  | Error (i.e., Sample) | 4 | 458 |  |  | 457.9 | 70.7 |
|  | Species diversity | Site type | 1 | 12.4 | 0.81 | 0.426 | 0.0 | 0.0 |
|  |  | Year | 1 | 46.8 | 5.08 | 0.100 | 6.0 | 20.0 |
|  |  | Month | 2 | 34.0 | 1.87 | 0.209 | 2.8 | 9.2 |
|  |  | Site type*Year | 1 | 0.67 | 0.07 | 0.808 | 0.0 | 0.0 |
|  |  | Site type*Month | 2 | 22.3 | 1.23 | 0.349 | 1.5 | 4.8 |
|  |  | Year*Month | 2 | 9.5 | 0.48 | 0.658 | 0.0 | 0.0 |
|  |  | Site type*Year*Month | 2 | 19.9 | 1.00 | 0.453 | 0.0 | 0.2 |
|  |  | Pool(Site type) | 5 | 7.8 | 0.39 | 0.853 | 0.0 | 0.0 |
|  |  | Year*Pool(Site type) | 4 | 1.7 | 0.09 | 0.981 | 0.0 | 0.0 |
|  |  | Month*Pool(Site type) | 8 | 17.5 | 0.89 | 0.609 | 0.0 | 0.0 |
|  |  | Error (i.e., Sample) | 4 | 19.8 |  |  | 19.8 | 65.8 |
|  | Species evenness | Site type | 1 | 12.4 | 0.81 | 0.464 | 0.0 | 0.0 |
|  |  | Year | 1 | 46.8 | 5.08 | 0.105 | 6.0 | 20.0 |
|  |  | Month | 2 | 34.0 | 1.87 | 0.213 | 2.8 | 9.2 |
|  |  | Site type*Year | 1 | 0.67 | 0.07 | 0.817 | 0.0 | 0.0 |
|  |  | Site type*Month | 2 | 22.3 | 1.23 | 0.344 | 1.5 | 4.8 |
|  |  | Year*Month | 2 | 9.5 | 0.48 | 0.683 | 0.0 | 0.0 |
|  |  | Site type*Year*Month | 2 | 19.9 | 1.00 | 0.445 | 0.0 | 0.2 |
|  |  | Pool(Site type) | 5 | 7.8 | 0.39 | 0.844 | 0.0 | 0.0 |
|  |  | Year*Pool(Site type) | 4 | 1.7 | 0.09 | 0.969 | 0.0 | 0.0 |
|  |  | Month*Pool(Site type) | 8 | 17.5 | 0.89 | 0.581 | 0.0 | 0.0 |
|  |  | Error (i.e., Sample) | 4 | 19.8 |  |  | 19.8 | 65.8 |
|  | Total number of individuals | Site type | 1 | 21790 | 2.67 | 0.183 | 615.4 | 13.9 |
|  |  | Year | 1 | 986 | 0.59 | 0.609 | 0.0 | 0.0 |
|  |  | Month | 2 | 1854 | 1.01 | 0.406 | 0.7 | 0.0 |
|  |  | Site type*Year | 1 | 2500 | 1.50 | 0.249 | 75.1 | 1.7 |
|  |  | Site type*Month | 2 | 2101 | 1.14 | 0.348 | 34.6 | 0.8 |
|  |  | Year*Month | 2 | 795 | 0.29 | 0.933 | 0.0 | 0.0 |
|  |  | Site type*Year*Month | 2 | 2512 | 0.93 | 0.481 | 0.0 | 0.0 |
|  |  | Pool(Site type) | 6 | 8283 | 3.06 | 0.009 | 983.8 | 22.3 |
|  |  | Year*Pool(Site type) | 6 | 1643 | 0.61 | 0.847 | 0.0 | 0.0 |
|  |  | Month*Pool(Site type) | 12 | 1816 | 0.67 | 0.873 | 0.0 | 0.0 |
|  |  | Error (i.e., Sample) | 11 | 2708 |  |  | 2708.4 | 61.3 |

**Table S5.3** Permutational ANOVA results for differences in species richness, diversity (Inverse Simpson Index), evenness (Simpson Evenness), and total number of individuals for nekton communities in intertidal creeks, on marsh platforms, and in salt pools in Wallace Bay reference and restoring salt marshes in 2022–2023. Empty invertebrate activity traps were removed prior to analysis. Univariate analyses conducted using PERMANOVA, with Bray-Curtis coefficient (Anderson et al., 2008). Site type, Year and Month are fixed factors, and Pool is a random factor. See Table S1 for dates of sampling rounds. P-values bolded for significant and interpretable fixed effects; 980–999 unique permutations. Analysis of components of variation conducted for both fixed and random effects to evaluate relative importance of spatiotemporal scales. Note that for minnow traps in salt pools, the error term is estimated by Year*Month*Pool(Site type); for invertebrate activity traps in salt pools, the error term comes from pooling available traps (that were not empty) over two years.

|  |  |  | PERMANOVA | | | | | Components of variation | |
| --- | --- | --- | --- | --- | --- | --- | --- | --- | --- |
| Subhabitat | Capture method | Community | Sources of variation | df | MS | Pseudo-F | P (perm) | Estimate | % |
| Creek | Fyke net | Species richness | Site type | 1 | 1795 | 24.46 | 0.001 | 143.5 | 18.45 |
|  |  |  | Year | 1 | 56 | 0.77 | 0.440 | 0.0 | 0.00 |
|  |  |  | Month | 2 | 1312 | 17.87 | 0.002 | 154.8 | 19.91 |
|  |  |  | Site type*Year | 1 | 555 | 7.56 | 0.011 | 80.3 | 10.32 |
|  |  |  | Site type*Month | 2 | 769 | 10.48 | 0.002 | 173.9 | 22.36 |
|  |  |  | Year*Month | 2 | 110 | 1.50 | 0.228 | 9.2 | 1.18 |
|  |  |  | Site type*Year*Month | 2 | 358 | 4.88 | **0.011** | 142.5 | 18.33 |
|  |  |  | Error (i.e., Net) | 12 | 73 |  |  | 73.4 | 9.44 |
|  |  | Species diversity | Site type | 1 | 1018 | 2.09 | 0.158 | 44.3 | 7.60 |
|  |  |  | Year | 1 | 191 | 0.39 | 0.576 | 0.0 | 0.00 |
|  |  |  | Month | 2 | 326 | 0.67 | 0.533 | 0.0 | 0.00 |
|  |  |  | Site type*Year | 1 | 150 | 0.31 | 0.669 | 0.0 | 0.00 |
|  |  |  | Site type*Month | 2 | 653 | 1.34 | 0.289 | 41.5 | 7.13 |
|  |  |  | Year*Month | 2 | 526 | 1.08 | 0.378 | 9.8 | 1.68 |
|  |  |  | Site type*Year*Month | 2 | 85 | 0.18 | 0.894 | 0.0 | 0.00 |
|  |  |  | Error (i.e., Net) | 12 | 487 |  |  | 487.1 | 83.60 |
|  |  | Species evenness | Site type | 1 | 1018 | 2.09 | 0.157 | 44.3 | 7.60 |
|  |  |  | Year | 1 | 191 | 0.39 | 0.576 | 0.0 | 0.00 |
|  |  |  | Month | 2 | 326 | 0.67 | 0.531 | 0.0 | 0.00 |
|  |  |  | Site type*Year | 1 | 150 | 0.31 | 0.675 | 0.0 | 0.00 |
|  |  |  | Site type*Month | 2 | 653 | 1.34 | 0.269 | 41.5 | 7.13 |
|  |  |  | Year*Month | 2 | 526 | 1.08 | 0.344 | 9.8 | 1.68 |
|  |  |  | Site type*Year*Month | 2 | 85 | 0.18 | 0.895 | 0.0 | 0.00 |
|  |  |  | Error (i.e., Net) | 12 | 487 |  |  | 487.1 | 83.60 |
|  |  | Total number of individuals | Site type | 1 | 1363 | 1.15 | 0.346 | 15.0 | 0.62 |
|  |  |  | Year | 1 | 2338 | 1.98 | 0.123 | 96.2 | 3.97 |
|  |  |  | Month | 2 | 3889 | 3.29 | **0.024** | 338.1 | 13.94 |
|  |  |  | Site type*Year | 1 | 2816 | 2.38 | 0.089 | 272.1 | 11.22 |
|  |  |  | Site type*Month | 2 | 2899 | 2.45 | 0.072 | 428.8 | 17.68 |
|  |  |  | Year*Month | 2 | 1551 | 1.31 | 0.301 | 92.0 | 3.79 |
|  |  |  | Site type*Year*Month | 2 | 842 | 0.71 | 0.613 | 0.0 | 0.00 |
|  |  |  | Error (i.e., Net) | 12 | 1184 |  |  | 1183.5 | 48.79 |
| Marsh platform | Seine net | Species richness | Site type | 1 | 3559 | 12.33 | 0.002 | 181.7 | 21.86 |
|  |  |  | Year | 1 | 1783 | 6.18 | 0.019 | 83.0 | 9.99 |
|  |  |  | Month | 2 | 97 | 0.33 | 0.833 | 0.0 | 0.00 |
|  |  |  | Site type*Year | 1 | 184 | 0.64 | 0.453 | 0.0 | 0.00 |
|  |  |  | Site type*Month | 2 | 629 | 2.18 | 0.113 | 56.7 | 6.82 |
|  |  |  | Year*Month | 2 | 168 | 0.58 | 0.613 | 0.0 | 0.00 |
|  |  |  | Site type*Year*Month | 2 | 951 | 3.30 | **0.041** | 220.9 | 26.59 |
|  |  |  | Error (i.e., Haul) | 24 | 289 |  |  | 288.6 | 34.74 |
|  |  | Species diversity | Site type | 1 | 672 | 3.57 | 0.039 | 26.9 | 3.23 |
|  |  |  | Year | 1 | 331 | 1.76 | 0.187 | 7.9 | 0.95 |
|  |  |  | Month | 2 | 750 | 3.98 | 0.030 | 46.8 | 5.62 |
|  |  |  | Site type*Year | 1 | 31 | 0.16 | 0.759 | 0.0 | 0.00 |
|  |  |  | Site type*Month | 2 | 29274 | 0.16 | 0.918 | 0.0 | 0.00 |
|  |  |  | Year*Month | 2 | 63 | 0.33 | 0.768 | 0.0 | 0.00 |
|  |  |  | Site type*Year*Month | 2 | 1876 | 9.95 | **0.001** | 562.6 | 67.56 |
|  |  |  | Error (i.e., Haul) | 24 | 189 |  |  | 188.5 | 22.64 |
|  |  | Species evenness | Site type | 1 | 672 | 3.57 | 0.061 | 26.9 | 3.23 |
|  |  |  | Year | 1 | 331 | 1.76 | 0.207 | 7.9 | 0.95 |
|  |  |  | Month | 2 | 750 | 3.98 | 0.032 | 46.8 | 5.62 |
|  |  |  | Site type*Year | 1 | 31 | 0.16 | 0.744 | 0.0 | 0.00 |
|  |  |  | Site type*Month | 2 | 29274 | 0.16 | 0.908 | 0.0 | 0.00 |
|  |  |  | Year*Month | 2 | 63 | 0.33 | 0.773 | 0.0 | 0.00 |
|  |  |  | Site type*Year*Month | 2 | 1876 | 9.95 | **0.003** | 562.6 | 67.56 |
|  |  |  | Error (i.e., Haul) | 24 | 189 |  |  | 188.5 | 22.64 |
|  |  | Total number of individuals | Site type | 1 | 8562 | 6.39 | 0.001 | 401.2 | 11.57 |
|  |  |  | Year | 1 | 1766 | 1.32 | 0.246 | 23.7 | 0.68 |
|  |  |  | Month | 2 | 2147 | 1.60 | 0.158 | 67.2 | 1.94 |
|  |  |  | Site type*Year | 1 | 3318 | 2.47 | 0.074 | 219.7 | 6.34 |
|  |  |  | Site type*Month | 2 | 7427 | 5.54 | **0.001** | 1014.4 | 29.26 |
|  |  |  | Year*Month | 2 | 931 | 0.69 | 0.620 | 0.0 | 0.00 |
|  |  |  | Site type*Year*Month | 2 | 2542 | 1.90 | 0.086 | 400.4 | 11.55 |
|  |  |  | Error (i.e., Haul) | 24 | 1341 |  |  | 1340.6 | 38.67 |
| Salt pool | Minnow trap | Species richness | Site type | 1 | 5152 | 13.23 | 0.005 | 215.0 | 27.90 |
|  |  |  | Year | 1 | 127 | 0.40 | 0.563 | 0.0 | 0.00 |
|  |  |  | Month | 2 | 645 | 2.11 | 0.152 | 22.9 | 2.97 |
|  |  |  | Site type*Year | 1 | 196 | 0.61 | 0.480 | 0.0 | 0.00 |
|  |  |  | Site type*Month | 2 | 138 | 0.45 | 0.727 | 0.0 | 0.00 |
|  |  |  | Year*Month | 2 | 1587 | 5.10 | **0.024** | 171.7 | 22.28 |
|  |  |  | Site type*Year*Month | 2 | 431 | 1.38 | 0.276 | 32.1 | 4.17 |
|  |  |  | Pool(Site type) | 6 | 391 | 1.26 | 0.338 | 14.1 | 1.83 |
|  |  |  | Year*Pool(Site type) | 6 | 321 | 1.03 | 0.463 | 3.5 | 0.45 |
|  |  |  | Month*Pool(Site type) | 12 | 306 | 0.98 | 0.499 | 0.0 | 0.00 |
|  |  |  | Error (i.e., Sample) | 11 | 311 |  |  | 311.4 | 40.41 |
|  |  | Species diversity | Site type | 1 | 2571 | 10.61 | 0.022 | 105.1 | 15.65 |
|  |  |  | Year | 1 | 196 | 2.04 | 0.186 | 4.5 | 0.67 |
|  |  |  | Month | 2 | 457 | 1.15 | 0.337 | 4.1 | 0.61 |
|  |  |  | Site type*Year | 1 | 759 | 7.91 | **0.038** | 59.9 | 8.92 |
|  |  |  | Site type*Month | 2 | 165 | 0.42 | 0.716 | 0.0 | 0.00 |
|  |  |  | Year*Month | 2 | 1675 | 16.09 | **0.002** | 211.4 | 31.48 |
|  |  |  | Site type*Year*Month | 2 | 94 | 0.90 | 0.442 | 0.0 | 0.00 |
|  |  |  | Pool(Site type) | 6 | 246 | 2.36 | 0.073 | 25.0 | 3.72 |
|  |  |  | Year*Pool(Site type) | 6 | 96 | 0.92 | 0.494 | 0.0 | 0.00 |
|  |  |  | Month*Pool(Site type) | 12 | 406 | 3.90 | 0.007 | 157.6 | 23.46 |
|  |  |  | Error (i.e., Sample) | 11 | 104 |  |  | 104.1 | 15.49 |
|  |  | Species evenness | Site type | 1 | 2571 | 10.61 | 0.017 | 105.1 | 15.65 |
|  |  |  | Year | 1 | 196 | 2.04 | 0.204 | 4.5 | 0.67 |
|  |  |  | Month | 2 | 457 | 1.15 | 0.331 | 4.1 | 0.61 |
|  |  |  | Site type*Year | 1 | 759 | 7.91 | **0.033** | 59.9 | 8.92 |
|  |  |  | Site type*Month | 2 | 165 | 0.42 | 0.704 | 0.0 | 0.00 |
|  |  |  | Year*Month | 2 | 1675 | 16.09 | **0.002** | 211.4 | 31.48 |
|  |  |  | Site type*Year*Month | 2 | 94 | 0.90 | 0.433 | 0.0 | 0.00 |
|  |  |  | Pool(Site type) | 6 | 246 | 2.36 | 0.080 | 25.0 | 3.72 |
|  |  |  | Year*Pool(Site type) | 6 | 96 | 0.92 | 0.511 | 0.0 | 0.00 |
|  |  |  | Month*Pool(Site type) | 12 | 406 | 3.90 | 0.011 | 157.6 | 23.46 |
|  |  |  | Error (i.e., Sample) | 11 | 104 |  |  | 104.1 | 15.49 |
|  |  | Total number of individuals | Site type | 1 | 472 | 0.20 | 0.832 | 0.0 | 0.00 |
|  |  |  | Year | 1 | 2628 | 1.43 | 0.256 | 35.9 | 1.45 |
|  |  |  | Month | 2 | 5568 | 3.59 | **0.025** | 270.4 | 10.91 |
|  |  |  | Site type*Year | 1 | 2546 | 1.39 | 0.259 | 64.3 | 2.60 |
|  |  |  | Site type*Month | 2 | 3366 | 2.17 | 0.085 | 244.3 | 9.86 |
|  |  |  | Year*Month | 2 | 1849 | 1.12 | 0.340 | 27.6 | 1.11 |
|  |  |  | Site type*Year*Month | 2 | 769 | 0.47 | 0.795 | 0.0 | 0.00 |
|  |  |  | Pool(Site type) | 6 | 2341 | 1.42 | 0.218 | 123.0 | 4.96 |
|  |  |  | Year*Pool(Site type) | 6 | 1837 | 1.12 | 0.390 | 68.1 | 2.75 |
|  |  |  | Month*Pool(Site type) | 12 | 1548 | 0.94 | 0.572 | 0.0 | 0.00 |
|  |  |  | Error (i.e., Sample) | 11 | 1644 |  |  | 1644.2 | 66.36 |
|  | Invertebrate activity trap | Species richness | Site type | 1 | 19 | 0.14 | 0.747 | 0.0 | 0.00 |
|  |  |  | Month | 2 | 151 | 0.72 | 0.509 | 0.0 | 0.00 |
|  |  |  | Site type*Month | 2 | 1020 | 4.83 | **0.030** | 182.5 | 38.00 |
|  |  |  | Pool(Site type) | 6 | 121 | 0.40 | 0.888 | 0.0 | 0.00 |
|  |  |  | Month*Pool(Site type) | 10 | 206 | 0.69 | 0.729 | 0.0 | 0.00 |
|  |  |  | Error (i.e., Sample) | 10 | 298 |  |  | 297.8 | 62.00 |
|  |  | Species diversity | Site type | 1 | 16 | 0.09 | 0.749 | 0.0 | 0.00 |
|  |  |  | Month | 2 | 236 | 1.83 | 0.189 | 12.1 | 3.64 |
|  |  |  | Site type*Month | 2 | 679 | 5.25 | **0.023** | 123.9 | 37.37 |
|  |  |  | Pool(Site type) | 6 | 170 | 0.87 | 0.567 | 0.0 | 0.00 |
|  |  |  | Month*Pool(Site type) | 10 | 125 | 0.64 | 0.791 | 0.0 | 0.00 |
|  |  |  | Error (i.e., Sample) | 10 | 196 |  |  | 195.7 | 59.00 |
|  |  | Species evenness | Site type | 1 | 16 | 0.09 | 0.765 | 0.0 | 0.00 |
|  |  |  | Month | 2 | 236 | 1.83 | 0.191 | 12.1 | 3.64 |
|  |  |  | Site type*Month | 2 | 679 | 5.25 | **0.025** | 123.9 | 37.37 |
|  |  |  | Pool(Site type) | 6 | 170 | 0.87 | 0.565 | 0.0 | 0.00 |
|  |  |  | Month*Pool(Site type) | 10 | 125 | 0.64 | 0.797 | 0.0 | 0.00 |
|  |  |  | Error (i.e., Sample) | 10 | 196 |  |  | 195.7 | 59.00 |
|  |  | Total number of individuals | Site type | 1 | 3620 | 1.44 | 0.285 | 46.3 | 1.29 |
|  |  |  | Month | 2 | 3031 | 1.37 | 0.267 | 51.3 | 1.43 |
|  |  |  | Site type*Month | 2 | 5151 | 2.33 | 0.082 | 367.7 | 10.24 |
|  |  |  | Pool(Site type) | 6 | 2509 | 0.80 | 0.625 | 0.0 | 0.00 |
|  |  |  | Month*Pool(Site type) | 12 | 2210 | 0.71 | 0.824 | 0.0 | 0.00 |
|  |  |  | Error (i.e., Sample) | 24 | 3125 |  |  | 3125.1 | 87.04 |

**Supplement 6. Similarity percentage (SIMPER) analyses for site type differences in visiting nekton communities, salt pool conditions and faunal communities, and prey assemblages of tomcods and mummichogs in Aulac and Wallace Bay, Maritime Canada**

**Table S6.1** SIMPER results for fish and invertebrate communities captured in creeks, on marsh platforms and in salt pools, and abiotic conditions in salt pools, identifying taxa or variables contributing most to significant site type differences detected by PERMANOVA, in reference (Ref) and restoring (Rest) salt marshes in Aulac and Wallace Bay captured from May–July in 2022–2023 (see Tables 2 and 3). Prior to analysis, taxa densities (number caught per fyke net, seine haul, minnow, or invertebrate activity trap) were 4^th^ root transformed and Bray-Curtis similarity used; abiotic conditions were normalized and Euclidean distance used. Juvenile life stage indicated by “(j)” and *Alosa* spp. are alewives (*A. aestivalis*) and blueback herring (*A. pseudoharengus*), and *Gasterosteus* spp. are threespine (*G. aculeatus*) and blackspotted (*G. wheatlandi*) sticklebacks. Dissimilarity/SD is the ratio of average dissimilarity to standard deviation of dissimilarities; a value > 1 indicates a contribution consistent among site types. Cut-off of 90% cumulative contribution used.

|  |  |  |  |  | Mean taxon density (4^th^ root no. individuals per sample) or variable (normalized per sample) | |  |  |  |
| --- | --- | --- | --- | --- | --- | --- | --- | --- | --- |
| Project | Subhabitat | Capture method or | Overall average dissimilarity or Average squared distance | Taxon | Site type | | Average dissimilarity | Dissimilarity/ SD | Contribution (%) |
|  |  |  |  |  | Ref | Rest |  |  |  |
| Aulac | Creek | Fyke net | 44.14 | *Anguilla rostrata* | 0.77 | 1.36 | 4.31 | 1.15 | 9.76 |
|  |  |  |  | *Fundulus heteroclitus* | 1.41 | 1.73 | 4.29 | 0.98 | 9.71 |
|  |  |  |  | *Pungitius pungitius* (j) | 1.40 | 0.33 | 3.99 | 0.99 | 9.03 |
|  |  |  |  | *Pungitius pungitius* | 0.92 | 0.20 | 3.71 | 1.14 | 8.41 |
|  |  |  |  | *Gasterosteus aculeatus* | 1.17 | 1.05 | 3.30 | 1.32 | 7.47 |
|  |  |  |  | *Gasterosteus wheatlandi* | 1.70 | 1.35 | 3.13 | 1.19 | 7.10 |
|  |  |  |  | *Anguilla rostrata* (j) | 0.28 | 0.56 | 3.11 | 0.87 | 7.05 |
|  |  |  |  | *Menidia menidia* | 0.67 | 0.51 | 3.05 | 1.09 | 6.90 |
|  |  |  |  | *Microgadus tomcod* | 2.43 | 2.62 | 2.90 | 1.15 | 6.57 |
|  |  |  |  | *Crangon septemspinosa* | 0.60 | 0.82 | 2.86 | 1.04 | 6.48 |
|  |  |  |  | *Gasterosteus aculeatus* (j) | 0.56 | 0.18 | 1.92 | 0.57 | 4.36 |
|  |  |  |  | *Fundulus heteroclitus* (j) | 0.42 | 0.59 | 1.55 | 0.68 | 3.52 |
|  |  |  |  | *Gasterosteus wheatlandi* (j) | 0.40 | 0.00 | 1.42 | 0.54 | 3.22 |
|  |  |  |  | *Microgadus tomcod* (j) | 0.23 | 0.08 | 1.27 | 0.56 | 2.87 |
|  | Platform | Seine net | 51.86 | *Menidia menidia* | 1.78 | 1.01 | 6.71 | 1.20 | 12.95 |
|  |  |  |  | *Microgadus tomcod* | 0.46 | 0.76 | 4.78 | 0.83 | 9.21 |
|  |  |  |  | *Gasterosteus wheatlandi* | 1.38 | 1.00 | 4.64 | 1.16 | 8.94 |
|  |  |  |  | *Osmerus mordax* (j) | 0.50 | 0.39 | 4.35 | 0.75 | 8.38 |
|  |  |  |  | *Apeltes quadracus* | 0.65 | 0.18 | 3.69 | 0.68 | 7.12 |
|  |  |  |  | *Fundulus heteroclitus* | 0.88 | 0.57 | 3.46 | 0.91 | 6.67 |
|  |  |  |  | *Pungitius pungitius* | 0.50 | 0.33 | 3.33 | 0.81 | 6.41 |
|  |  |  |  | *Anguilla rostrata* (j) | 0.43 | 0.22 | 2.56 | 0.66 | 4.93 |
|  |  |  |  | *Pungitius pungitius* (j) | 0.62 | 0.34 | 2.47 | 0.60 | 4.76 |
|  |  |  |  | *Fundulus diaphanus* | 0.26 | 0.00 | 1.95 | 0.49 | 3.76 |
|  |  |  |  | *Gasterosteus aculeatus* | 0.45 | 0.23 | 1.89 | 0.48 | 3.64 |
|  |  |  |  | *Fundulus heteroclitus* (j) | 0.19 | 0.22 | 1.79 | 0.54 | 3.46 |
|  |  |  |  | *Gasterosteus aculeatus* (j) | 0.18 | 0.18 | 1.73 | 0.56 | 3.34 |
|  |  |  |  | *Anguilla rostrata* | 0.18 | 0.17 | 1.65 | 0.50 | 3.18 |
|  |  |  |  | *Crangon septemspinosa* | 0.00 | 0.23 | 1.52 | 0.52 | 2.93 |
|  |  |  |  | *Gasterosteus wheatlandi* (j) | 0.22 | 0.00 | 1.49 | 0.44 | 2.87 |
|  | Salt pool | Abiotic conditions | 9.80 | Sedimet penetrability (cm) | -0.64 | 0.61 | 2.84 | 0.82 | 28.92 |
|  |  |  |  | Water depth (cm) | 0.53 | -0.51 | 2.40 | 0.67 | 24.48 |
|  |  |  |  | Water pH | 0.26 | -0.25 | 1.98 | 0.87 | 20.20 |
|  |  |  |  | Water dissolved oxygen (mg/L) | 0.01 | -0.01 | 1.28 | 0.91 | 13.05 |
|  |  |  |  | Water salinity (ppt) | -0.16 | 0.15 | 0.85 | 1.04 | 8.66 |
|  |  | Minnow trap | 86.59 | *Fundulus heteroclitus* | 0.45 | 0.99 | 37.95 | 1.14 | 43.83 |
|  |  |  |  | *Tritia* (*Ilyanassa*) *obsoleta* | 0.00 | 0.43 | 20.57 | 0.58 | 23.76 |
|  |  |  |  | *Gasterosteus aculeatus* | 0.25 | 0.17 | 10.33 | 0.50 | 11.93 |
|  |  |  |  | *Fundulus heteroclitus* (j) | 0.13 | 0.35 | 9.34 | 0.62 | 10.79 |
|  |  | Invertebrate activity trap | 100.00 | Hydrobiidae | 0.71 | 0.07 | 26.47 | 0.76 | 26.47 |
|  |  |  |  | Culicidae (larva) | 0.82 | 0.00 | 24.83 | 0.68 | 24.83 |
|  |  |  |  | Corixidae | 1.06 | 0.00 | 18.73 | 0.80 | 18.73 |
|  |  |  |  | Gammaridae | 0.68 | 0.00 | 15.28 | 0.68 | 15.28 |
|  |  |  |  | *Gasterosteus* spp. (j) | 0.32 | 0.00 | 10.85 | 0.47 | 10.85 |
| Wallace Bay | Creek | Fyke net | 43.41 | *Fundulus heteroclitus* | 4.30 | 2.39 | 6.00 | 1.07 | 13.82 |
|  |  |  |  | *Fundulus heteroclitus* (j) | 1.83 | 0.80 | 4.56 | 1.00 | 10.50 |
|  |  |  |  | *Crangon septemspinosa* | 1.47 | 2.72 | 4.41 | 1.64 | 10.15 |
|  |  |  |  | *Palaemon paludosus* | 0.76 | 1.29 | 3.02 | 1.33 | 6.96 |
|  |  |  |  | *Carcinus maenas* | 0.00 | 0.88 | 2.84 | 1.06 | 6.54 |
|  |  |  |  | *Morone americana* | 2.02 | 2.09 | 2.62 | 1.25 | 6.04 |
|  |  |  |  | *Rhithropanopeus harrisii* | 0.73 | 1.05 | 2.24 | 1.27 | 5.15 |
|  |  |  |  | *Menidia menidia* | 0.18 | 0.97 | 2.00 | 0.90 | 4.60 |
|  |  |  |  | *Apeltes quadracus* | 0.54 | 1.09 | 1.74 | 0.99 | 4.02 |
|  |  |  |  | *Gasterosteus wheatlandi* | 0.67 | 1.05 | 1.74 | 0.91 | 4.01 |
|  |  |  |  | *Pseudopleuronectes americanus* (j) | 0.00 | 0.64 | 1.69 | 0.94 | 3.89 |
|  |  |  |  | *Pungitius pungitius* | 0.52 | 0.28 | 1.66 | 0.56 | 3.82 |
|  |  |  |  | *Gasterosteus aculeatus* | 0.80 | 0.97 | 1.65 | 0.86 | 3.79 |
|  |  |  |  | *Osmerus mordax* | 0.00 | 0.47 | 0.97 | 0.65 | 2.24 |
|  |  |  |  | *Alosa* spp. | 0.08 | 0.13 | 0.85 | 0.41 | 1.97 |
|  |  |  |  | *Morone saxatilis* | 0.32 | 0.49 | 0.74 | 0.63 | 1.71 |
|  |  |  |  | *Panopeus herbstii* | 0.00 | 0.18 | 0.61 | 0.44 | 1.40 |
|  | Platform | Seine net | 64.20 | *Fundulus heteroclitus* (j) | 1.25 | 0.80 | 13.01 | 1.44 | 20.26 |
|  |  |  |  | *Fundulus heteroclitus* | 1.20 | 1.01 | 12.22 | 1.16 | 19.04 |
|  |  |  |  | *Menidia menidia* (j) | 0.06 | 0.99 | 10.54 | 0.66 | 16.42 |
|  |  |  |  | *Menidia menidia* | 0.18 | 0.72 | 7.30 | 0.73 | 11.37 |
|  |  |  |  | *Crangon septemspinosa* | 0.15 | 0.63 | 6.84 | 0.60 | 10.65 |
|  |  |  |  | *Gasterosteus aculeatus* | 0.17 | 0.29 | 4.84 | 0.64 | 7.53 |
|  |  |  |  | *Palaemon paludosus* | 0.06 | 0.22 | 3.24 | 0.52 | 5.05 |
|  | Salt pool | Abiotic conditions | 6.08 | Water depth (cm) | 0.13 | -0.13 | 2.55 | 0.89 | 41.93 |
|  |  |  |  | Sediment penetrability (cm) | 0.11 | -0.11 | 1.77 | 0.76 | 29.17 |
|  |  |  |  | Water pH | 0.06 | -0.06 | 1.34 | 0.75 | 22.05 |
|  |  | Minnow trap | 61.30 | *Fundulus heteroclitus* | 1.85 | 1.36 | 21.92 | 1.09 | 35.76 |
|  |  |  |  | *Tritia* (*Ilyanassa*) *obsoleta* | 0.00 | 0.69 | 10.81 | 0.80 | 17.63 |
|  |  |  |  | *Fundulus heteroclitus* (j) | 0.74 | 0.86 | 9.61 | 0.90 | 15.68 |
|  |  |  |  | *Crangon septemspinosa* | 0.00 | 0.22 | 4.15 | 0.46 | 6.77 |
|  |  |  |  | *Gasterosteus aculeatus* | 0.09 | 0.13 | 3.16 | 0.46 | 5.15 |
|  |  |  |  | *Palaemon paludosus* | 0.00 | 0.19 | 2.70 | 0.44 | 4.41 |
|  |  |  |  | *Pungitius pungitius* | 0.13 | 0.04 | 2.26 | 0.40 | 3.69 |
|  |  |  |  | *Carcinus maenas* | 0.00 | 0.17 | 2.15 | 0.43 | 3.50 |
|  |  | Invertebrate activity trap | 82.47 | Hydrobiidae | 0.62 | 0.35 | 32.74 | 0.85 | 39.70 |
|  |  |  |  | Gammaridae | 0.28 | 0.18 | 12.36 | 0.58 | 14.99 |
|  |  |  |  | Culicidae (larva) | 0.04 | 0.10 | 9.33 | 0.35 | 11.31 |
|  |  |  |  | *Fundulus heteroclitus* (j) | 0.25 | 0.00 | 7.25 | 0.43 | 8.79 |
|  |  |  |  | Corixidae | 0.29 | 0.00 | 7.23 | 0.45 | 8.76 |
|  |  |  |  | *Gasterosteus* spp. (j) | 0.05 | 0.00 | 4.44 | 0.21 | 5.39 |
|  |  |  |  | Mysida | 0.04 | 0.01 | 4.26 | 0.31 | 5.16 |

**Table S6.2** SIMPER results for prey assemblage in tomcods (*Microgadus tomcod*) and mummichogs (*Fundulus heteroclitus*), identifying prey items contributing most to significant site type differences detected by PERMANOVA, in fish captured in creeks and on platforms in reference (Ref) and restoring (Rest) salt marshes in Aulac and Wallace Bay in May–July in 2022–2023 (see Table 4). Prey composition data (number per gut item) were converted to presence-absence prior to analysis; this was done to include vegetal matter (detritus and macroalgae) in the analysis. Dissimilarity/SD is the ratio of average dissimilarity to standard deviation of dissimilarities; a value > 1 indicates a contribution consistent among site types. Cut-off of 90% cumulative contribution used.

|  |  |  |  |  | Mean gut item occurrence (presence-absence no. occurrences per sample) | |  |  |  |
| --- | --- | --- | --- | --- | --- | --- | --- | --- | --- |
| Project | Species | Subhabitat | Average Dissimilarity | Gut item | Site type | | Average dissimilarity | Dissimilarity/ SD | Contribution (%) |
|  |  |  |  |  | Ref | Rest |  |  |  |
| Aulac | Tomcod | Creek | 50.02 | Nereididae | 0.63 | 0.57 | 8.92 | 0.90 | 17.83 |
|  |  |  |  | Gammaridae | 0.68 | 0.21 | 8.81 | 0.99 | 17.62 |
|  |  |  |  | Detritus | 0.46 | 0.29 | 5.98 | 0.86 | 11.96 |
|  |  |  |  | *Crangon septemspinosa* | 0.24 | 0.29 | 4.23 | 0.65 | 8.46 |
|  |  |  |  | Macroalgae | 0.15 | 0.10 | 2.55 | 0.50 | 5.10 |
|  |  |  |  | Mysida | 0.32 | 0.31 | 2.34 | 0.47 | 4.69 |
|  |  |  |  | Diptera other (larva) | 0.07 | 0.10 | 2.09 | 0.44 | 4.17 |
|  |  |  |  | Polychaeta other | 0.05 | 0.12 | 2.01 | 0.39 | 4.02 |
|  |  |  |  | Gasterosteidae | 0.12 | 0.00 | 1.92 | 0.36 | 3.84 |
|  |  |  |  | *Corophium volutator* | 0.93 | 0.95 | 1.86 | 0.31 | 3.73 |
|  |  |  |  | Nematoda | 0.10 | 0.00 | 1.40 | 0.33 | 2.79 |
|  |  |  |  | *Fundulus heteroclitus* | 0.05 | 0.05 | 1.27 | 0.32 | 2.53 |
|  |  |  |  | *Limecola* | 0.00 | 0.12 | 1.22 | 0.34 | 2.45 |
|  |  |  |  | Hemiptera | 0.07 | 0.00 | 0.95 | 0.27 | 1.70 |
|  | Mummichog | Creek | 88.07 | Gammaridae | 0.59 | 0.22 | 11.92 | 0.84 | 13.53 |
|  |  |  |  | Nereididae | 0.59 | 0.22 | 11.76 | 0.88 | 13.36 |
|  |  |  |  | Chironomidae (adult) | 0.12 | 0.39 | 8.35 | 0.80 | 9.48 |
|  |  |  |  | Detritus | 0.29 | 0.17 | 6.16 | 0.59 | 7.00 |
|  |  |  |  | Diptera other (larva) | 0.06 | 0.28 | 5.97 | 0.61 | 6.78 |
|  |  |  |  | *Corophium volutator* | 0.12 | 0.28 | 5.65 | 0.63 | 6.42 |
|  |  |  |  | Nematoda | 0.24 | 0.06 | 5.60 | 0.56 | 6.36 |
|  |  |  |  | Hemiptera | 0.24 | 0.00 | 3.82 | 0.44 | 4.34 |
|  |  |  |  | Diptera other (adult) | 0.12 | 0.06 | 3.72 | 0.39 | 4.22 |
|  |  |  |  | Polychaeta other | 0.00 | 0.17 | 3.28 | 0.43 | 3.72 |
|  |  |  |  | Copepoda | 0.06 | 0.11 | 2.92 | 0.43 | 3.31 |
|  |  |  |  | Hymenoptera | 0.06 | 0.06 | 2.46 | 0.30 | 2.79 |
|  |  |  |  | Isopoda | 0.06 | 0.11 | 2.35 | 0.43 | 2.67 |
|  |  |  |  | Coleoptera | 0.06 | 0.11 | 2.15 | 0.41 | 2.44 |
|  |  |  |  | Platyhelminthes | 0.06 | 0.00 | 1.70 | 0.28 | 1.93 |
|  |  | Platform | 82.13 | Nereididae | 0.67 | 0.19 | 15.05 | 0.81 | 18.32 |
|  |  |  |  | Gammaridae | 0.13 | 0.44 | 8.59 | 0.83 | 10.46 |
|  |  |  |  | Diptera other (adult) | 0.33 | 0.19 | 7.49 | 0.69 | 9.11 |
|  |  |  |  | Detritus | 0.13 | 0.25 | 5.49 | 0.58 | 6.68 |
|  |  |  |  | Hemiptera | 0.07 | 0.19 | 5.09 | 0.52 | 6.20 |
|  |  |  |  | *Corophium volutator* | 0.27 | 0.31 | 4.84 | 0.48 | 5.89 |
|  |  |  |  | Egg (fish or invertebrate) | 0.00 | 0.19 | 4.30 | 0.46 | 5.23 |
|  |  |  |  | Hydrobiidae | 0.07 | 0.25 | 4.15 | 0.53 | 5.05 |
|  |  |  |  | Macroalgae | 0.07 | 0.13 | 3.52 | 0.47 | 4.28 |
|  |  |  |  | Diptera other (larva) | 0.07 | 0.13 | 3.42 | 0.49 | 4.17 |
|  |  |  |  | Polychaeta other | 0.00 | 0.19 | 2.72 | 0.45 | 3.31 |
|  |  |  |  | Coleoptera | 0.07 | 0.13 | 2.33 | 0.37 | 2.83 |
|  |  |  |  | Foraminifera | 0.00 | 0.13 | 2.30 | 0.39 | 2.80 |
|  |  |  |  | Nematoda | 0.07 | 0.06 | 2.29 | 0.39 | 2.79 |
|  |  |  |  | Hymenoptera | 0.07 | 0.13 | 1.95 | 0.35 | 2.37 |
|  |  |  |  | Corixidae | 0.00 | 0.13 | 1.88 | 0.39 | 2.29 |
| Wallace Bay | Mummichog | Creek | 86.92 | Mysida | 0.23 | 0.65 | 21.92 | 1.38 | 25.21 |
|  |  |  |  | Detritus | 0.23 | 0.55 | 11.98 | 0.98 | 13.79 |
|  |  |  |  | *Crangon septemspinosa* | 0.23 | 0.55 | 11.75 | 0.90 | 13.51 |
|  |  |  |  | Macroalgae | 0.05 | 0.40 | 9.36 | 0.70 | 10.76 |
|  |  |  |  | Nereididae | 0.27 | 0.20 | 7.31 | 0.71 | 8.41 |
|  |  |  |  | Gammaridae | 0.00 | 0.10 | 1.66 | 0.31 | 1.91 |
|  |  |  |  | Acariformes | 0.09 | 0.00 | 1.61 | 0.29 | 1.85 |
|  |  |  |  | Diptera other (larva) | 0.05 | 0.05 | 1.42 | 0.29 | 1.64 |
|  |  |  |  | Platyhelminthes | 0.05 | 0.00 | 1.42 | 0.21 | 1.63 |
|  |  | Platform | 74.18 | Detritus | 0.67 | 0.60 | 10.06 | 0.86 | 13.56 |
|  |  |  |  | Macroalgae | 0.50 | 0.67 | 7.15 | 0.98 | 9.63 |
|  |  |  |  | Hemiptera | 0.67 | 0.53 | 5.89 | 0.85 | 7.93 |
|  |  |  |  | Copepoda | 0.28 | 0.33 | 5.65 | 0.92 | 7.61 |
|  |  |  |  | Diptera other (larva) | 0.33 | 0.33 | 5.29 | 0.87 | 7.13 |
|  |  |  |  | Foraminifera | 0.39 | 0.07 | 4.34 | 0.73 | 5.85 |
|  |  |  |  | Acariformes | 0.28 | 0.13 | 4.13 | 0.75 | 5.57 |
|  |  |  |  | Diptera other (adult) | 0.44 | 0.20 | 4.05 | 0.66 | 5.46 |
|  |  |  |  | Hydrobiidae | 0.22 | 0.13 | 3.23 | 0.65 | 4.35 |
|  |  |  |  | Ostracoda | 0.33 | 0.00 | 3.09 | 0.56 | 4.17 |
|  |  |  |  | Nereididae | 0.06 | 0.20 | 2.47 | 0.49 | 3.33 |
|  |  |  |  | Gammaridae | 0.17 | 0.00 | 2.14 | 0.48 | 2.88 |
|  |  |  |  | Polychaeta other | 0.00 | 0.20 | 2.06 | 0.48 | 2.78 |
|  |  |  |  | Hymenoptera | 0.11 | 0.07 | 1.81 | 0.39 | 2.44 |
|  |  |  |  | Odonata (larva) | 0.11 | 0.07 | 1.68 | 0.40 | 2.26 |
|  |  |  |  | Mysida | 0.11 | 0.00 | 1.62 | 0.38 | 2.18 |
|  |  |  |  | Coleoptera | 0.11 | 0.00 | 1.52 | 0.38 | 2.06 |
|  |  |  |  | *Melampus bidentatus* | 0.11 | 0.00 | 1.28 | 0.38 | 1.73 |

**Supplement 7. Additional PERMANOVA results for within-year differences in platform nekton communities, or salt pool conditions and small-bodied faunal communities between reference and restoring salt marshes in Aulac and Wallace Bay, Maritime Canada**

**Table S7.1** PERMANOVA results for within-year differences in platform nekton communities, salt pool abiotic conditions, and/or minnow and invertebrate activity trap communities in reference and restoring salt marshes in Aulac and Wallace Bay in 2022–2023; this table complements Tables 2 and 3. See Table S1 for sampling dates, Table 1 for taxa included in each biotic community, and Table S8.1 for abiotic variables measured in salt pools. Taxa densities 4^th^ root transformed, and abiotic variables normalized prior to analysis. Site type and Month are fixed factors, and Pool is a random factor. P-values bolded for significant and interpretable fixed effects. When the number of permutations is low (< 100), P-values obtained by Monte Carlo simulations (P(MC)) should be used. PERMDISP tests conducted for significant fixed effects to assess amount of multivariate dispersion among groups; df1 and df2 represent the numerator and denominator degrees of freedom for the F-ratio, respectively. Analysis of components of variation conducted for both fixed and random effects to evaluate relative importance of spatiotemporal scales.

| Project | Community | Year | PERMANOVA | | | | | | | Components of variation | | PERMDISP | | |
| --- | --- | --- | --- | --- | --- | --- | --- | --- | --- | --- | --- | --- | --- | --- |
|  |  |  | Source of variation | df | MS | Pseudo-F | P(perm) | Unique permutations | P(MC) | Estimate | % | F | df 1, df 2 | P |
| Aulac | Salt pool abiotic variables | 2022 | Site type | 1 | 11.3 | 4.62 | 0.022 | 35 | 0.020 | 0.7 | 12.9 | 8.74 | 1, 22 | 0.012 |
|  |  |  | Month | 2 | 21.1 | 16.26 | 0.001 | 999 | 0.001 | 2.5 | 43.0 | 1.10 | 2, 21 | 0.400 |
|  |  |  | Site type*Month | 2 | 4.7 | 3.64 | **0.003** | 999 | 0.006 | 0.9 | 14.9 | 5.68 | 5, 18 | 0.010 |
|  |  |  | Pool(Site type) | 6 | 2.5 | 1.89 | 0.034 | 998 | 0.048 | 0.4 | 6.7 |  |  |  |
|  |  |  | Month*Pool (Site type) (i.e., Sample) | 12 |  |  |  |  |  | 1.3 | 22.6 |  |  |  |
|  |  | 2023 | Site type | 1 | 25.1 | 8.36 | 0.008 | 275 | 0.004 | 2.0 | 17.4 | 58.86 | 1, 21 | 0.001 |
|  |  |  | Month | 2 | 33.6 | 10.84 | 0.001 | 999 | 0.001 | 4.1 | 35.8 | 0.95 | 2, 20 | 0.611 |
|  |  |  | Site type*Month | 2 | 11.6 | 3.73 | **0.003** | 999 | 0.012 | 2.3 | 19.9 | 9.04 | 5, 17 | 0.016 |
|  |  |  | Pool(Site type) | 6 | 3.0 | 0.97 | 0.489 | 999 | 0.490 | 0.0 | 0.0 |  |  |  |
|  |  |  | Month*Pool(Site type) (i.e., Sample) | 11 | 3.1 |  |  |  |  | 3.1 | 27.0 |  |  |  |
| Wallace Bay | Platform nekton community | 2022 | Site type | 1 | 6079 | 4.29 | 0.027 | 998 | 0.026 | 517.8 | 13.6 | 0.41 | 1, 16 | 0.588 |
|  |  |  | Month | 2 | 7599 | 5.36 | 0.001 | 999 | 0.001 | 1030.1 | 27.0 | 0.20 | 2, 15 | 0.842 |
|  |  |  | Site type*Month | 2 | 3945 | 2.78 | **0.039** | 999 | 0.030 | 842.1 | 22.1 | 2.97 | 5, 12 | 0.301 |
|  |  |  | Error (i.e., Net) | 12 | 1419 |  |  |  |  | 1418.6 | 37.2 |  |  |  |
|  |  | 2023 | Site type | 1 | 4764 | 5.97 | 0.003 | 999 | 0.006 | 440.7 | 13.5 | 6.07 | 1, 16 | 0.094 |
|  |  |  | Month | 2 | 4633 | 5.81 | 0.001 | 999 | 0.001 | 639.3 | 19.5 | 0.62 | 2, 15 | 0.645 |
|  |  |  | Site type*Month | 2 | 4985 | 6.25 | **0.001** | 998 | 0.002 | 1395.8 | 42.6 | 7.44 | 5, 12 | 0.032 |
|  |  |  | Error (i.e., Net) | 12 | 797 |  |  |  |  | 797.4 | 24.4 |  |  |  |
|  | Salt pool minnow trap community | 2022 | Site type | 1 | 16272 | 17.48 | 0.044 | 35 | 0.002 | 1278.4 | 31.8 | 35.45 | 1, 22 | 0.001 |
|  |  |  | Month | 2 | 6503 | 7.00 | 0.001 | 998 | 0.002 | 696.7 | 17.3 | 25.86 | 2, 21 | 0.001 |
|  |  |  | Site type*Month | 2 | 5410 | 5.82 | **0.001** | 999 | 0.001 | 1120.1 | 27.8 | 6.11 | 5, 18 | 0.010 |
|  |  |  | Pool(Site type) | 6 | 931 | 1.00 | 0.511 | 998 | 0.485 | 0.6 | 0.0 |  |  |  |
|  |  |  | Month*Pool(Site type) (i.e., Sample) | 12 | 929 |  |  |  |  | 929.4 | 23.1 |  |  |  |
|  |  | 2023 | Site type | 1 | 2880 | 2.63 | **0.023** | 35 | 0.065 | 148.9 | 6.4 | 0.04 | 1, 22 | 0.876 |
|  |  |  | Month | 2 | 3154 | 1.71 | 0.129 | 998 | 0.146 | 164.0 | 7.1 |  |  |  |
|  |  |  | Site type*Month | 2 | 2496 | 1.36 | 0.271 | 998 | 0.260 | 163.7 | 7.1 |  |  |  |
|  |  |  | Pool(Site) | 6 | 1093 | 0.59 | 0.921 | 997 | 0.875 | 0.0 | 0.0 |  |  |  |
|  |  |  | Month*Pool(Site type) (i.e., Sample) | 12 | 1842 |  |  |  |  | 1841.7 | 79.4 |  |  |  |
|  | Salt pool invertebrate activity trap community | 2022 | Site type | 1 | 1944 | 0.86 | 0.563 | 35 | 0.505 | 0.0 | 0.0 |  |  |  |
|  |  |  | Month | 2 | 7942 | 2.51 | 0.029 | 998 | 0.036 | 596.9 | 11.5 | 0.49 | 2, 21 | 0.705 |
|  |  |  | Site type*Month | 2 | 8868 | 2.80 | 0.016 | 999 | 0.016 | 1425.1 | 27.5 | 8.47 | 5, 18 | 0.011 |
|  |  |  | Pool(Site type) | 6 | 2267 | 0.72 | 0.843 | 999 | 0.810 | 0.0 | 0.0 |  |  |  |
|  |  |  | Month*Pool(Site type) (i.e., Sample) | 12 | 3167 |  |  |  |  | 3167.2 | 61.0 |  |  |  |
|  |  | 2023 | Site type | 1 | 5824 | 3.60 | 0.061 | 35 | 0.040 | 350.4 | 8.2 | 1.05 | 1, 22 | 0.379 |
|  |  |  | Month | 2 | 4706 | 1.37 | 0.255 | 997 | 0.233 | 159.7 | 3.7 |  |  |  |
|  |  |  | Site type*Month | 2 | 4803 | 1.40 | 0.244 | 997 | 0.248 | 343.6 | 8.0 |  |  |  |
|  |  |  | Pool(Site type) | 6 | 1619 | 0.47 | 0.928 | 998 | 0.921 | 0.0 | 0.0 |  |  |  |
|  |  |  | Month*Pool(Site type) (i.e., Sample) | 12 | 3428 |  |  |  |  | 3428.1 | 80.1 |  |  |  |

**Supplement 8. Abiotic conditions in salt pools and the surrounding estuary, as well as vegetation cover in salt pools in Aulac and Wallace Bay, Maritime Canada**

**Table S8.1** Mean ± SE of the abiotic conditions in reference salt pools and restoring pools in the Aulac and Wallace Bay restoration projects. Measurements were taken in pools in the evening at low tide in May–July in 2022–2023 (see Table S1 for sampling dates). Four pools were sampled in each site type during each round. No water measurements were recorded in the Aulac restoring site in May 2023 due to malfunctioning equipment.

| Project | Year | Month | Site type | Sediment penetrability (cm) | Water depth (cm) | pH | Salinity (ppt) | Temperature  (°C) | Dissolved oxygen (mg/L) |
| --- | --- | --- | --- | --- | --- | --- | --- | --- | --- |
| Aulac | 2022 | May | Reference | 27.4±2.7 | 22.3±1.6 | 7.70±0.12 | 6.03±0.82 | 10.85±0.80 | 6.73±0.45 |
|  |  |  | Restoring | 35.7±3.7 | 13.8±2.8 | 7.72±0.08 | 15.70±0.47 | 15.30±1.37 | 8.00±1.14 |
|  |  | June | Reference | 27.3±3.2 | 28.0±4.4 | 8.05±0.22 | 24.81±0.98 | 17.63±0.69 | 5.35±0.51 |
|  |  |  | Restoring | 35.3±4.1 | 20.4±1.9 | 7.60±0.04 | 17.03±0.98 | 16.13±0.14 | 6.95±0.44 |
|  |  | July | Reference | 25.4±4.3 | 28.9±4.5 | 7.62±0.12 | 27.48±0.22 | 22.33±0.59 | 3.45±0.21 |
|  |  |  | Restoring | 36.2±2.2 | 18.4±3.3 | 7.76±0.04 | 28.78±0.32 | 22.00±0.26 | 3.85±0.09 |
|  | 2023 | May | Reference | 14.4±3.4 | 29.1±8.0 | 8.50±0.09 | 25.25±0.50 | 15.00±0.18 | 11.48±0.70 |
|  |  |  | Restoring | 37.8±2.4 | 15.4±1.6 | n/a | n/a | n/a | n/a |
|  |  | June | Reference | 20.9±8.8 | 33.5±8.2 | 8.10±0.17 | 7.74±1.54 | 25.48±0.34 | 6.41±1.47 |
|  |  |  | Restoring | 35.5±1.8 | 20.1±0.5 | 7.81±0.03 | 18.54±0.53 | 24.90±0.51 | 7.53±0.44 |
|  |  | July | Reference | 27.5±3.0 | 30.1±9.6 | 7.17±0.17 | 14.65±1.48 | 26.23±1.13 | 6.70±2.12 |
|  |  |  | Restoring | 35.2±2.2 | 18.8±4.5 | 7.63±0.03 | 11.59±0.05 | 20.58±0.06 | 6.15±0.49 |
| Wallace Bay | 2022 | May | Reference | 35.0±2.3 | 31.5±4.1 | 7.74±0.16 | 15.96±0.31 | 13.15±0.25 | 7.93±0.30 |
|  |  |  | Restoring | 33.6±0.9 | 18.9±1.2 | 8.01±0.21 | 18.04±0.31 | 13.33±0.43 | 8.05±0.18 |
|  |  | June | Reference | 28.3±9.7 | 23.5±4.1 | 8.14±0.13 | 23.23±0.24 | 17.90±0.29 | 8.05±0.09 |
|  |  |  | Restoring | 26.3±4.3 | 21.3±5.0 | 8.38±0.03 | 24.22±0.13 | 18.33±0.28 | 7.23±0.20 |
|  |  | July | Reference | 22.8±4.0 | 28.0±2.1 | 7.55±0.14 | 24.50±0.24 | 23.03±0.14 | 3.08±0.33 |
|  |  |  | Restoring | 32.3±3.3 | 23.0±1.9 | 7.62±0.06 | 24.97±0.17 | 22.28±0.09 | 3.63±1.34 |
|  | 2023 | May | Reference | 25.8±4.1 | 24.6±4.0 | 7.88±0.10 | 0.16±0.002 | 7.95±0.24 | 11.53±0.44 |
|  |  |  | Restoring | 19.8±5.8 | 17.9±3.7 | 7.77±0.04 | 0.19±0.001 | 7.73±0.25 | 10.73±0.10 |
|  |  | June | Reference | 25.4±5.0 | 27.0±3.8 | 7.65±0.24 | 15.48±1.69 | 16.05±0.12 | 8.04±0.72 |
|  |  |  | Restoring | 18.6±4.6 | 15.8±3.1 | 7.35±0.05 | 14.77±0.10 | 14.83±0.09 | 6.57±0.41 |
|  |  | July | Reference | 25.5±5.8 | 26.6±3.5 | 7.85±0.37 | 16.93±0.14 | 29.43±0.14 | 5.38±0.82 |
|  |  |  | Restoring | 18.8±5.0 | 13.7±0.5 | 7.71±0.12 | 18.30±0.37 | 29.48±0.23 | 4.77±0.30 |

**Table S8.2** Mean ± SE of representative surface water conditions in the adjacent estuary of the Aulac and Wallace Bay restoration projects. Measurements were taken during the daytime high tide in May–July in 2022–2023 (see Table S1 for sampling dates). Between three to four locations in the main salt marsh channel or in the bay were selected for taking these abiotic measurements.

| Project | Waterbody | Year | Month | pH | Salinity (ppt) | Temperature  (°C) | Dissolved oxygen (mg/L) |
| --- | --- | --- | --- | --- | --- | --- | --- |
| Aulac | Bay of Fundy | 2022 | May | 7.57±0.05 | 25.98±0.10 | 11.83±0.38 | 9.53±0.10 |
|  |  |  | June | 7.88±0.01 | 26.51±0.20 | 18.63±0.19 | 6.93±0.07 |
|  |  |  | July | 7.94±0.006 | 28.48±0.10 | 21.48±0.12 | 2.83±0.05 |
|  |  | 2023 | May | 7.72±0.02 | 25.99±0.05 | 11.83±0.34 | 9.86±0.09 |
|  |  |  | June | 7.69±0.02 | 21.24±0.07 | 20.10±0.06 | 5.82±0.20 |
|  |  |  | July | 7.79±0.01 | 16.02±0.07 | 27.68±0.75 | 5.43±0.06 |
| Wallace Bay | Northumberland Strait | 2022 | May | 7.89±0.03 | 13.86±0.10 | 11.65±0.15 | 9.05±0.06 |
|  |  |  | June | 8.11±0.004 | 24.89±0.02 | 15.15±0.09 | 6.90±0.07 |
|  |  |  | July | 7.92±0.01 | 24.44±0.01 | 22.90±0.00 | 2.68±0.06 |
|  |  | 2023 | May | 7.89±0.09 | 0.20±0.005 | 7.83±0.44 | 11.83±0.09 |
|  |  |  | June | 7.55±0.05 | 13.49±0.70 | 14.60±0.18 | 8.20±0.10 |
|  |  |  | July | 7.44±0.01 | 18.25±0.05 | 25.88±0.09 | 4.53±0.20 |

**Table S8.3** Mean ± SE of *Ruppia maritima* and green macroalgae percent cover in salt pools and creek pools in reference and restoring salt marshes in Aulac and Wallace Bay in 2022–2023 (see Table S1 for sampling dates). Vegetation cover was recorded in four pools in each site type during each round except for the Aulac reference site in July 2023 where only three pools were sampled.

| Project | Site type | Year | Month | *Ruppia maritima* (%) | Green macroalgae (%) |
| --- | --- | --- | --- | --- | --- |
| Aulac | Reference | 2022 | May | 0 | 0 |
|  |  |  | June | 50.0±15.9 | 26.5±7.5 |
|  |  |  | July | 79.3±9.7 | 18.9±9.3 |
|  |  | 2023 | May | 30.3±11.1 | 8.25±4.2 |
|  |  |  | June | 15.3±10.5 | 30.6±10.5 |
|  |  |  | July | 5.0±2.9 | 20.8±7.1 |
|  | Restoring | 2022 | May | 0 | 0 |
|  |  |  | June | 0 | 0 |
|  |  |  | July | 0 | 3.1±2.4 |
|  |  | 2023 | May | 0 | 0 |
|  |  |  | June | 0 | 0 |
|  |  |  | July | 0 | 0 |
| Wallace Bay | Reference | 2022 | May | 0 | 0 |
|  |  |  | June | 8.8±5.2 | 18.1±5.0 |
|  |  |  | July | 41.3±8.8 | 22.5±3.1 |
|  |  | 2023 | May | 1.8±1.8 | 18.3±15.8 |
|  |  |  | June | 37.5±6.6 | 20.7±1.7 |
|  |  |  | July | 73.8±8.5 | 63.8±12.8 |
|  | Restoring | 2022 | May | 0 | 0.6±0.6 |
|  |  |  | June | 0 | 22.5±4.2 |
|  |  |  | July | 23.8±17.1 | 3.4±1.0 |
|  |  | 2023 | May | 0 | 3.1±2.4 |
|  |  |  | June | 0 | 2.0±0.9 |
|  |  |  | July | 0 | 2.5±1.0 |

**Table S8.4** Mixed-model PERMANOVA results for percent cover of aquatic vegetation (*Ruppia maritima* and green macroalgae) in salt pools and creek pools in reference and restoring salt marshes in Aulac and Wallace Bay in 2022–2023. Site type, Year and Month are fixed factors, and Pool is a random factor. Vegetation cover was compared between site types in June and July of both years; May of both years was omitted from the analysis because lack of vegetation likely reflected that it was too early in the growing season. Data missing for a reference salt pool in Aulac in July 2023. See Table S1 for sampling dates. P-values bolded for significant and interpretable fixed effects involving Site type; 908–999 unique permutations. Analysis of components of variation conducted for both fixed and random effects to evaluate relative importance of spatiotemporal scales.

| Community | PERMANOVA |  |  |  |  | Components of Variation | |
| --- | --- | --- | --- | --- | --- | --- | --- |
|  | Source | df | MS | Pseudo-F | P(perm) | Estimate | % |
| Aulac pool vegetation | Site type | 1 | 13338 | 28.46 | 0.001 | 916.0 | 34.2 |
|  | Year | 1 | 5225 | 11.15 | 0.003 | 338.5 | 12.7 |
|  | Month | 1 | 303 | 0.81 | 0.443 | 0.0 | 0.0 |
|  | Site type*Year | 1 | 5236 | 11.17 | **0.006** | 678.7 | 25.4 |
|  | Site type*Month | 1 | 421 | 1.12 | 0.320 | 6.5 | 0.2 |
|  | Year*Month | 1 | 725 | 1.93 | 0.189 | 49.9 | 1.9 |
|  | Site type*Year*Month | 1 | 692 | 1.84 | 0.171 | 90.2 | 3.4 |
|  | Pool(Site type*Year) | 12 | 493 | 6.31 | 0.295 | 222.3 | 8.3 |
|  | Month*Pool(Site type*Year) (i.e., Sample) | 10 | 374 |  |  | 373.9 | 14.0 |
| Wallace Bay pool vegetation | Site type | 1 | 13936 | 51.00 | 0.001 | 853.9 | 29.9 |
|  | Year | 1 | 955 | 3.49 | 0.046 | 42.6 | 1.5 |
|  | Month | 1 | 4693 | 10.17 | 0.004 | 264.5 | 9.3 |
|  | Site type*Year | 1 | 5737 | 21.00 | **0.001** | 683.0 | 24.0 |
|  | Site type*Month | 1 | 3195 | 6.92 | **0.005** | 341.7 | 12.0 |
|  | Year*Month | 1 | 1900 | 4.12 | 0.040 | 179.9 | 6.3 |
|  | Site type*Year*Month | 1 | 560 | 1.21 | 0.307 | 24.6 | 0.9 |
|  | Pool(Site type*Year) | 12 | 273 | 0.59 | 0.895 | 0.0 | 0.0 |
|  | Month*Pool(Site type*Year) (i.e., Sample) | 12 | 461 |  |  | 461.3 | 16.2 |

**Supplement 9. Faunal densities in invertebrate activity traps in salt pools or creek pools in Aulac and Wallace Bay, Maritime Canada**

**Table S9.1** Mean ± SE densities of juvenile fish and invertebrate taxa captured in reference salt pools and restoring pools in Aulac and Wallace Bay salt marshes over three months (May–July) in 2022–2023. Averaged over 4 invertebrate activity traps per sampling round per site type (*n* = 4). Units are numbers of individuals per trap.

|  | **Aulac** | | | | | | | | | | | |
| --- | --- | --- | --- | --- | --- | --- | --- | --- | --- | --- | --- | --- |
|  | **Reference** | | | | | | **Restoring** | | | | | |
| Taxa | May 2022 | May 2023 | June 2022 | June 2023 | July 2022 | July 2023 | May 2022 | May 2023 | June 2022 | June 2023 | July 2022 | July 2023 |
| *Fundulus heteroclitus* | 0.0 ± 0.0 | 0.0 ± 0.0 | 0.0 ± 0.0 | 0.0 ± 0.0 | 0.50 ± 0.29 | 0.67 ± 0.67 | 0.0 ± 0.0 | 0.0 ± 0.0 | 0.0 ± 0.0 | 0.0 ± 0.0 | 0.0 ± 0.0 | 0.0 ± 0.0 |
| Gasterosteidae | 0.0 ± 0.0 | 0.0 ± 0.0 | 0.50 ± 0.29 | 0.25 ± 0.25 | 0.75 ± 0.25 | 1.0 ± 1.0 | 0.0 ± 0.0 | 0.0 ± 0.0 | 0.0 ± 0.0 | 0.0 ± 0.0 | 0.0 ± 0.0 | 0.0 ± 0.0 |
| *Crangon septemspinosa* | 0.0 ± 0.0 | 0.0 ± 0.0 | 0.0 ± 0.0 | 0.0 ± 0.0 | 0.0 ± 0.0 | 0.0 ± 0.0 | 0.0 ± 0.0 | 0.0 ± 0.0 | 0.0 ± 0.0 | 0.0 ± 0.0 | 0.0 ± 0.0 | 0.0 ± 0.0 |
| Mysida | 0.0 ± 0.0 | 0.0 ± 0.0 | 0.0 ± 0.0 | 0.0 ± 0.0 | 0.0 ± 0.0 | 0.0 ± 0.0 | 0.0 ± 0.0 | 0.0 ± 0.0 | 0.0 ± 0.0 | 0.0 ± 0.0 | 0.0 ± 0.0 | 0.0 ± 0.0 |
| Gammaridae | 1.25 ± 0.75 | 0.0 ± 0.0 | 0.50 ± 0.50 | 0.25 ± 0.25 | 2.25 ± 0.63 | 21.67 ± 7.88 | 0.0 ± 0.0 | 0.0 ± 0.0 | 0.0 ± 0.0 | 0.0 ± 0.0 | 0.0 ± 0.0 | 0.0 ± 0.0 |
| *Corophium volutator* | 0.0 ± 0.0 | 0.0 ± 0.0 | 0.0 ± 0.0 | 0.0 ± 0.0 | 0.0 ± 0.0 | 0.0 ± 0.0 | 0.0 ± 0.0 | 0.0 ± 0.0 | 0.0 ± 0.0 | 0.0 ± 0.0 | 0.0 ± 0.0 | 0.25 ± 0.25 |
| Hydrobiidae | 1.75 ± 1.75 | 3.00 ± 2.38 | 0.5 ± 0.5 | 8.00 ± 3.34 | 19.25 ± 18.59 | 0.0 ± 0.0 | 0.0 ± 0.0 | 0.0 ± 0.0 | 0.0 ± 0.0 | 0.0 ± 0.0 | 0.0 ± 0.0 | 2.50 ± 2.50 |
| Corixidae | 0.0 ± 0.0 | 0.0 ± 0.0 | 0.25 ± 0.25 | 8.25 ± 7.28 | 160.25 ± 116.40 | 56.33 ± 29.38 | 0.0 ± 0.0 | 0.0 ± 0.0 | 0.0 ± 0.0 | 0.0 ± 0.0 | 0.0 ± 0.0 | 0.0 ± 0.0 |
| Culicidae larva | 76.50 ± 30.31 | 13.75 ± 8.3 | 0.0 ± 0.0 | 1.75 ± 1.44 | 0.0 ± 0.0 | 0.0 ± 0.0 | 0.0 ± 0.0 | 0.0 ± 0.0 | 0.0 ± 0.0 | 0.0 ± 0.0 | 0.0 ± 0.0 | 0.0 ± 0.0 |
| Other Diptera larva | 0.0 ± 0.0 | 0.0 ± 0.0 | 0.0 ± 0.0 | 0.0 ± 0.0 | 0.25 ± 0.25 | 0.0 ± 0.0 | 0.0 ± 0.0 | 0.0 ± 0.0 | 0.0 ± 0.0 | 0.0 ± 0.0 | 0.0 ± 0.0 | 0.0 ± 0.0 |
| Isopoda | 0.25 ± 0.25 | 0.0 ± 0.0 | 0.0 ± 0.0 | 0.0 ± 0.0 | 0.0 ± 0.0 | 0.0 ± 0.0 | 0.0 ± 0.0 | 0.0 ± 0.0 | 0.0 ± 0.0 | 0.0 ± 0.0 | 0.0 ± 0.0 | 0.0 ± 0.0 |
| Platyhelminthes | 0.0 ± 0.0 | 0.0 ± 0.0 | 0.0 ± 0.0 | 0.0 ± 0.0 | 0.0 ± 0.0 | 0.0 ± 0.0 | 0.0 ± 0.0 | 0.0 ± 0.0 | 0.0 ± 0.0 | 0.0 ± 0.0 | 0.0 ± 0.0 | 0.0 ± 0.0 |
|  | **Wallace Bay** | | | | | | | | | | | |
|  | **Reference** | | | | | **Restoring** | | | | | | |
| Taxa | May 2022 | May 2023 | June 2022 | June 2023 | July 2022 | July 2023 | May 2022 | May 2023 | June 2022 | June 2023 | July 2022 | July 2023 |
| *Fundulus heteroclitus* | 0.0 ± 0.0 | 0.0 ± 0.0 | 0.0 ± 0.0 | 0.0 ± 0.0 | 1.5 ± 0.87 | 4.25 ± 2.46 | 0.0 ± 0.0 | 0.0 ± 0.0 | 0.0 ± 0.0 | 0.0 ± 0.0 | 0.0 ± 0.0 | 0.0 ± 0.0 |
| Gasterosteidae | 0.0 ± 0.0 | 0.0 ± 0.0 | 0.5 ± 0.5 | 0.0 ± 0.0 | 0.0 ± 0.0 | 0.0 ± 0.0 | 0.0 ± 0.0 | 0.0 ± 0.0 | 0.0 ± 0.0 | 0.0 ± 0.0 | 0.0 ± 0.0 | 0.0 ± 0.0 |
| *Crangon septemspinosa* | 0.0 ± 0.0 | 0.0 ± 0.0 | 0.0 ± 0.0 | 0.0 ± 0.0 | 0.0 ± 0.0 | 0.0 ± 0.0 | 0.25 ± 0.25 | 0.0 ± 0.0 | 0.0 ± 0.0 | 0.0 ± 0.0 | 0.25 ± 0.25 | 0.0 ± 0.0 |
| Mysida | 0.25 ± 0.25 | 0.0 ± 0.0 | 0.0 ± 0.0 | 0.0 ± 0.0 | 0.0 ± 0.0 | 0.0 ± 0.0 | 0.5 ± 0.29 | 0.0 ± 0.0 | 0.0 ± 0.0 | 0.0 ± 0.0 | 0.0 ± 0.0 | 0.0 ± 0.0 |
| Gammaridae | 0.0 ± 0.0 | 0.0 ± 0.0 | 0.0 ± 0.0 | 0.75 ± 0.75 | 3.25 ± 1.18 | 0.25 ± 0.25 | 0.5 ± 0.29 | 0.0 ± 0.0 | 0.0 ± 0.0 | 0.0 ± 0.0 | 1.0 ± 0.71 | 0.0 ± 0.0 |
| *Corophium volutator* | 0.0 ± 0.0 | 0.0 ± 0.0 | 0.0 ± 0.0 | 0.0 ± 0.0 | 0.0 ± 0.0 | 0.0 ± 0.0 | 0.0 ± 0.0 | 0.0 ± 0.0 | 0.0 ± 0.0 | 0.0 ± 0.0 | 0.0 ± 0.0 | 0.0 ± 0.0 |
| Hydrobiidae | 0.0 ± 0.0 | 1.5 ± 0.65 | 0.25 ± 0.25 | 6.0 ± 5.02 | 0.25 ± 0.25 | 30.75 ± 28.15 | 0.5 ± 0.29 | 0.75 ± 0.48 | 0.0 ± 0.0 | 1.0 ± 0.71 | 0.0 ± 0.0 | 3.0 ± 3.0 |
| Corixidae | 0.0 ± 0.0 | 0.0 ± 0.0 | 0.0 ± 0.0 | 0.0 ± 0.0 | 3.75 ± 3.75 | 16.0 ± 14.68 | 0.0 ± 0.0 | 0.0 ± 0.0 | 0.0 ± 0.0 | 0.0 ± 0.0 | 0.0 ± 0.0 | 0.0 ± 0.0 |
| Culicidae larva | 0.0 ± 0.0 | 0.0 ± 0.0 | 0.25 ± 0.25 | 0.0 ± 0.0 | 0.0 ± 0.0 | 0.0 ± 0.0 | 0.0 ± 0.0 | 0.0 ± 0.0 | 0.0 ± 0.0 | 1.0 ± 0.71 | 0.0 ± 0.0 | 0.0 ± 0.0 |
| Other Diptera larva | 0.0 ± 0.0 | 0.0 ± 0.0 | 0.0 ± 0.0 | 0.0 ± 0.0 | 0.0 ± 0.0 | 0.0 ± 0.0 | 0.0 ± 0.0 | 0.0 ± 0.0 | 0.0 ± 0.0 | 0.0 ± 0.0 | 0.0 ± 0.0 | 0.0 ± 0.0 |
| Isopoda | 0.0 ± 0.0 | 0.0 ± 0.0 | 0.0 ± 0.0 | 0.0 ± 0.0 | 0.0 ± 0.0 | 0.0 ± 0.0 | 0.0 ± 0.0 | 0.0 ± 0.0 | 0.0 ± 0.0 | 0.0 ± 0.0 | 0.0 ± 0.0 | 0.0 ± 0.0 |
| Platyhelminthes | 0.0 ± 0.0 | 0.0 ± 0.0 | 0.0 ± 0.0 | 0.0 ± 0.0 | 0.0 ± 0.0 | 0.0 ± 0.0 | 0.0 ± 0.0 | 0.0 ± 0.0 | 0.0 ± 0.0 | 0.25 ± 0.25 | 0.0 ± 0.0 | 0.0 ± 0.0 |

**Supplement 10. Site maps for Aulac and Wallace Bay, Maritime Canada**

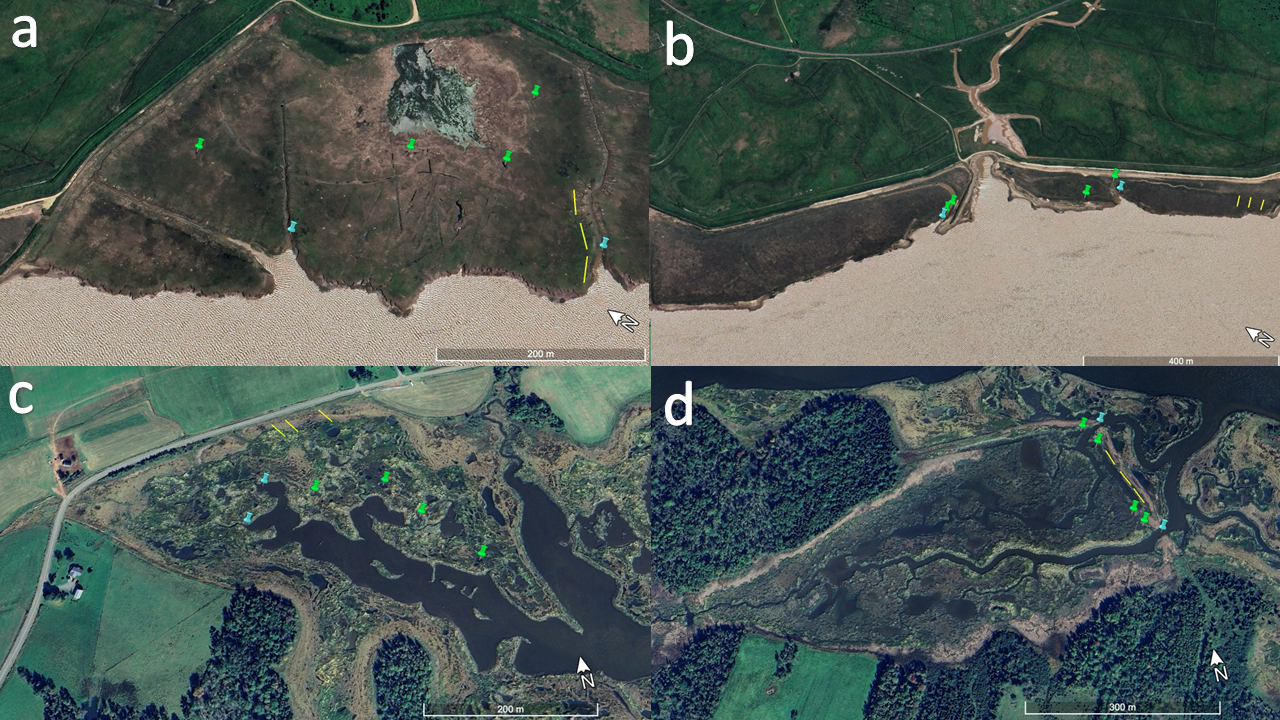

Google Earth™ images of the Reference and Restoring salt marshes in Aulac and Wallace Bay sampled in 2022–2023. The nekton communities in intertidal creeks, on the marsh platform, and in salt pools were sampled during three sampling rounds (May, June, July) at each site and year. Aulac sites: a) Reference (imagery date: 20 June 2024), b) Restoring (imagery date: 20 June 2024). Wallace Bay sites: c) Reference (imagery date: 18 May 2023), d) Restoring (imagery date: 18 May 2023). Blue pins are locations of fyke nets in creeks. Green pins are locations of salt pools or creek pools (exemplified for 2022) sampled with minnow traps and invertebrate activity traps, and for abiotic conditions. Pool locations were somewhat similar between years; small location changes occurred because pools can change shape, depth, and hydrological connectivity between years due in part to winter ice disturbance. Yellow lines represent example seining transects (30 m long) on the marsh platform and the general area where seining occurred at each site. Exact seine haul transects were different between sampling rounds because of the variation in inundation patterns between rounds.
